# Single-cell analysis reveals multi-faceted features of B cell development together with age-associated B cell subpopulations

**DOI:** 10.1101/2024.12.06.627090

**Authors:** Xiujia Yang, Haipei Tang, Chunhong Lan, Weiting He, Sen Chen, Huikun Zeng, Danfeng Liu, Haoyu Wu, Wenjian Wang, Zhenhai Zhang

**Affiliations:** Guangdong Cardiovascular Institute, Guangdong Provincial People’s Hospital, Guangdong Academy of Medical Sciences, Guangzhou 510080, China; Center for Precision Medicine, Medical Research Institute, Guangdong Provincial People’s Hospital (Guangdong Academy of Medical Sciences), Southern Medical University, Guangzhou 510080, China; Guangdong-Hong Kong Joint Laboratory on Immunological and Genetic Kidney Diseases, Guangdong Provincial People’s Hospital (Guangdong Academy of Medical Sciences), Southern Medical University, Guangzhou 510080, China; Department of Bioinformatics, School of Basic Medical Sciences, Southern Medical University, Guangzhou 510515, China; Division of Nephrology, Guangdong Provincial People’s Hospital (Guangdong Academy of Medical Sciences), Southern Medical University, Guangzhou, 510080, China; Key Laboratory of Mental Health of the Ministry of Education, Guangdong-Hong Kong-Macao Greater Bay Area Center for Brain Science and Brain-Inspired Intelligence, Southern Medical University, Guangzhou 510515, China

**Keywords:** B cell development, B cell receptor, age-associated B cell, single cell, high-throughput sequencing

## Abstract

The development and maturation of B lymphocytes involve intricate orchestrated processes, where dedicated gene regulations (GR) take place within specific microenvironments shaped by both extracellular matrix and neighboring cells. Despite extensive investigations aimed at deepening our comprehension of these mechanisms, there remains a dearth of high-dimensional and integrated analysis concerning B cell heterogeneity, gene regulation, and external factors implicated in B cell development. In this study, we scrutinized single-cell transcriptomic data and B cell receptor (BCR) sequencing data obtained from B cells and their surrounding counterparts in the bone marrow, tonsil, and peripheral blood. A full picture of the GR dynamics, the heterogeneity of conventional B cells and cell-cell interactions (CCIs) along B cell development axis was depicted. We found immature B cells represent the most quiescent stage characterized by the least number of expressed genes and low RNA velocity. The homeostatic proliferation and activation of naïve B cells is niche-confined and individualized, respectively. Two development models for memory B cell subpopulations seem not mutually exclusive and warrant in-depth investigation. Moreover, CCI analysis reveals a pivotal role of myeloid cells and two dominant and stage-dependent CCI categories, TNF and adhesion signaling, in B cell development. Besides, we unexpectedly identified two age-associated B cell subpopulations that respectively express S100A8/S100A9 and C1q and experimentally confirmed the secretion of S100A8/A9 from human B cells *in vitro*, suggesting a senescence-associated secretion phenotype. Our integrated analysis provides valuable insights into GR dynamics, the evolution of B cells, and potential intercellular communication networks involved in B cell development and revealed novel phenotypes of age-associated B cell aberrance. This study serves as a valuable resource for in-depth exploration of the intricacies of B cell biology.

---

B lymphocytes (or B cells) descend from bone marrow (BM) hematopoietic stem cells (HSC) and mediate humoral immunity by secreting antibodies. Early studies based on a selected set of surface and intracellular markers and specific gene knockout mice have greatly added to the knowledge repertoire of B cells. They include but are not limited to, i) B cell development stages as defined by the status of VDJ gene rearrangement of heavy and light chains, ii) central and peripheral tolerance that ensure removal or anergy of self-reactive B cells, iii) well-organized BM niches required for B cell lymphogenesis, and iv) the germinal center (GC) reaction model that contributes to the genesis of affinity-matured B cells with the facilitation of follicular helper T cells (T_fh_) and dendritic cells (FDC).

As pointed out by both Lee *et al*.^1^ and King *et al*.^2^, previous low-dimensional and low-throughput methodologies fall short of full demarcation of distinct B cell subsets and deconvolution of intrinsically connected biological processes. The advent of single-cell technologies now has bridged this gap and led to breakthroughs in understanding B cell heterogeneity, lineage trajectories, B cell receptor (BCR) repertoires, and cell-cell communication networks^3^. For example, Lee *et al*. coupled single-cell RNA sequencing (scRNA-seq) and CITE-Seq (Cellular Indexing of Transcriptomes and Epitopes by Sequencing) and revealed two distinct proliferative phases (i.e. pre-BCR-dependent and pre-BCR-independent) in the pre-B cell expansion stage in mice^1^. Furthermore, by correlating with the canonical markers, they identified YBX3 and EBF1 as the defining features for the two phases, respectively. Another investigation, performed by King *et al*., focused on B cell maturation pathway delineation by integrating single-cell transcriptomic and antibody repertoire data^2^. The collaborators reported a pre-GC state programmed to undergo class switch and the antibody-class-dictated B cell fate decision. More than these, single-cell techniques also prove to be powerful in resolving B cell subset heterogeneity^4,5^, revealing new niches where B cell develops^6^, and shedding light on B cell compartment aberrance in B-cell-related carcinoma^7^.

Given these progresses, there still lacks a comprehensive high-dimensional single-cell analysis that covers a continuum of B cell stages ranging from early progenitor to mature naïve compartment and then to antigen-experienced memory compartment or terminally differentiated PC. Neither the heterogeneity of certain conventional B cell subsets nor the environmental factors contributing to the development of B cells in humans were carefully examined.

Inspired by the success of aforementioned B cell studies, we analyzed the single-cell transcriptomic and BCR sequencing data of B cells from BM, peripheral blood (PBL) and tonsil GC procured from either in-house experiments or external studies. We provided a gene regulation landscape for 18 well-defined B cell subpopulations covering a continuum of developmental stages. By dissecting the heterogeneity of conventional B cell subsets (i.e. immature, naïve and memory B cells), we revealed interesting properties related to several biological processes including B cell senescence, homeostatic proliferation (HP) and memory B cell differentiation. Finally, we constructed tissue-specific interaction networks between B and non-B cells and demonstrated the interaction attributes both quantitatively and qualitatively along the B cell developmental axis. Our work resolved the gene regulation dynamics, the heterogeneity of conventional B cells, and the underlying role of surrounding non-B cell types and signaling as B cell develops and matures, which will serve as a valuable resource for future studies.

## Results

### Study design and the integrated single-cell dataset

This study aimed to identify a full spectrum of B cell subpopulations along the development axis, depict the underlying gene regulations and their relationships, and subsequently delineate cell-cell communications among them and other coexisting cell types. The scRNA-seq data of the enriched early B cell groups^7^, the tonsil GC B cells^2^, and peripheral blood mononuclear cells (PBMCs)^8^ were downloaded via previous works (Table S1). To acquire thorough B cell groups and their corresponding environmental cell types, we obtained BM aspirates and PBL from three individuals free from hematological disorders (FFHD). For each sample, we conducted scRNA-seq for both total mononuclear cells (BM only) and enriched B cells (CD3^-^CD41a^-^CD43^-^CD235a^-^) (see Materials and Methods). In addition, we also performed scVDJ-seq for the B cells and obtained bulk antibody repertoires for the PBL and BM samples via DUMPArts developed earlier in the lab^9^. Subsequently, we integrated our data with datasets from previous studies and thus secured a wealth of resources for B cell subpopulation identification and the successive analyses of gene regulation and cell-cell interactions (CCIs) crucial for B cell development and maturation (Fig. 1A).

**Figure 1.**
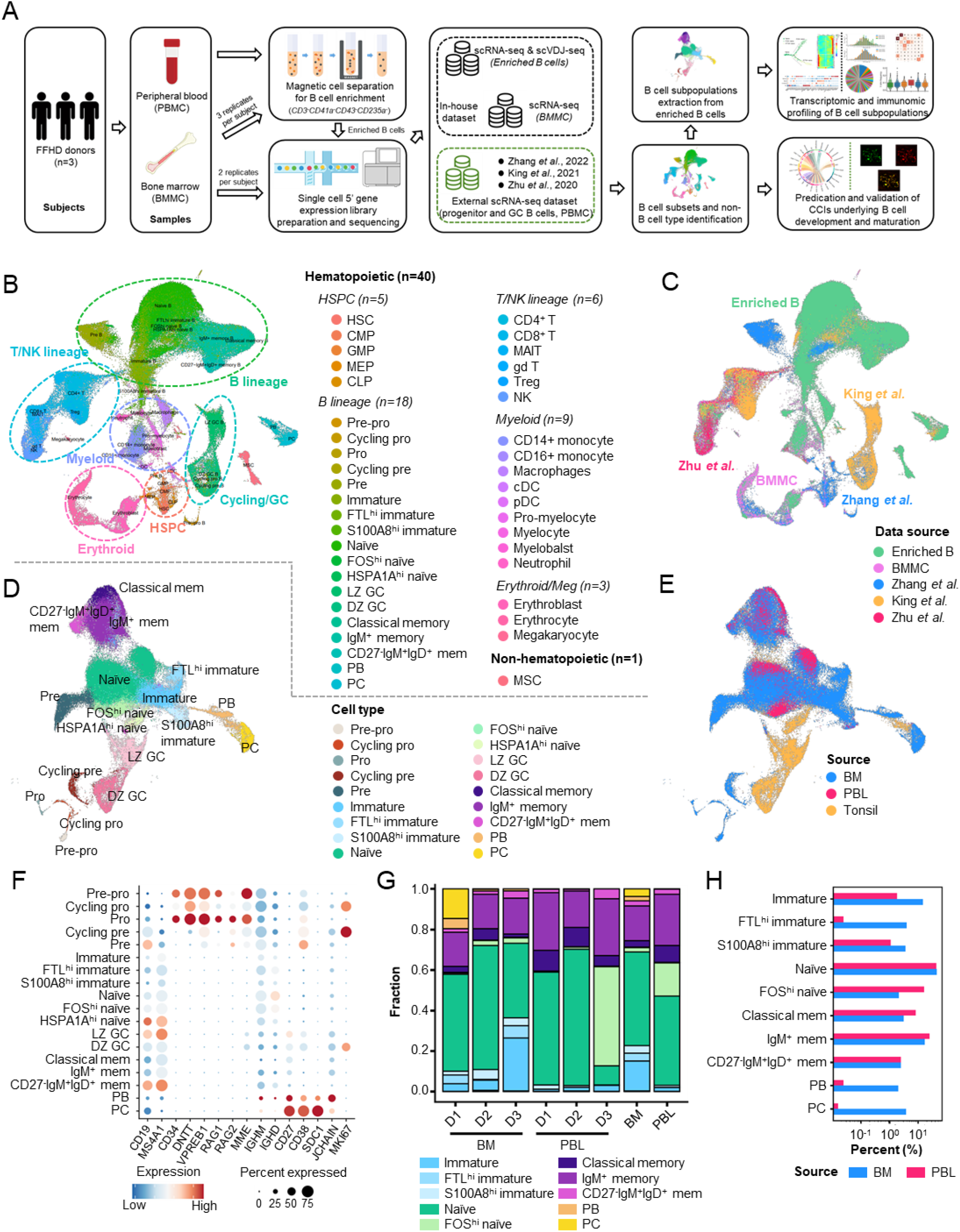
Study design and the landscape of cell populations in the integrated single-cell dataset. **(A)** Overview of the study design. **(B-C)** UMAP of 42 cell populations (B) and data sources (C) in the comprehensive integrated data comprising of in-house enriched B cells and unsorted BMMCs and external data (enriched early B progenitors, GC B cells and unsorted PBMC, see Table S1 for details). **(D-E)** UMAP of B cell subpopulations (D) and sample sources (E) in the enriched B cell dataset. **(F)** Expression of marker genes across 18 B cell subpopulations. **(G)** Tissue-specific composition of B cells across donors and as a whole. **(H)** Frequency comparison of B cell subpopulations between BM and PBL (only 10 subpopulations that represent at least 1% of the enriched B cells in either BM or PBL were shown).

This study involved 199,004 cells, of which 129,832 (65.2%) were newly generated in our lab and 69,172 (34.8%) were from external datasets. Quality-wise, the enriched B cells and unsorted BM mononuclear cells (BMMCs) captured medians of 1,420 and 1,414 genes, respectively. The median captured transcripts were 3,662 and 4,134. Table S2 and Fig. S1 provide detailed quality metrics for scRNA-seq and scVDJ-seq samples. Data integration was performed using Seurat’s ‘fast integration’ method to eliminate the batch effect. We then employed a two-round clustering to identify cell populations unbiasedly. The first round of clustering distinguished various cell lineages (Fig. S2A) and the second round revealed specific cell populations within a typical lineage (Fig. S2 B-E) (see Materials and Methods). These well-classified cell populations thus served as the starting point for the downstream analysis after retro-integrated to the original dataset.

As a result, we identified forty-two cell populations, including 5 hematopoietic stem and progenitor cell (HSPC) populations, 18 B lineage populations, 6 T/NK lineage populations, 9 myeloid populations, 3 erythroid/megakaryocyte populations, and 1 non-hematopoietic population (Fig. 1B). The data source distribution also reflects a reasonable cell identity assignment (Fig. 1C). Importantly, our result reproduced the frequencies of cell population in CD45^+^ BMMC^10^ and unsorted PBMC^8^ indicating the analyses approach was reliable and solid (Fig. S3). Table S3 provides detailed statistics, including cell population frequencies in the integrated data. Therefore, this well-annotated integrated dataset provided a solid foundation for investigating B cell heterogeneity and CCIs underlying B cell development and maturation.

Given that B lineage is the primary focus of this study, we carefully examined the resulting B cell clusters. Combining the collected canonical marker genes and computed cluster-specific genes, we identified 18 B cell subpopulations, covering the entire spectrum of B cell development and maturation, from early B cell progenitors to terminally differentiated plasma cells (PC) (Fig. 1 D-F and Fig. S4). The data and tissue source composition for each B cell subpopulation were provided as Fig. S5. Notably, aside the 14 well-known B cell subpopulations, four minor subpopulations with specific highly expressed genes were identified for B cells of immature and naïve phenotypes (i.e. FTL^hi^ and S100A8^hi^ for immature B cells, FOS^hi^ and HSPA1A^hi^ for naïve B cells) (Fig. 1F), which will be examined in depth later.

We then quantified these B cell subpopulations for the in-house enriched B cell compartment. Overall, B lineage cells represented 85.6% and 92.9% of enriched BM and PBL, respectively. The non-B compartment was dominated by myeloid cells (for both BM and PBL) and mesenchymal stem cells (for BM only) (Fig. S6). B cell subpopulation frequencies vary among donors (Fig. 1G and Table S4). For instance, D1 and D3 had a significantly higher proportion of BM PC (and plasmablast (PB)) and immature B cells, respectively, whereas D3 had a higher proportion of PBL FOS^hi^ naïve B cells. Despite the donor variation, the proportions of major B cell subsets (immature, naïve, memory B cells, and PC) in mixed samples recapitulated the estimation previously reported^11,12^ (Fig. 1H). Nonetheless, it is worth noting that we only observed a minimal fraction (<5‰) of early-stage B cells (i.e., pre-pro, (cycling) pro, and (cycling) pre-B cells) in the enriched BM B cell compartment. In addition to their low frequencies in BM B lineage cells^11^, the underrepresentation of these early-stage subpopulations can also be attributed to the B cell enrichment strategy^13^ (see Materials and Methods).

### A transcriptomic overview of the 18 identified B cell subpopulations

At the transcriptomic level, the number of captured genes for these subpopulations fluctuated significantly along the developmental axis (Fig. 2A). For example, the 3 proliferating groups, namely cycling pro, cycling pre and DZ GC, exhibited the maximum number of genes (medians of 3,664, 3,296 and 2268, respectively). Apart from these proliferating subpopulations, the gene numbers showed a V-shape as B cell matures, where the minimum number occurred in immature and PB subpopulations (577, 561 and 548 for immature, S100A8^hi^ immature and PB, respectively). The decreased number of capture genes in immature B cells can probably account for their low metabolism rate, as reported in a previous study^14^. It is worth noting that we confirmed these observations in independent projects to avoid potential bias caused by the batch effect (Fig. S7).

**Figure 2.**
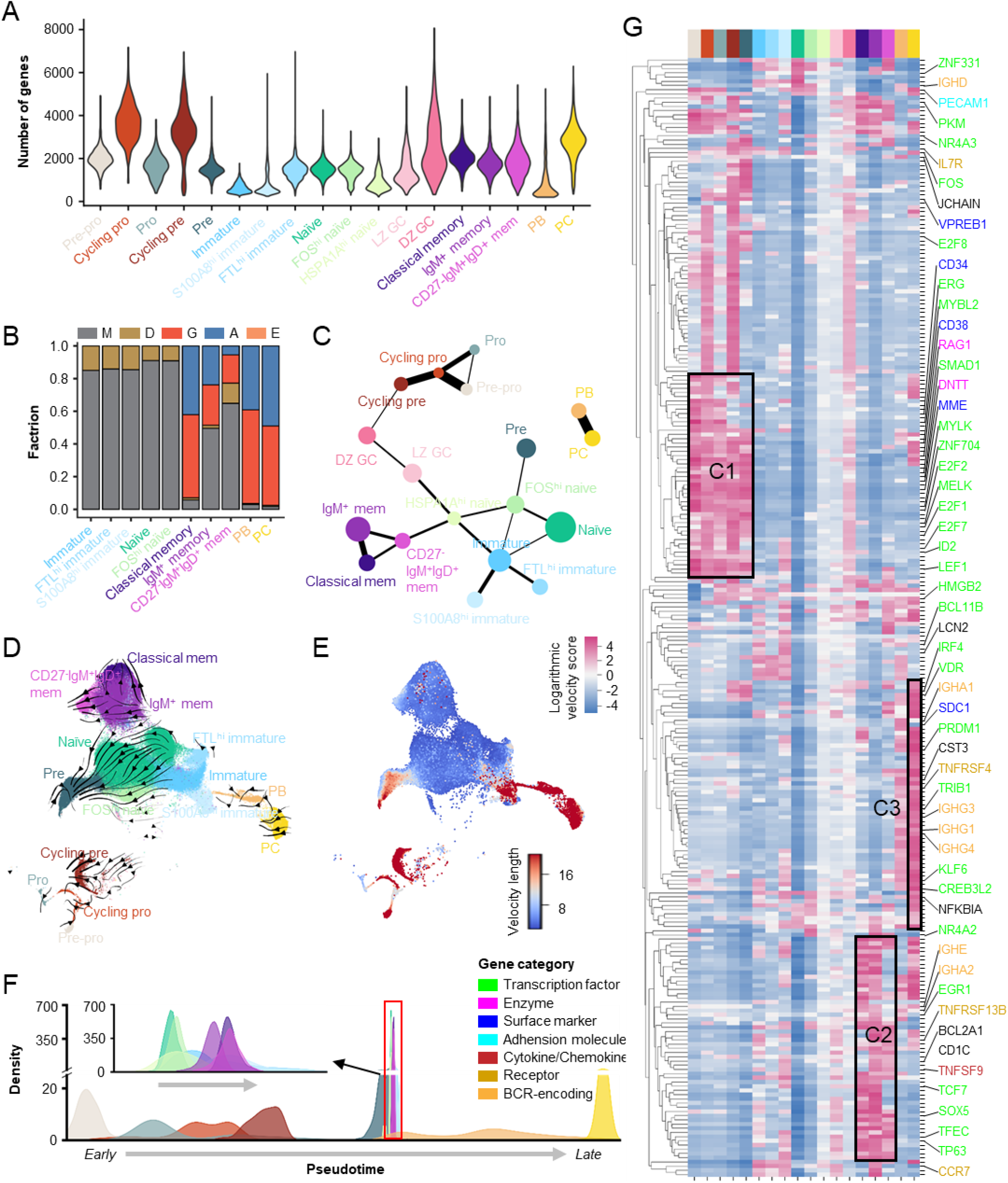
Gene regulation dynamics of B cell development revealed by scRNA-seq analysis. **(A)** Distribution of number of genes captured by scRNA-seq across the 18 identified B cell subpopulations. **(B)** BCR isotype composition for the 10 B cell subpopulations captured in our in-house scVDJ-seq data. **(C)** Connectivity of B cell subpopulations as shown by the partition-based graph abstraction (PAGA). The connectivity threshold was set as 0.1 for a concise representation of these B cell subpopulations. The circle size is proportional to the subpopulation size. Line width denotes the strength of connectivity. **(D)** Streamline-based visualization of RNA velocities of the 18 B cell subpopulations. **(E)** The differentiation speed of the 18 identified B cell subpopulations as given by the length of the velocity vector. **(F)** Subpopulation-specific pseudotime distribution inferred based on RNA velocity. The inset in the top left demonstrats a zoom-in pseudotime window occupied by immature, naïve and memory B cell subpopulations. **(G)** Profile of the top-active genes (measured by velocity score) for the 18 B cell subpopulations. All transcription factors and a selected list of representative genes were marked at the side. The gene category is color-coded.

For scVDJ-seq, the capture efficiency of BCRs varied among subpopulations (Table S5). Only less than 20% of primary and S100A8^hi^ immature subpopulations were captured with BCRs. In contrast, the BCR capture rates were 89.4% and 86.6% for PB and PC, respectively. IgM was the dominant isotype in all 3 immature (85.0%, 85.9% and 85.4%), 2 naïve (91.0% and 90.9%) and 2 memory subpopulations (49.5% and 64.9%) (Fig. 2B). Notably, IgM was not the exclusive isotype in the 3 immature subpopulations. IgD was also present in around 15% of the cells in all three immature subpopulations. In contrast, class-switched isotypes, mostly IgG and IgA, dominate the classical memory, PB and PC subpopulations.

Extensive mouse-based experiments have led to a well-established B cell development paradigm^15^. Although it has no debate on the time order of key stages in B cell development, our partition-based graph abstraction (PAGA) analysis revealed a seemingly non-successive transcriptomic stage transition in the pre-immature-naïve development axis. We found pre B cells were more confidently connected to FOS^hi^ naïve B cells rather than the intermediate immature stage (Fig. 2C). Despite this counterintuitive observation, the PAGA graph also reflected tight connections between immature and naïve subpopulations and between three cycling subpopulations (i.e. cycling pro, cycling pre and DZ GC) and the distinctness of PB and PC.

Then RNA velocity analysis was performed to investigate the gene expression kinetics of B cell development (HSPA1A^hi^ naïve and GC B cells were not included due to the unavailability of raw sequencing data required in this analysis). The streamlines projected on the UMAP (uniform manifold approximation and projection) showed B cell stage transitions in 4 compartments (part 1: pre-pro -> cycling pro -> pro -> cycling pre; part 2: immature -> naïve; part 3: IgM^+^ memory -> classical memory, IgM^+^ memory -> CD27^-^IgM^+^IgD^+^; part 4: PB->PC), separately delineating the local relationship between subpopulations (Fig. 2D). We subsequently investigated the differentiation speed of these B cell subpopulations and found a contrastingly higher RNA velocity for both the earliest stage B cells (from pre-pro to pre to S100A8^hi^ immature) and terminal differentiated PB and PC compared to the rest subpopulations (Fig. 2E). The lower differentiation speed for immature, naïve and memory subpopulations was also reflected by clearly narrow pseudotime windows they occupied, reflecting a relatively quiescent state (Fig. 2F). Lastly, we profiled the top-active genes (measured by velocity score) across all B cell subpopulations (Fig. 2G). Generally, three gene clusters stood out for their restricted activity in early B cells (cluster 1 or C1), memory B cells (cluster 2 or C2) and PC (cluster 3 or C3), respectively. On the contrary, immature and naïve subpopulations were characterized by a sparsely distributed and low-velocity gene expression pattern, suggesting a non-specific regulation machinery and a comparatively resting status.

A selected list of genes, including transcription factors (TF) and some canonical molecules (e.g. surface markers/receptors, recombination enzymes, BCR-encoded genes, and adhesion molecules) was marked out (Fig. 2G and Fig. S8). Specifically, the TFs, ERG, LEF1, SMAD1, MYLK, and ZNF704, were most active in the earliest stage of B cells (i.e. pre-pro B cells). In contrast, the E2F family TFs and MELK peaked at a later stage (i.e. pre B cells). ZNF331 was the sole TF found active in the naïve subpopulation. PKM, SOX5, TP63 and TFEC were active in memory B cells, and TRIB1, PRDM1, CREB3L2 and KLF6 were active in PC. Notably, MYBL2, IRF4 and VDR were active in both early B cells and PC.

### Two atypical immature B cell subpopulations are characterized by senescence-associated secretory phenotype

Since we identified two formerly unappreciated immature B cell subpopulations, it is of great interest and significance to map them on the developmental trajectory and decipher their functional status. Applying Monocle2 to the trajectory inference analysis, we found the two minor immature subpopulations distributed in different branches, suggesting a distinct development trajectory (Fig. 3A-B). This was consistent with the developmental topologies revealed by two additional methodologies, namely PAGA (Fig. 2C) and RNA velocity analysis (Fig. 3C). Notably, FTL^hi^ and S100A8^hi^ immature B cell subpopulations expressed C1q and S100A8/S100A9 molecules, respectively (Fig. S4). The two molecules are typically expressed by myeloid cells, such as dendritic cells, macrophages, monocytes and neutrophils, rather than by B lineage cells^16,17^. This unusual expression pattern further implied that the two subpopulations were out of the normal B cell differentiation pathway and represented two atypical B cell subpopulations. To double confirm it, we scrutinized 6 additional published human BM scRNA-seq datasets and found the two subpopulations were not common in these external BM samples^10,18–22^. Briefly, only S100A8^hi^ immature B cell subpopulation was recovered in two of the six datasets, while none of these datasets contained the FTL^hi^ immature B cell subpopulation (see the last section for details).

**Figure 3.**
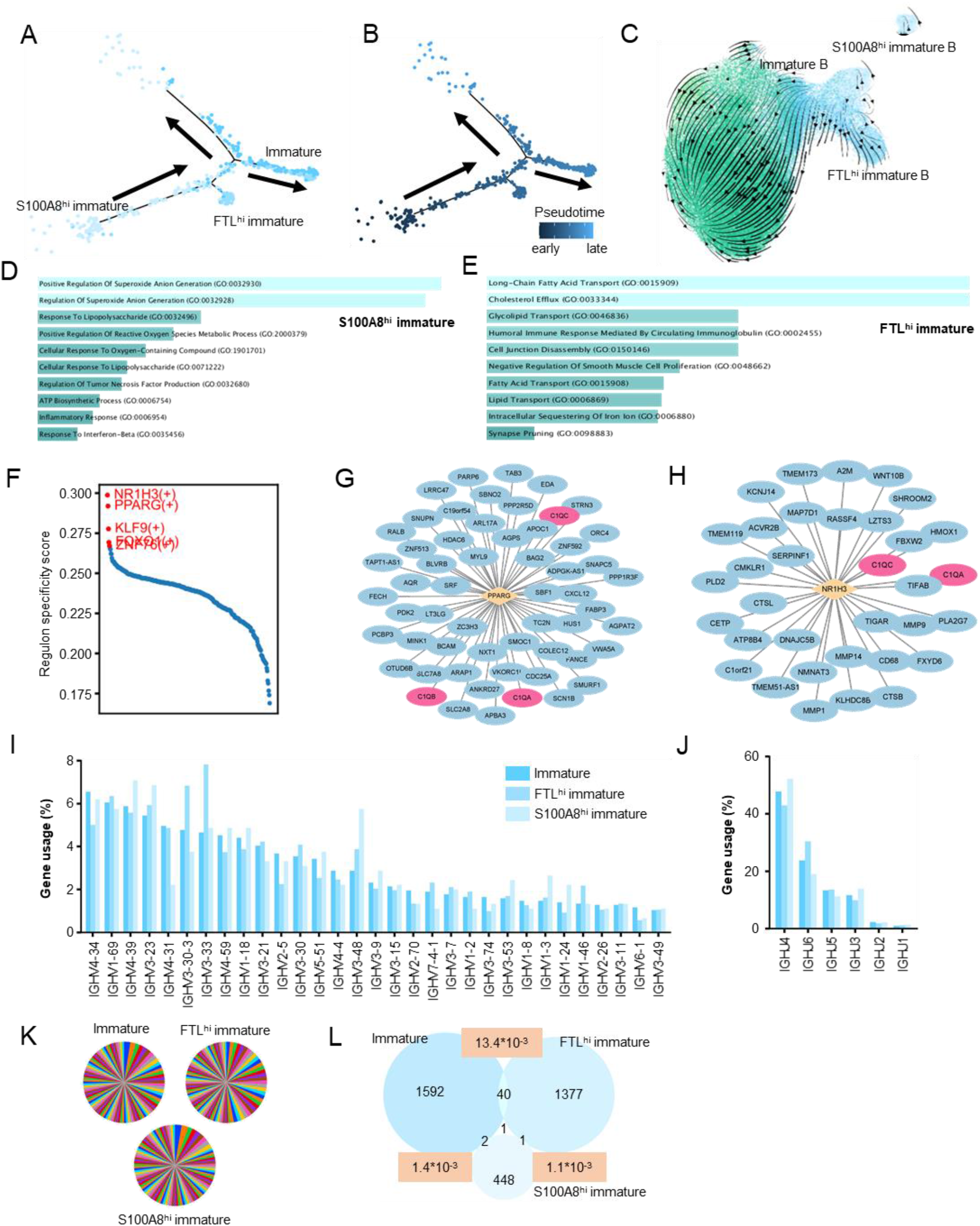
Two minor immature subpopulations are characterized by atypical differentiation and senescence-associated secretory phenotype. (A-B) Transcriptomic-based inference of developmental trajectory for 3 immature B cell subpopulations with monocle2. **(C)** Streamline-based visualization of RNA velocities of the immature and naïve B cell subpopulations. The color scheme is the same as that in Fig . 1D. **(D-E)** The top 10 enriched biological processes as sorted by significance score (p-value) for S100A8^hi^ (D) and FTL^hi^ (E) immature B cell subpopulations. A longer and lighter bar indicates a more significant sore. **(F)** The top 5 regulons specific for the FTL^hi^ immature B cell subpopulation as inferred by SCENIC workflow. **(G-H)** Regulation networks reflecting the two regulons (PPARG(+) (G) and NR1H3(+) (H)) predicted by SCENIC workflow. Transcription factors and their corresponding target genes are surrounded by orange diamonds and blue ellipses, respectively. C1q component genes, namely C1QA, C1QB, and C1QA, are highlighted in pink. **(I-J)** BCR heavy chain variable (I) and joining (J) gene usage as determined by scVDJ-seq data across 3 immature B cell subpopulations. For variable genes, only those with a usage frequency of at least 1% were shown. **(K)** Pie charts showing the relative frequencies of top 100 clonotypes for 3 immature B cell subpopulations. **(L)** Venn diagram showing clonotype sharing between 3 immature B cell subpopulations. The numbers on the diagram indicate the number of clonotypes in each compartment. The numbers with a pink background indicate the clonal similarity as measured by the Jaccard similarity index.

Subsequently, we performed GO enrichment analysis to interrogate the functional state of the two minor immature subpopulations. The result showed that the S100A8^hi^ immature B cell subpopulation was enriched for biological pathways like “*Regulation Of Superoxide Anion Generation*”, “*Positive Regulation Of Reactive Oxygen Species Metabolic Process*”, and “*Inflammatory Response*” (Fig. 3D), which seemed to correlate with the senescence-induced inflammatory process^23^. As for FTL^hi^ immature B cell subpopulation, lipid metabolism-related pathways including “*Long-Chain Fatty Acid Transport*”, “*Cholesterol Efflux*” and “*Glycolipid Transport*” were enriched (Fig. 3E). Considering the high expression of C1q components (i.e. C1QA, C1QB and C1QC) in this subpopulation, these lipid metabolism pathways may facilitate the exportation of this multi-faced effector component of the innate immune response. It has been reported that serum C1q levels in plasma increase with aging^24^. The atypical expression of C1q by B lymphocyte probably represents another source of these inflammatory molecules during aging.

We then investigated the regulatory factors driving the atypical expression C1q with SCENIC workflow. The result revealed that NR1H3 and PPARG were the two most specific TFs in FTL^hi^ immature B cell subpopulation among all B cell subpopulations (Fig. 3F and Fig. S9). Moreover, three C1q component genes were also among the gene lists of their associated regulons (Fig. 3G- H).

We further characterized their expressed BCR repertoire with scVDJ-seq data. Although variations of heavy chain variable gene usage were observed among three immature B cell subpopulations, they could not be reproduced in individual donors (Fig. 3I and Fig. S10A), which is also the case for heavy chain CDR3 length distribution (Fig. S10B). In contrast, the heavy chain joining gene, light chain variable and joining genes, and light chain CDR3 demonstrated comparable usage frequencies or length distributions (Fig. 3J and Fig. S10C-E), which suggested a BCR-independent induction pathway for the two atypical B cell subpopulations. The clonality analysis indicated that all three subpopulations featured an evenly distributed clonal size without dominant clonotypes, agreeing with their immature stage and, thus, innocent phenotype (Fig. 3K). However, we indeed found a clonal relationship between these subpopulations, indicating a modest homeostasis-like proliferation of immature B cells. Notably, we observed a ten-fold higher clonal similarity (as measured by the Jaccard similarity index) between primary and FTL^hi^ subpopulations than between S100A8^hi^ and either of the other two subpopulations, suggesting a closer relationship between the two subpopulations (Fig. 3L). However, it should be noted that the shared clonotypes (n=40) between primary and FTL^hi^ immature B cell subpopulations all come from a single donor (D3). Therefore, whether this observation can be extrapolated remains to be validated.

### Naïve B cells show a confined HP and an individualized activation pattern

Naïve B cells are generally deemed as a homogenous B cell subset. However, King *et al.* described two additional naïve-like subsets coined as ‘activated’ and ‘preGC’ in their GC response study^2^. With this in mind, we compared the two naïve subpopulations (i.e. FOS^hi^ and HSPA1A^hi^ naïve B cells) in this work with the two they described. A high consistency in the top 10 highly expressed genes was observed between the FOS^hi^ naïve B cells and the ‘activated’ subset (Fig. S11), pointing to the activated phenotype of the FOS^hi^ subpopulation. The activated state of the FOS^hi^ subpopulation was also implied by both PAGA analysis (Fig. 2C) and the development trajectory constructed by Monocle2, where a primary naïve -> FOS^hi^ -> HSPA1A^hi^ development axis was revealed (Fig. 4A-B). Notably, both primary and FOS^hi^ naïve B cell subpopulations were found in all three sources (BM, PBL and tonsil), whereas nearly all cells in the HSPA1A^hi^ subpopulation come from the tonsil (Fig. S5B). This particular sample source for the HSPA1A^hi^ subpopulation, together with the niche it occupied in the development trajectory, led us to assume that it represents the ‘preGC’ subset termed by King *et al*. However, we could not observe additional evidence to support this assumption. The marker genes expressed in the ‘preGC’ subset were not pronounced in the HSPA1A^hi^ subpopulation (Fig. S12). Moreover, tracing back to King’s dataset showed that these cells primarily come from the ‘naïve’ subset.

**Figure 4.**
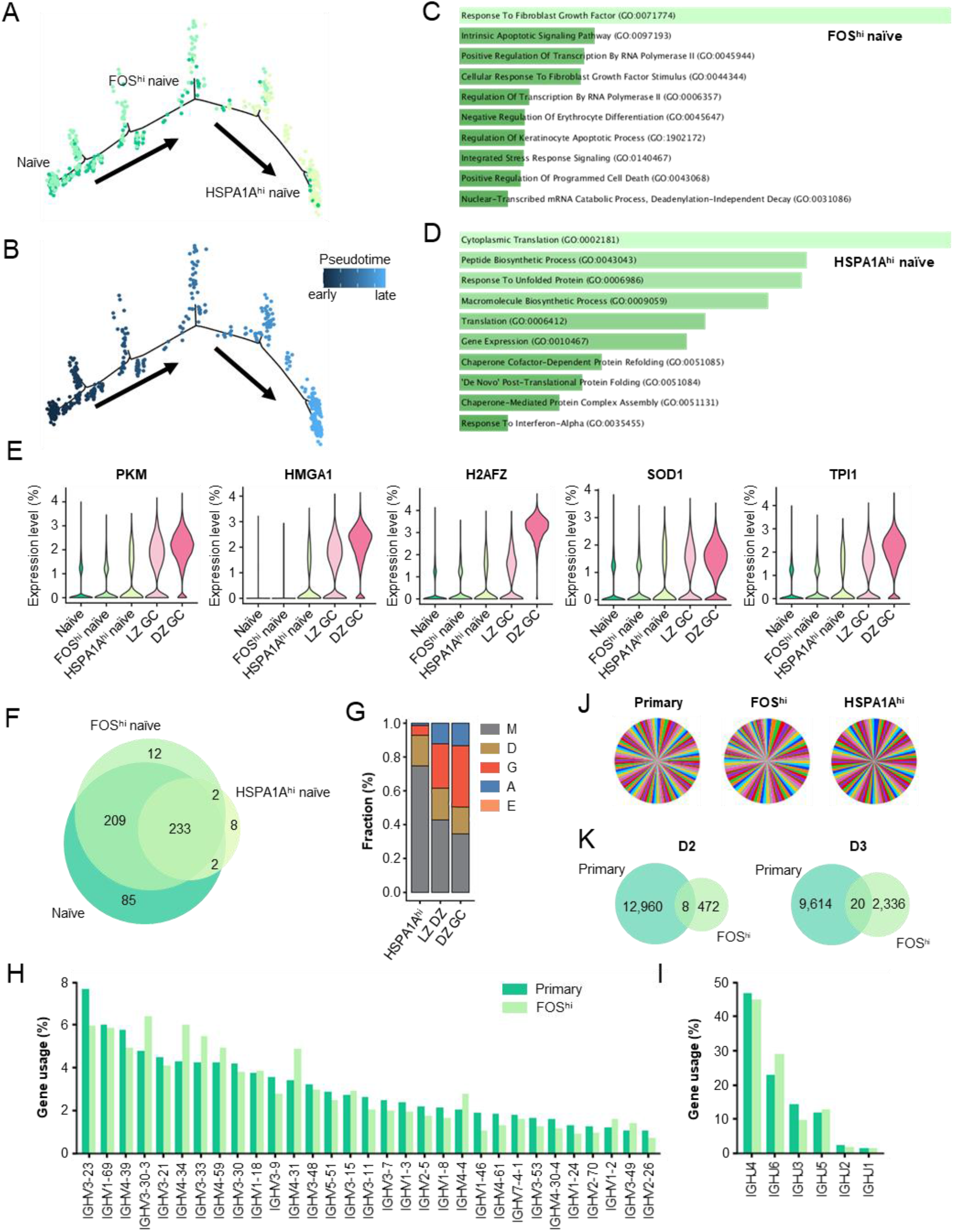
Naïve B cell is characterized by a confined HP and an individualized activation pattern. (A-B) Transcriptomic-based inference of developmental trajectory for 3 naïve B cell subpopulations with monocle2. **(C-D)** The top 10 enriched biological processes as sorted by significance score (p-value) for FOS^hi^ (C) and HSPA1A^hi^ (D) naïve B cell subpopulations. A longer and lighter bar indicates a more significant score. **(E)** Gene expression of 5 TFs highly expressed in the HSPA1A^hi^ subpopulation in 3 naïve subpopulations and the succeeding GC B cell subpopulations. **(F)** Overlapping of commonly expressed genes (CEGs) of three naïve subpopulations. **(G)** BCR isotype composition for HSPA1^hi^, LZ GC and DZ GC B cells (based on King’s dataset). **(H-I)** BCR heavy chain variable (H) and joining (I) gene usage for the primary and FOS^hi^ naïve B cell subpopulations. For variable genes, only those with a usage frequency of at least 1% were shown. **(J)** Pie charts showing the relative frequencies of the top 100 clonotypes for 3 naïve B cell subpopulations. **(K)** Venn diagram showing clonotype sharing between primary and FOS^hi^ naïve B cell subpopulations for two donors (D2 and D3). Numbers on the diagram indicate the number of clonotypes in each compartment.

To gain insight into the two particular naïve-like subsets, we performed GO enrichment analysis to investigate their underlying biological processes. In concordance with the activated phenotype, the FOS^hi^ subpopulation was enriched with the term “positive regulation of transcription by RNA polymerase II” (Fig. 4C). In contrast, the HSPA1A^hi^ subpopulation was enriched with multiple terms related to translations as well as the term “response to unfolded protein” (Fig. 4D). Remarkably, the latter is indeed constituted by genes encoding heat shock proteins (HSPs), as the markers of this subpopulation. It has been reported that HSP90 is upregulated following CD3/CD28 stimulation, suggesting that HSP expression might be regulated via TCR^25^. However, it is unknown whether this elevated HSP expression we observed can be attributed to the BCR-dependent activation. We also identified 5 TFs (i.e. PKM, HMGA1, H2AFZ, SOD1 and TPI1) with elevated expressions as B cell matures, which possibly are implicated in the emergence of this subpopulation (Fig. 4E). Furthermore, we compared the commonly expressed genes (CEGs, expressed in at least 50% cells within a subpopulation) between three naïve subpopulations. It showed that the HSPA1A^hi^ subpopulation expressed the minimum set of CEGs, which were included mainly by the two other subpopulations (Fig. 4F). The biological significance of this simplified gene expression machinery should be further elucidated.

Then we characterized the expressed BCRs for these naïve subpopulations (King’s BCR data was used for the inclusion of the HSPA1A^hi^ subpopulation for comparison). Notably, a higher percentage of the HSPA1A^hi^ B cells were found with BCRs of switched isotypes (IgG and IgA) compared with the other two subpopulations (Fig. 4G vs Fig. 2B), suggesting the emergence of class switch recombination. Although variations of gene usage were observed between the primary and FOS^hi^ naïve subpopulations (HSPA1A^hi^ has a different donor source and is thus not feasible for gene usage comparison) (Fig. 4H-I and Fig. S13A-B), they could not be reproduced across individuals (Fig. S13C-D), indicating no clear gene usage preference for BCRs primed for activation in physiological condition. We also did not observe tissue-specific gene usage pattern for the two subpopulations (Fig. S13E-F). Lastly, clonality analysis revealed that all three subpopulations had no clonal expansions (Fig. 4J). However, we observed a clonal relationship between primary and FOS^hi^ naïve subpopulation for both donors (Fig. 4K). Remarkably, this clonal relationship between subpopulations was only observed in the same source, reflecting a local and confined B cell HP^26^.

### Evidence supporting two distinct development models for memory B cells coexists

The heterogeneity of memory B cells has been appreciated for decades^27^. A combination of CD27 and several Ig classes (e.g. IgM, IgD, IgG and IgA) is the most widely-used surface marker set in research examining memory B cell subsets. With these surface markers, a dozen memory subsets were isolated artificially, among which some may overlap with each other depending on the selection of marker sets^28^. Thus, a limited and predefined set of markers does not guarantee a biologically meaningful category of memory subsets featured by distinct origin or function. Benefiting from the high dimensional transcriptome data, we found that there existed two major memory B cell branches in three representative sources (BM, PBL and tonsil). To be exact, classical (referred to as C) and IgM+ (M) memory B cells represented a branch, while CD27-IgM+IgD+ (MD) represented another branch. In the constructed development trajectory, MD was found as an independent population that contained few cells in a transitional state towards the other two subpopulations, suggesting a gap between the two major branches (Fig. 5A-B). The distinctness of the two branches was also demonstrated by the 3D UMAP plot (Supp. Data 1, umap.gif) as well as by a lower connectivity between MD and C/M than between C and M (Fig. 2C). These observations coincided with the two well-recognized memory generation pathways, namely T cell-dependent (TD, for C and M) and -independent (TI, for MD) pathways^29,30^. Both M and C subpopulations have a TD activation history but differ from each other in that the former comes from a primary GC response while the latter comes from a secondary or consecutive GC response^28,31^. A longer maturation history for C subpopulation was reflected by a delayed emergence in the development axis compared to M subpopulation (Fig. 2F).

**Figure 5.**
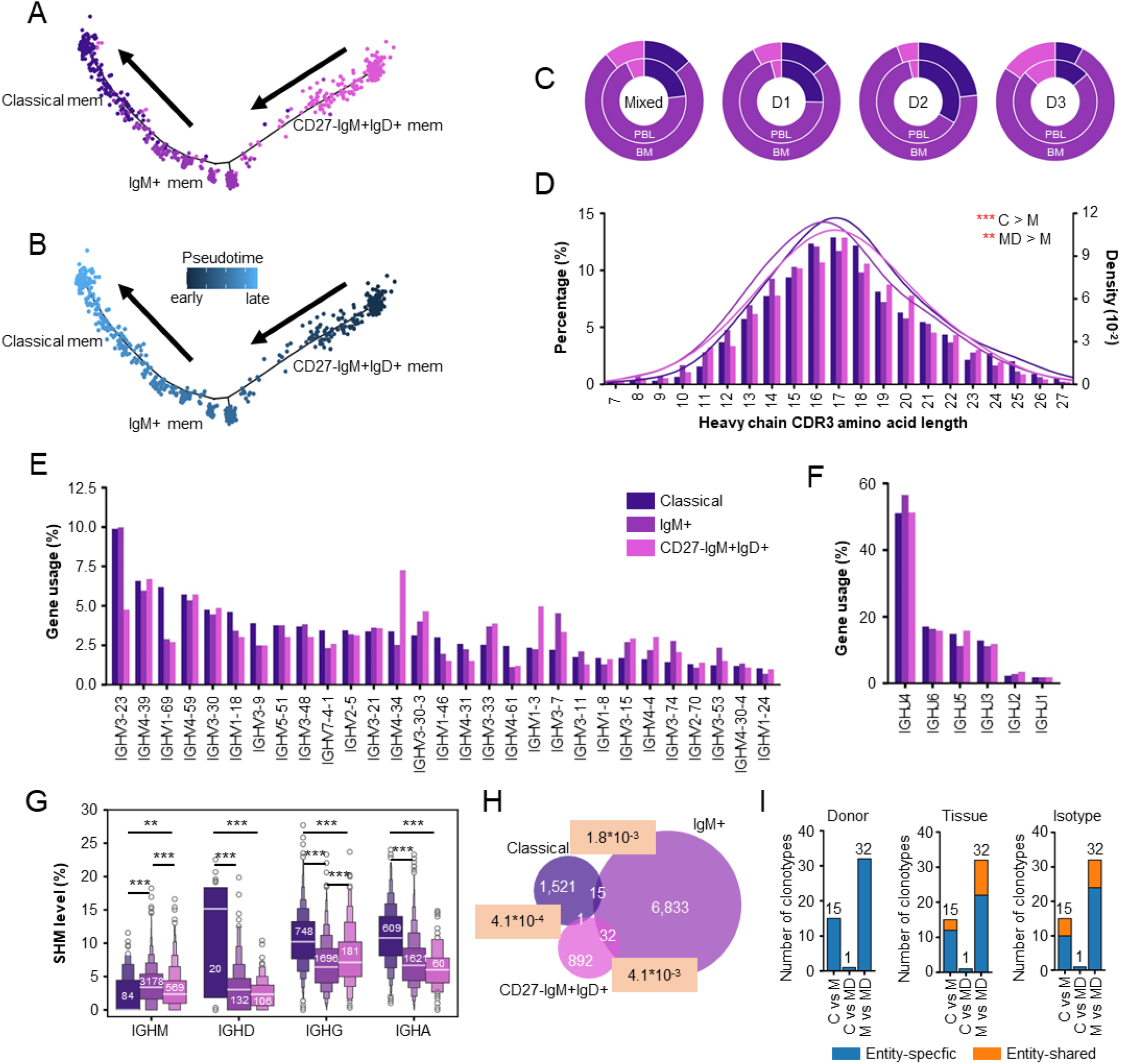
Integrated analysis reveals two compatiable development models for memory B cells. (A-. **B)** Transcriptomic-based inference of developmental trajectory for 3 memory B cell subpopulations with monocle2. **(C)** Doughnut plot showing the tissue-specific composition of three memory subpopulations for the mixed samples and 3 donors. **(D)** CDR3 length distribution. The paired smooth curves are distributions fitted from kernel density estimates. Classical, IgM^+^ and CD27^-^IgM^+^IgD^+^ subpopulations were denoted as ‘C’, ‘M’ and ‘MD’, respectively. **(E-F)** BCR heavy chain variable (H) and joining (I) gene usage for the primary and FOS^hi^ naïve B cell subpopulations. For variable genes, only those with a usage frequency of at least 1% for classical memory were shown. **(G)** Isotype-specific comparison of SHM level of heavy chain variable genes between three memory subpopulations. Numbers on the boxes indicated the number of unique clonotypes. *, *p*<0.05; **, *p*<0.01; ***, *p*<0.001; two independent samples t test, two-sided. **(H)** Venn diagram showing clonotype sharing between classical, IgM^+^ and CD27^-^IgM^+^IgD^+^ memory B cell subpopulations. The numbers with a pink background indicate the clonotype similarity as measured by the Jaccard similarity index. **(I)** Barplots showing the fraction of clonotype sharing between donors, sources and isotypes.

We then investigated the tissue-specific composition of the three memory subpopulations. We observed a higher percentage of M and MD subpopulations in BM compared to PBL (Fig. 5C). This tissue difference can be reproduced in donors separately and also by comparing the in-house BMMC and external PBMC dataset (Fig. 5C and Fig. S14A), consistent with a previous study^32^. Since M and MD subpopulations are not as mature as C, the higher percentage of M and MD suggested BM as a niche with a broader immune potential compared to PBL.

Furthermore, we characterized their BCR repertoires. We found M subpopulation had significantly shorter heavy chain CDR3 length compared to the other two subpopulations (mean values for M, C and MD are 16.8, 17.4, and 17.1) (Fig. 5D). The CDR3 length difference between M and MD subpopulations has been reported in previous studies^33,34^. However, we also observed a significant difference between C and M subpopulations (*p*<0.001). Since they have the same origin, this CDR3 length difference probably reflects the antigen-based selection in the subsequent GC response. For heavy chain gene usage, C subpopulation preferred to use IGHV1 family genes (e.g. 1-69, 1-18, 1-46) (*p*<0.05) and IGHJ5 (*p*<0.01) (Fig. 5E-F). In contrast, M subpopulation preferred to use IGHV3-7 (*p*<0.05), while MD subpopulation preferred to use IGHV4-34 (*p*<0.05). We also compared the VH gene somatic hypermutation (SHM) level between 3 memory subpopulations in an isotype-specific manner. For switched isotypes (IgG and IgA), the SHM level for C subpopulation was higher than both M and MD subpopulations, agreeing with previous studies (Fig. 5G). Remarkably, IgM was found with a higher SHM level in M and MD rather than in C subpopulation. In addition, M had a higher IgM SHM level compared to MD. Finally, we investigated the clonal relationship between these memory subpopulations. C subpopulation was found to have a closer relationship with M than with MD, consistent with the notion that C and M share the same generation pathway (Fig. 5H). However, M subpopulation were found with a higher clone similarity score to MD rather than C. A careful examination confirmed that all 32 clonotypes shared by M and MD came from D3 (Fig. S14B-D). Moreover, we found the two subpopulations had almost equal SHM levels for each of the shared clonotypes (Fig. S14E). A careful examination revealed that all shared clonotypes came from the same donor but could be shared by tissues and isotypes (Fig. 5I), suggesting both the high-level privacy of BCR repertoire and the circulation of mature B cells between BM and PBL. Overall, these results not only supported the well-accepted TD and TI maturation paradigm but also provided evidence correlating M with MD memory B cell subpopulations.

### Myeloid cells represent a pivotal component in B cell development and maintenance

The development and maturation of human B cells are straightforwardly reflected by the phenotype transition of B cell itself, which can occur in genome, epigenome, transcriptome and proteome levels^4^. These multi-omics changes, however, are regulated delicately by different well-organized microenvironments provided by, for example, BM niches and GCs in second lymphoid organs. Signals B cells receive in these microenvironments can be mediated by either the surface- bound ligands from adjacent cells or the secreted signaling molecules such as cytokines, chemokines and adhesion molecules. Having identified both B and surrounding non-B cell types in three sources (i.e. BM, GC and PBL), we are capable of investigating the latent cell-cell interactions (CCIs) that are possibly critical for B cell development and maturation.

Since this study focuses on extrinsic factors contributing to B cell development, we took into consideration only CCIs where B cell serves as a signal receiver and CCIs mediated by adhesion molecules. Applying CellPhoneDB to the CCI (hereafter defined as a unique combination of two interacting molecules) inference, we identified 144, 67, and 61 CCIs for BM, GC and PBL, respectively (see Materials and Methods). The higher number of CCIs in BM revealed a more complex microenvironment maintained by this niche than by the two other sources. Specifically, among 23 non-B cell types in BM, MSC was found with a maximum number (88) of CCIs with B cell subpopulations (Fig. 6A), agreeing with the paradigm that BM stromal cell (BMSC) represents the paramount component in support of B cell development^35^. Following MSCs, the myeloid cell, cDC, was found with 60 CCIs. The large number of CCIs between cDC and B lineage subpopulations also reflected a pivotal role of myeloid cells in maintaining B cell homeostasis in BM. From the point of view of B cell subpopulations, we found cycling pro (69) B cell, three memory B cell subpopulations (75, 71, and 64) and PC (82) had more CCIs with surrounding non-B cells (Fig. 6A). In contrast, the primary and S1008A^hi^ immature B cells and PB presented a minimum number of CCIs.

**Figure 6.**
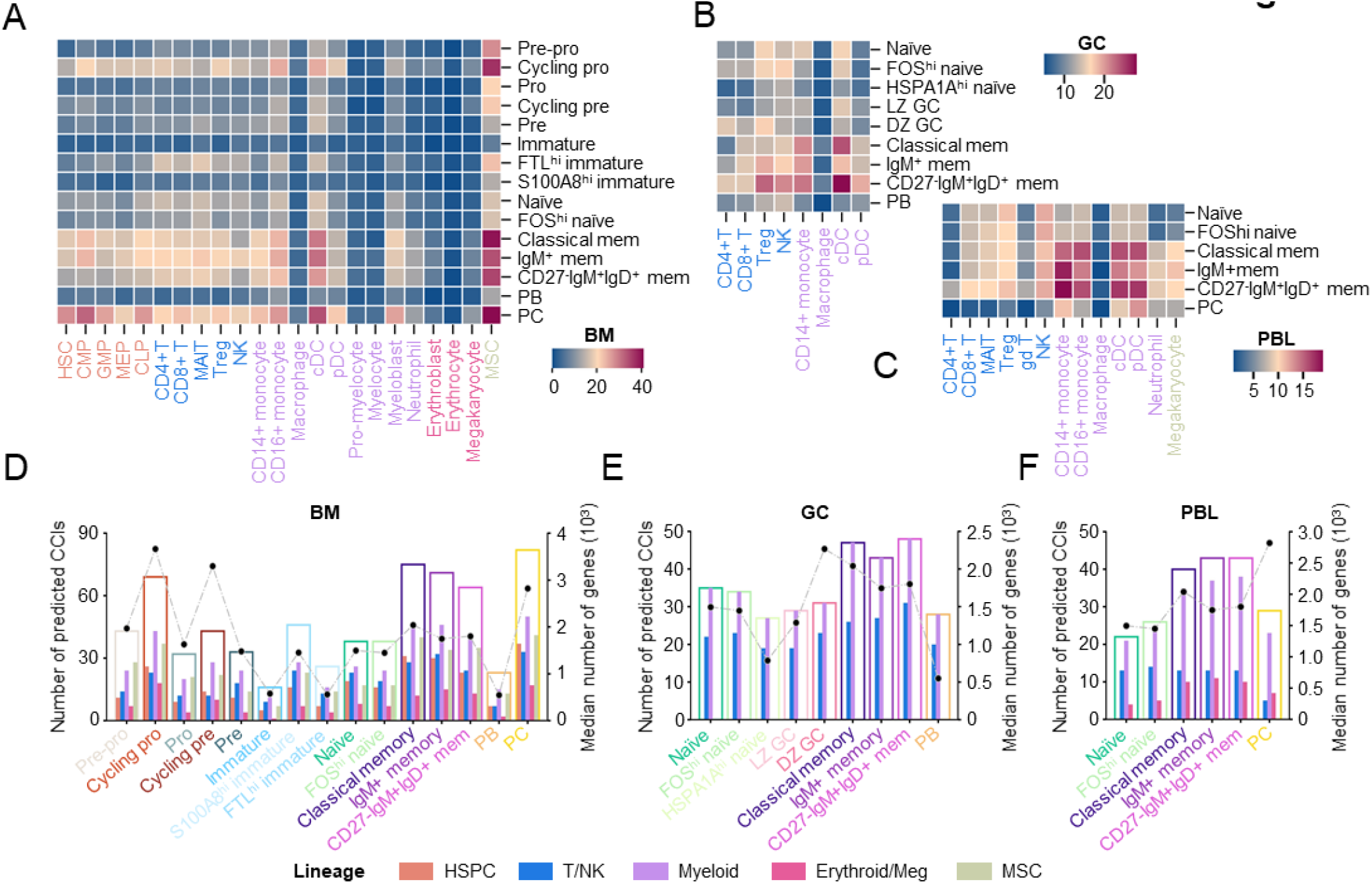
Signaling category between B and non-B cells in three tissues. (A-C) The number of CCIs predicted by CellPhoneDB between B and non-B cells in BM (A), GC (B) and PBL (C) as shown by heatmaps. **(D-F)** The number of CCIs between B and non-B cells in BM (D), GC (E) and PBL (F) as shown in a B cell subpopulation-specific and lineage-specific manner by complex bar plots. The outer hollow bar represents the total number of CCIs predicted for a typical B cell subpopulation. Within the outer hollow bar are solid bars representing the number of CCIs between a typical B cell subpopulation and a typical lineage. The black dot plot overhanging the bar plot denotes the median number of genes for a typical B cell subpopulation.

Resembling BM, myeloid cells also represented a vital lineage interacting with B cells in both GC and PBL (Fig. 6B-C and 6D-F). cDC (38) and CD14+ monocyte (29) were the two cell types predicted with the maximum number of CCIs with B cells in GC and PBL, respectively. Remarkably, for each B cell subpopulation, all CCIs can be recovered in myeloid cells in GC (Fig. 6E). Among B cell subpopulations in GC and PBL, three memory B cell subpopulations remained the subset presenting the maximum number of CCIs (Fig. 6B-C). Typically, LZ GC B cell presented the maximum number of CCIs with cDC (i.e. follicular DC), recapitulating the GC response paradigm^36^ (Fig. 6B). Overall, these results suggested a critical role of the myeloid cells (particular cDC and monocytes) in B cell development and maintenance.

### TNF signaling and adhesion interaction dominate B and non-B CCIs in BM, GC and PBL

We then investigated the signaling mediated by the predicted CCIs. Overall, 40, 20 and 20 signaling categories were identified for BM, GC and PBL, respectively (Fig. S15). By sorting the signaling categories according to the number of subject CCIs, we found signaling by tumor necrosis factors (TNF) was the most frequent signaling category in all three tissues (Fig. 7A-C). The TNF superfamily represents a multifunctional group of cytokines that activate signaling pathways for cell survival, apoptosis, inflammatory responses and cellular differentiation^37^. The enrichment of TNF signaling across tissues suggested an intricate regulation machinery of cell differentiation, survival and apoptosis in B cell development and maintenance. Despite sharing this signaling, the involved CCIs showed discrepancies between tissues. To be specific, BM and GC were found with 3 and 2 tissue-specific TNF CCIs, respectively (Fig. 7E). In BM, these tissue-specific CCIs included TNF_TNFRSF1A, TNFSF10_TNFRSF10D and TNFSF4_TNFRSF4. The former CCI was predicted exclusively for B cell precursors (pre-pro, cycling pro, and pro B cells), whereas the latter two CCIs were exclusively for the terminally differentiated PC (Fig. S16). The two GC-specific CCIs were mediated by the same ligand LTA (TNFSF1) and included LTA_TNFRSF1B and LTA_TNFRSF14 (Fig. 7E). The former was memory B cell-specific while the latter was additionally predicted for naïve B cells (Fig. S16). This stage-dependent regulation by the TNF superfamily could be found not only in the tissue-specific CCIs but also in the shared compartment (Fig. S16).

**Figure 7.**
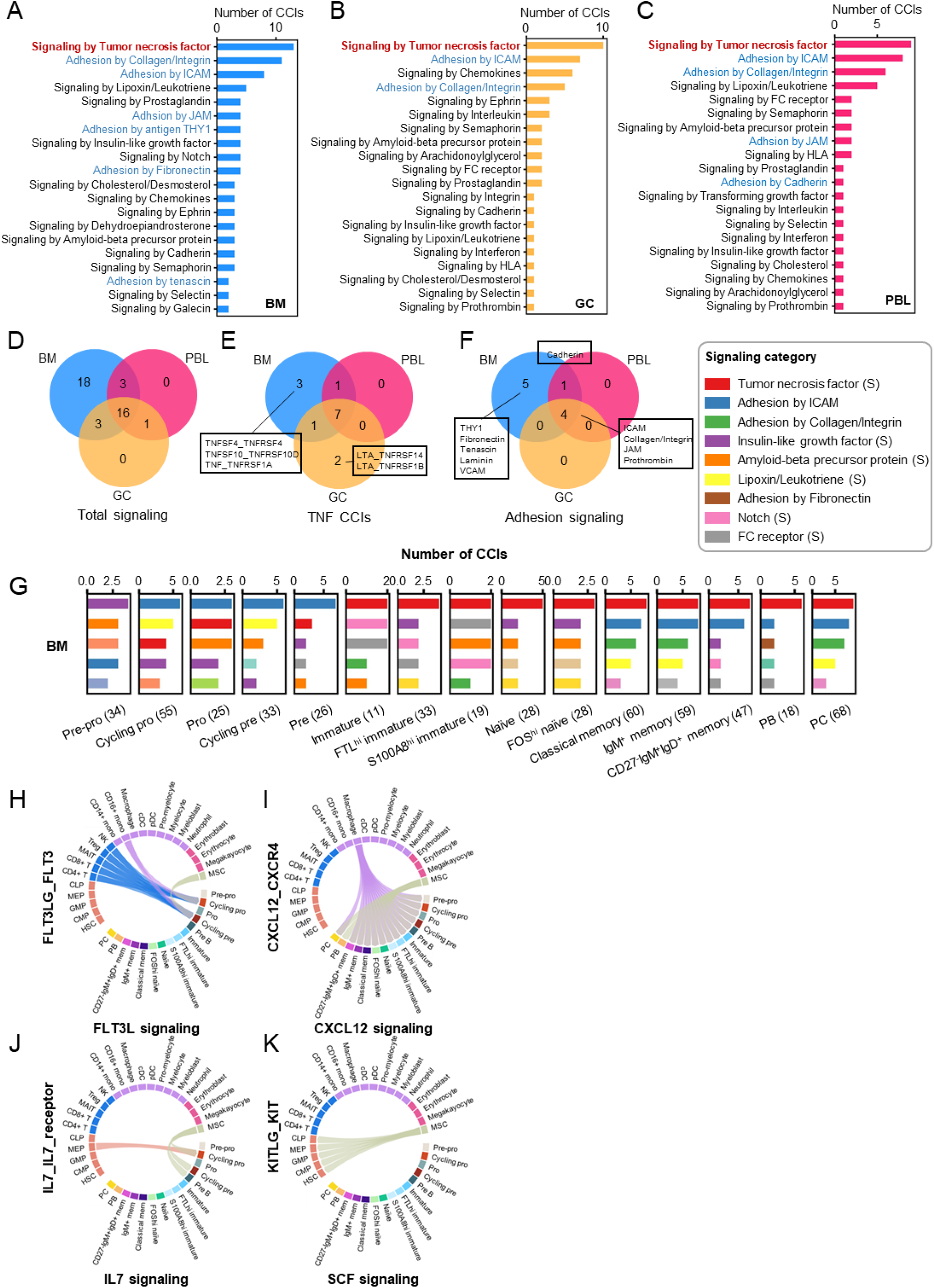
Signaling category between B and non-B cells in three tissues. (A-C) Top 20 signaling categories for CCIs between B and non-B cells in BM (A), GC (B) and PBL (C). The top ‘TNF signaling category’ is highlighted in red. Adhesion categories are highlighted in blue. **(D)** Signaling category overlap between three tissues. **(E)** Tumor necrosis family CCI overlap between three tissues. BM- and GC- specific CCIs are marked. **(F)** Adhesion category overlap between three tissues. All adhesion category subsets are marked. **(G)** Top 5 signaling categories across B cell subpopulations in BM. In parentheses is the number of total CCIs in a typical B cell subpopulation. The signaling category is color-coded as the legend. Notably, only a subset of signaling categories is provided in the legend to save space. For a complete list of signal categories, please refer to Fig. S19. ‘S’ in the parentheses denotes ‘signaling’ in contrast to adhesion CCIs. **(H-K)** Four well-known CCIs (H, FLT3L; I, CXCL12; J, IL7; K, SCF) critical for B cell development in BM predicted by CellPhoneDB as shown by the chord diagrams. Non-B cells are colored according to the lineage. Interactions are represented by ribbons linking two entities, with the one pointed by the arrow being the signal receiver.

Apart from TNF signaling, multiple adhesion CCI categories were also pronounced in the top signaling category list, particularly for BM B cells. Notably, all adhesion CCI categories (n=5, ICAM, Collagen/Integrin, JAM, Prothrombin and Cadherin) predicted in GC and PBL can also be found in BM (Fig. 7F and Fig. S15). Half of the adhesion CCI categories (n=5, THY1, Fibronectin, Tenascin, Laminin and VCAM) predicted in this study were BM-specific (Fig. 7F and Fig. S15), including the well-known VCAM-mediated adhesion^38^ (Fig. S17).

We also investigated the lineage-specific (LS) signaling for non-B cells in three tissues. In BM, 10 signaling categories were found to be lineage-specific, of which 7 were MSC-specific (Fig. S18A). These MSC-specific signaling included 3 aforementioned BM-specific adhesion categories (THY1, Fibronectin and Tenascin) and signaling mediated by notch, R-spondin, transferrin and retinoid acid. While in GC and PBL, the lineage-specific signaling categories were mediated mainly by myeloid cells (Fig. S18B-C). In viewpoint of B cell subpopulations, we found a top signaling category transition from ICAM-mediated adhesion to TNF-mediated signaling along B cell development in BM. The enrichment of ICAM-mediated adhesion further addressed the its fundament role in early B cell development (including Cycling pro, cycling pre and pre B cells) (Fig. 7G). As is in BM, TNF- and ICAM-mediated signaling also dominate in B cell subpopulations in GC and PBL (Fig. S19). Remarkably, DZ GC B cells had an increased number of ICAM-mediated CCIs compared to its LZ counterpart, indicating a requirement for more delicate adhesion pattern for this proliferating subpopulation.

Five well-known essential factors for B lineage commitment and development are FLT3L, CXCL12, IL-7, SCF and RANKL^35^. All five factors were successfully reported by CellPhoneDB even though RANKL signaling was not significant (*p*>0.05, not shown here) (Fig. 7H-K). Notably, SCF (KITLG_KIT) signaling was predicted only between MSC and HSPC (Fig. 7K). The missing of SCF signaling predicted between MSC and early B cells might reflect its elusive role in early B cell development given that SCF-KIT axis is redundant in fetal, neonatal and young mice^39^ but seems essential in adult mice^40^. It is widely accepted that the delicate niche required for early B cell development is provided by the coordinated network of different BMSCs, which includes osteoblasts, reticular cells and IL-17-expressing cells^35^. Consistent with this notion, MSC, which is comprised of BMSCs here, took part in all these canonical signaling pathways (Fig. 7H-K). However, additional cell types of other lineages were also found as potential signal senders (FLT3L by T/NK cells and CD16+ monocytes, CXCL12 by macrophages and IL7 by MEP). These results indicate a role for cell populations other than BMSCs in the niche construction for B cell development.

### S100A8^hi^ immature B cells represent an age-associated B cell subset

The calprotectin (S100A8/A9) is constitutively expressed by myeloid cells, especially in neutrophils, monocytes and macrophages^41–44^. Although it was previously reported that this protein was also detected in B cells in pathological conditions^45,46^, we assumed that its expression in B cells in the physiological state represented a kind of unusual ectopic expression and deserved further investigation. Considering the characteristics of enrolled subjects (more than 50 years) in this study and S100A8/A9’s association with aging in a previous report^47^, we hypothesized that this S100A8^hi^ immature B cell subpopulation represented an age-associated B cell (ABC) subset and the ectopic expression of S100A8 in B lymphocyte in physiological state is relevant to senescence-associated secretory phenotype (SASP). To verify this, we conducted *in silico* analyses and qPCR and *in vitro* cell culture experiments.

Firstly, we validated the existence of S100A8^hi^ B cells (C8 and C12) in two independent single- cell studies^18,20^ (Fig. 8A and 8B). For the former study, the cells were derived from the BMMC of three healthy subjects used as controls in this study. For the latter one, the cells are derived from hematologically healthy BM of patients undergoing total hip arthroplasty. All subjects from both studies are older than 50 years. To consolidate the verification of its association with age, we also dissected BM samples of younger subjects (with age from 2 to 50) from another four available single-cell studies^10,19,21,22^ and found no evidence supporting S100A8 expression in B cells (Fig. S20). Due to the limited accessibility to human BM samples, we investigated the age-associated expression of S100A8/A9 in B cells from multiple tissues (spleen, kidney, mammary gland, limb muscle, lung and so on) of mice of different ages from an aging study^48^. We observed a significant age-associated expression pattern in mRNA level (Fig. 8C, Spearman rank-order correlation; CC, 0.89; *p*<0.05). Inspired by this mouse-based result, we performed quantitative PCR (qPCR) and FACS on B cells purified from BM, spleen, and PBL of a cohort of C57BL/6 mice of different ages (3m, young (Y); 12m, middle age (M); 22m, aged (A), n=3 for each group). The qPCR experiment revealed a higher S100A8/A9 expression level in the aged group than in the middle age group (S100A8, 0.065 (A) vs 0.049 (Y) vs 0.041 (M); S100A9, 0.031 (A) vs 0.022 (M) in spleen) though that the young group was shown with a comparable high S100A9 expression level as their aged counterpart (0.040 (Y) vs 0.031 (A) in spleen) (Fig. 8D). However, we did not observe the same trend in BM B cells (Fig. S21). In the FACS experiment, the proportion of the S100A8-positive population in total B cells (CD45+CD19+) was measured and we observed a higher percentage of such cells in the aged group than the middle age group in all three tissue sources (PBL, 37.6% vs 13.5%; BM, 25.1% vs 14.8%; SP, 27.2% vs 37.4%) (Fig. S22).

**Figure 8.**
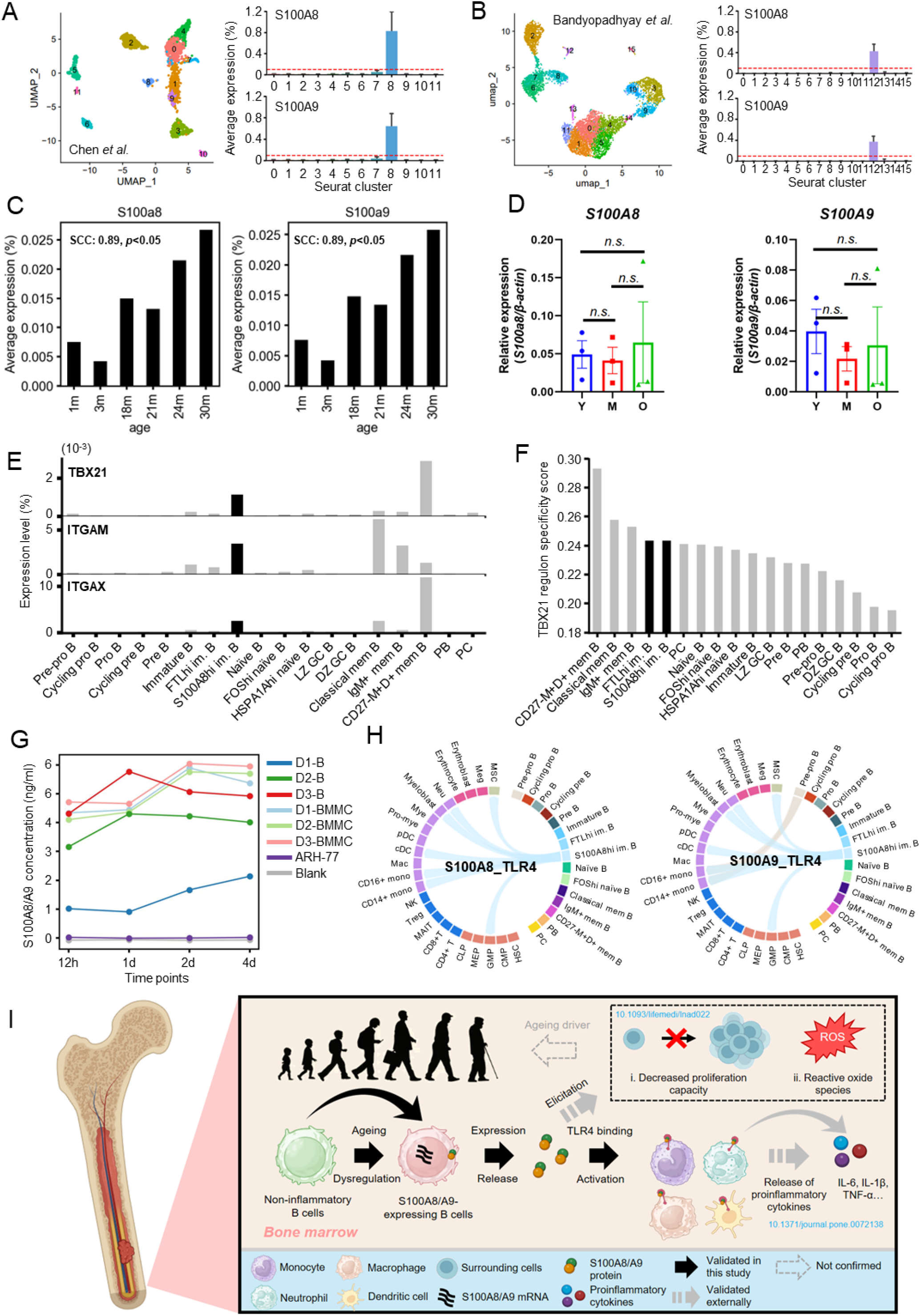
Age-associated expression of S100A8/A9 in B lymphocytes. (A-B) UMAP of BM B cell clusters from aged healthy human subjects from two external single-cell studies and the average expression of S100A8 and S100A9 across clusters. The red dashed lines mark an average expression level of 0.1%. **(C)** Age-associated expression of S100A8/A9 in mouse B cells from a mixture of tissues as measured based on scRNA-seq data. SCC, Spearman’s correlation coefficient. **(D)** Age-associated expression of S100A8/A9 in the splenic B cells of C57BL/6 mice as measured by qPCR. Y, young (3m); M, middle age (12m); O, old (22m). **(E)** The average expression level of three ABC marker genes (TBX21, ITGAM, and ITGAX) across B cell subpopulations. **(F)** TBX21 regulon specificity score comparison across B cell subpopulations. **(G)** ELISA assay demonstrating the ability to secrete S100A8 for human BM B cells. The concentration of S100A8/S100A9 in the cell culture medium was measured in four time points (12h, 1d, 2d and 4d) for each sample. ARH-77 and blank culture medium were used as negative controls while the unsorted human BMMC was used as a positive control. **(H)** Chord diagram showing the predicted S100A8/A9-TLR4 signaling between B cell subpopulations and surrounding bone marrow non-B cell types. **(I)** Schematic diagram demonstrating the occurrence of S100A8^hi^ B cells in the aged population and how they participate in the inflammageing process.

Since ABCs have been reported in previous studies (reviewed by de Mol *et al*.^49^ and Yu *et al*.^50^), we compared their molecular expression signature with S100A8^hi^ immature B cells to evaluate their developmental relationship. A widely accepted gene set, including TBX21, ITGAM and ITGAX, was reported to distinguish ABC from other B cell subsets in both humans and mice^49^. We thus profiled the expression of the three genes across B cell subpopulations and found weak expression of ITGAM and ITGAX in classical memory and CD27-IgM+IgD+ memory B cells, respectively (Fig. S23). With a high-resolution mean expression comparison, however, relatively higher expressions of these genes were found with S100A8^hi^ immature B cells than other non-memory B cell subpopulations, demonstrating a similar molecular expression signature between this subpopulation and known ABC (Fig. 8E). Among the 3 genes, TBX21 encodes a TF sharing a common DNA-binding domain, the T-box. To investigate its transcription regulation activity, we performed single-cell regulatory network inference and clustering (SCENIC) analysis and found S100A8^hi^ immature B cells had the highest regulon specificity score among all non-memory B cell subpopulations (Fig. 8F). Notably, it has been documented that in healthy adults a small but readily identifiable memory B cell subset expresses T-bet (TBX21)^51^. This probably accounts for the high expression of these markers and high TBX21 activity in memory B cell subpopulations. Therefore, the highest expression level of known ABC marker genes and the highest transcription activity among all non-memory B cells possibly suggested a shared mechanism contributing to their aged phenotypes.

Although we confirmed the age-associated expression of S100A8/A9 in B lymphocytes, it remains unclear this unusual expression pattern can result in the secretion of this molecule into the microenvironment and then contribute to SASP. To address this issue, we conducted short-term cell culture experiments with a serum-free medium (see Materials and Methods). Unsorted human BMMC and blank medium were used as a positive and a negative control, respectively. A human B lymphoblast cell line, called ARH-77, was also included as a negative control. S100A8/A9 level in cell supernatant or blank medium was measured in 12 h, 1 d, 2 d and 4 d by ELISA. The result demonstrated that the S100A8 level in human B cell supernatant was clearly higher than the blank medium but was lower than their respective BMMC control, supporting the secretion ability of B lymphocytes (Fig. 8G). Following this result, we further predicted S100A8/A9_TLR4 signaling between S100A8^hi^ immature B cells and certain myeloid cell types (monocyte, neutrophil, DC) in our own data with CellPhoneDB (Fig. 8H). Moreover, Zhang *et al*. have revealed an aging driver role of S100A8/A9 with the recombinant human antibody by in *vitro* experiments^47^. They demonstrated that S100A8/S100A9 could elicit oxidative stress, inflammatory response, and proliferative impairment. Considering these facts, the secretion of S100A8 from B lymphocytes from the aged probably participates in the senescence process through multiple mechanisms. (Fig. 8I). Taken together, these results implicate an elevated expression of S100A8 (and S100A9) in B lymphocytes in the aged population and an involvement of B lymphocytes in senescence.

## Discussion

Single-cell sequencing technology significantly advanced our knowledge of the development and maturation of B lymphocytes, shedding light on their heterogeneity, developmental trajectory and BCR repertoire features in both physiological and diseased status. However, a comprehensive single-cell and high-dimensional examination of human B lymphocytes from the earliest progenitors to terminally differentiated PCs was lacking. In this work, we compiled scRNA-seq and scVDJ-seq data of B cells from human BM, PBL and tonsil and provided an integrative analysis of the gene regulation, conventional B cell heterogeneity and cell-cell communication network along the B cell development axis. We found a least active stage for immature B cells, a local and subject-specific HP pattern for naïve B cells, and two compatible development models for memory B cell subpopulations. Moreover, we also identified the myeloid cells as a key cell category in addition to stromal cells in B cell development and maturation, where TNF and adhesion signaling dominate all external signaling types and prevail over the other in a stage-dependent manner. Last but not least, we identified two age-associated B cell subpopulations, S100A8- and C1q-expression B cells, which probably contribute to the SASP.

The developmental trajectory of B cell progenitor in BM has been constructed in previous studies^7,52,53^, along with which the gene expression kinetics was depicted, particularly for those TFs. However, the dynamics of transcription activity when B cell develops and matures remains unresolved. We reported immature B cells as the least transcriptionally active stage along the developmental axis, featured by the minimum number of expressed genes (Fig. 2A) and the lowest level of gene activity as measured by RNA velocity (Fig. 2E). This observation coincided with the lowest metabolic rate for immature B cells demonstrated by Zeng *et al*.^14^ (Fig. 1A in their publication). Given that both heavy and light chain gene arrangements are completed, this downregulated gene expression activity in such a stage reflects a resting state poised for efflux from BM and subsequent peripheral tolerance before entering mature B cell repertoire.

B-cell HP is reported in both physiological^54^ and B-cell deficit^55,56^ settings. It takes place in response to B cell depletion and represents a mechanism to compensate for this cell loss, thereby inhibited in a cell dose-dependent manner by feedback from mature B cells^26^. We provided evidence of HP by demonstrating clonal sharing between different naïve B cell subpopulations that showed no antigen-dependent proliferation history (Fig. 4K). We further found that the clonal sharing between these naïve phenotype subpopulations was tissue-confined in both BM and PBL. Since HP has only be reported previously for peripheral mature B cells^54^, our finding that naïve B cell HP co-occurs in BM extended the scope for this particular process. Because periphery B cell HP has been related to lymphogenesis regulation in BM^55,56^, this local B cell HP could have a more straightforward role in this feedback. This hypothesis, together with the interplay between BM and peripheral HP, however, remains to be investigated. For the HSPA1A^hi^ naïve B cell subpopulation, Yang *et al*. recently reported a stress-response memory B cell subset with a similar gene expression pattern in tumor microenvironment (TME) in a pan-cancer study^57^. In their work, the stress response was suggested as a common molecular characteristic across different cell lineages in tumors. Similarly, the HSPA1A^hi^ naïve B cell subpopulation in this study comes from pediatric patients with recurrent tonsillitis and the inflammatory microenvironment could have mimicked that observed in TME.

The heterogeneity of memory B cells has long been appreciated. Using single-cell transcriptomic data, we unbiasedly identified three memory B cell subpopulations, namely classical (C), IgM+ (M), CD27-IgM+IgD+ (MD). The subsequent development trajectory analysis and BCR repertoire characterization presented substantial evidence supportive of a close relationship between C and M subpopulations, including a transitional developmental pathway (Fig. 5A), a similar Ig subclass composition (Fig. 5J), and a closer clonal relationship (in two of three donors) (Fig. S13B- C), agreeing with the notion that they are both post-GC and developmentally continuous^28^. However, we also noticed the non-negligible clonal relationship between M and MD subpopulations in one of the three enrolled subjects (Fig. S13D). By examining the shared clonotypes (n=32), we found a substantial fraction of them carried distinct constant genes and varied SHM levels in the two memory subpopulations, suggesting a dynamical transition between the two (Fig. S13E). As is reviewed by Seifert and Küppers^58^, the origin of MD subpopulation is disputed among three viewpoints, consisting of a means of naïve B cell diversification, T-dependent and T-independent immune responses. Here, we proposed distinct but compatible models to account for the origin of MD subpopulation.

This integrated single-cell data also enabled us to investigate the CCIs involved in the development and maturation of B cells. According to the conventional paradigm, BMSC is the most paramount component in maintaining the niches required for B cell lymphogenesis, encompassing adipocytes, endothelial cells, osteoblasts and fibroblastic reticular cells^59^. Upon encountering antigen, activated B cells will undergo several rounds of proliferation and affinity-based selection facilitated by follicular helper T cell (T_FH_) and DC (FDC) in the GC^36^. However, several non-canonical cell types were also reported to reside in GC, such as macrophages, natural killer and CD8+ T cells (reviewed by Victora *et al*.^60^). In the circulating blood, direct DC-to-B cell contact was also observed in mice with flow cytometry and microscopic imaging^61^. With the latest manually curated interaction database, we predicted a great number of candidate CCIs between B cells and myeloid cells across the three representative tissues (i.e. BM, GC and PBL) (Fig. 6D-F), implicating a pivot role for myeloid cells in B cell lymphogenesis and peripheral homeostasis. Moreover, we also detected a top CCI category shift from adhesion interaction to TNF signaling (Fig. 7G), clarifying a particular feature of the external signal dynamics along the B cell development axis. Overall, our analysis revealed the complexity of the microenvironment supporting the development and maturation of B cells.

Lastly, it is worth mentioning that we identified two age-associated B cell subpopulations that express S100A8/S100A9 and C1q, respectively. Both proteins are typically expressed in myeloid cells, such as macrophages, monocytes, neutrophils and DCs. To the best of our knowledge, their expression in B cells is exclusively reported in diseased settings or organ transplantation cases (i.e S100A8/S100A9 in SLE^45^ and COVID-19^46^; C1q in transplantation^62^). However, we identified the two subpopulations in donors without reported inflammatory responses. For the S100A8-expressing subpopulation, we validated it in a subset of external BM scRNA-seq data from aged human donors and found a significant correlation between S100A8/S100A9 expression level and age in mice. Moreover, we experimentally validated the S100A8/A9 secretion ability of B cells, suggesting its role in SASP. In contrast, the C1q-expressing subpopulation was not found in any enrolled external dataset, which could be accounted for by its particularity and rarity. Both subpopulations were found in BM and had a transcriptomic phenotype similar to the primary immature B cell subpopulation. Whether this observation reflects their niche dependency remains to be elucidated.

It should be noted that certain limitations are present in our work. Firstly, this study does not cover all well-known B cell subsets, i.e. transitional B cells, regulatory B cells and B1 cells. To an unbiased delineation of the gene regulation dynamics, we did not delve into the data for these minor subpopulations. Second, the existence and functionality of novel B cell subpopulations present in this study remain to be validated. Similarly, our CCI analysis provides only candidate interactions, which take place when some prerequisites like cell juxtaposition are met. Moreover, the impact of these candidate interactions on B cell development also calls for careful examination.

Despite these limitations, our work reveals the heterogeneity of conventional human B cell subsets, resolves gene regulation dynamics and underlying CCIs along the B cell development axis, and sheds light on the attributes of several critical B cell biological processes. Thus, it represents a valuable resource for in-depth investigation of B cell development.

## Materials and Methods

### Sample collection and B cell enrichment

All human studies were performed in compliance with the guidelines of the Research Ethics Committee of Guangdong Provincial People’s Hospital. BM and matching PBL were collected from three patients who were more than 50 years old suffered from a herniated disk but were free from hematological disorders. Informed consent was obtained from all participants prior to sample collection. The isolation of PBMCs and BMMCs was performed using standard density-gradient centrifugation with Ficoll-Paque™ (Catalog. LTS1077 for PBMCs, TBD2013LHU for BMMCs, TBD). B lymphocytes were purified using the Human B Lymphocyte Enrichment Set-DM kit (Catalog. 558007, BD Biosciences) from fresh BM or PBL, following the manufacturer’s instructions. Cell counting and trypan blue exclusion viability were performed on TC20 Bio-Rad automated cell counter.

### Single-cell transcriptomic and immunomic library preparation and sequencing

To evaluate the concentration and viability of cell suspensions, filtered trypan blue was added and an automated cell counter (Bio-Rad, TC20) was used. The optimal concentration range for cell stock was determined to be 700-1,200 cells/μl to increase the likelihood of reaching the desired cell recovery target. A minimum sample viability of ≥90% was required to improve the recovery rate. Based on the recommended cell concentration and cell counter results, cells were resuspended in a suitable volume of cold PBS containing 0.04% BSA. For the analysis of B lymphocytes and BMMCs, cells were loaded onto the 10x Genomics chromium single-cell platform. To ensure sufficient cell numbers, two portions of BMMCs and four portions of enriched B lymphocytes (3 for BM and 1 for PBL) were loaded onto the platform. For BMMC analysis, libraries were constructed using the Chromium Next GEM Single Cell 5’ Kit v2 (10x Genomics, 1000263). And for B lymphocyte analysis, libraries were constructed using the Chromium Next GEM Single Cell 5’ Kit v2 (10x Genomics, 1000263) and the BCR Amplification kit (10x Genomics, 1000253). Briefly, GEM generation and barcoding were performed with 10,000 cells per reaction, followed by GEM-RT, post-GEM-RT cleanup, and cDNA amplification to isolate and amplify cDNA for library construction. Complementary DNA post-amplification, cDNA post-target enrichment, and final libraries were evaluated using Agilent Bioanalyzer chips (Catalog. no. 5067-5582) on an Agilent Bioanalyzer 4200. Finally, the libraries were sequenced using an Illumina platform with 150 bp pair-end sequencing.

### scRNA-seq data processing and cell population identification

Single-cell gene expression matrices for enriched B cell and unsorted BMMC samples were obtained using CellRanger (v6.1.2) ‘count’ utility with ‘refdata-gex-GRCh38-2020-À as the reference transcriptome. The external scRNA-seq data of early B cell samples was obtained from GSA repository (https://ngdc.cncb.ac.cn/gsa-human/, accession number: HRA000489) as raw sequencing reads in fastq format, of which the expression matrices were obtained following the procedure same as in-house data. The external scRNA-seq data of tonsil samples was obtained from ArrayExpress (https://www.ebi.ac.uk/biostudies/arrayexpress, accession number: E-MTAB-9005) as processing-ready expression matrices. The scRNA-seq data of PBMC from healthy subjects was obtained from CNGB Nucleotide Sequence Archive (https://db.cngb.org/cnsa, accession number: CNP0001102) as a processed Seurat object.

With all scRNA-seq data ready, we utilized Seurat (v4.1.1) to perform data preprocessing, integration, and subsequent procedures contained in a routine Seurat analysis pipeline. Specifically, we retained in each sample only the cells expressing at least 200 genes and the genes expressed in at least 3 cells. Cells expressing a high percentage (>=10%) of mitochondrial genes or predicted to be doublets by DoubletFinder (v2.0.3) were also discarded in the preprocessing step. Moreover, we removed BCR/TCR-related genes (including variable, junctional and diversity genes, with constant region genes retained) to avoid bias in the subsequent clustering step. Notably, the preprocessed PBMC dataset was not subject to the above quality control. Afterwards, the preprocessed expression matrices were integrated following Seurat ‘Fast integration using reciprocal principal component analysis (PCA)’ practice with default parameters. Notably, we specified the reference to be the unsorted BMMC sample (containing multiple cell types) with the maximum number of cells (i.e. “D2-BM2”, see Table S2) in the integration anchor identification step (‘FindIntegrationAnchors’). Subsequently, the integrated expression values were further processed with a routine workflow that includes data scaling, linear (PCA) and non-linear (UMAP) dimensional reduction and cell clustering, which were achieved by ‘ScaleDatà, ‘RunPCÀ, ‘RunUMAP’, ‘FindNeighbors’ and ‘FindClusters’ utilities with default parameters. Only 30 PCs were retained in the PCA step. The resolution in the cell clustering step was set to 0.8. Cell clusters identified in the first round of clustering were then manually annotated with the lineage information (i.e. HSPC, B, T/NK, myeloid, erythroid/megakaryocyte and MSC) according to the lineage markers (Fig. S2A).

After the lineage identification, an unbiased cell population identification within individual lineages (except for MSC) was conducted through five parallel secondary rounds of clustering. To be specific, cells from a typical lineage were extracted and the expression values from cells of different projects were integrated as that in the first round of clustering. The integration basis transitions from the sample to the project level to avoid the bias caused by the rarity of cells in some samples after the cell lineage split. The integrated data were then subject to the same workflow as above to identify cell clusters in a typical lineage with canonical marker genes (Fig. S2B-E). For B cell subpopulations, known marker genes (Fig. 1F) were used in combination with the computed differentially expressed (DE) genes (with the ‘FindAllMarkers’ utility) (Fig. S4) to define their identities. With all lineage annotation finished, cell populations of different lineages were recombined into a well-annotated integrated dataset ready for downstream analyses.

### scVDJ-seq data preprocessing, clonotype assembly and downstream analyses

CellRanger (v6.1.2) ‘vdj’ utility was employed to obtain VDJ contigs (‘filtered_contig.fasta’) from scVDJ-seq datasets. Subsequently, novel allele identification, genotyping and gene reassignment for heavy chain V genes were performed successively to achieve an accurate VDJ repertoire profiling (particularly for SHM level evaluation). IgDiscover (v0.15) was selected to infer novel V alleles for each donor based on bulk sequencing data as recommended by Yang *et al.*^63^. Then, we implemented a genotyping method as Zhu *et al*.^64^ before VDJ gene reassignment by IgBLAST with individualized reference sequence sets. The corrected V gene assignments for heavy chain contigs were a basis for SHM-level quantification. After correcting gene assignment, we further filtered the data to retain one heavy and one light chain contig for each cell. Cells without productive heavy or light chain contig or with multiple heavy or light chains supported by a similar number of umis (fold change<2, refer to the criteria employed in Dandelion^65^) will be discarded. The preprocessed contigs were then integrated with the annotated scRNA-seq data to assign B cell subpopulation labels. As a result, contigs from cells not captured by scRNA-seq data were also removed. Finally, the resulting clean and well-annotated contigs will be assembled into clonotypes, within each they share the same V and J genes and CDR3 amino acid sequences in both heavy and light chains. The downstream BCR repertoire analyses (including gene usage, CDR3 length distribution, clonotype sharing and SHM level comparison) were all on a clonal basis, which meant that a unique clonotype provides only an observation on the studied feature.

### RNA velocity analysis with scVelo

The spliced and unspliced expression matrices were obtained by running the Python package velocyto^66^ (v0.17) first on each sample individually (samples from King *et al*. were not included here due to the unavailability of raw sequencing data). The output loom files were combined using the ‘combine’ function in loompy (v3.0.7) and imported using the ‘read’ function in scVelo^67^ (v0.2.4). To take advantage of pre-computed dimensionality reduction UMAP embedding, the AnnData storing combined spliced/unspliced matrices were merged with the AnnData storing cell metadata and the associated UMAP embedding using the function ‘utils.merge’. Since this step resulted in a preprocessed AnnData object (retained only highly variable genes used to integrate samples and pre-computed PCA), we did not implement further preprocessing as indicated in the tutorial. Subsequently, each cell’s moment (means and uncentered variances) was computed using ‘pp.moments’ with default parameters, and these moments facilitated RNA velocity estimation implemented in the function ‘tl.velocity’, with mode set to ‘stochastic’. Based on estimated velocities, a velocity graph representing transition probabilities was constructed using the function ‘tl.velocity_graph’. The velocity graph was then used to embed RNA velocities in the pre-computed UMAP in the form of streamlines, with the function ‘pl.velocity_embedding_stream’. Top active/important genes were identified using the function ‘tl.rank_velocity_genes’. The velocity pseudotime was calculated using the function ‘tl.velocity_pseudotime’ with default settings and tuned manually by setting ‘root_key’ to be a selected pre-pro B cell and ‘end_key’ to be a selected plasma cell.

The Seurat object of single-cell data was first converted into ‘.h5ad’ format that is accessible by the Python-based single-cell data processing toolkit Scanpy^68^ (v1.9.1) with ‘SaveH5Seurat’ and ‘Convert’ utilities embedded in R package ‘SeuratDisk’ (v0.0.0.9020). The converted single-cell data was then imported with the ‘read_h5ad’ function in Scanpy. The partition-based graph abstraction was computed and visualized with the ‘tl.paga’ and ‘pl.paga’ utilities with default parameters, respectively. Notably, the weight threshold was set to 0.09 for a concise visualization of the connectivity between B cell subpopulations.

### Development trajectory inference with Monocle2

The development trajectory for immature, naïve and memory subpopulations was inferred by Monocle^69^ (v2.24.1) with default parameters. To improve computing efficiency and obtain a neat trajectory, 200 cells were randomly selected from each B cell subpopulation and the corresponding raw expression matrix was first extracted to construct a CellDataSet. Then, ‘estimateSizeFactors’ and ‘estimateDispersions’ were employed to normalize the data with the total library size and estimate negative binomial overdispersion for each gene, respectively. All variable genes defined by the differentialGeneTest function (cutoff of q<0.01) were used for cell ordering with the ‘setOrderingFilter’ function. Dimensionality reduction was performed with the DDRTree method in the ‘reduceDimension’ step. Cells were then represented onto a pseudotime trajectory using the ‘orderCells’ function and visualized using the ‘plot_cell_trajectory’ function.

### GO enrichment analysis

GO enrichment analysis in this study was performed by using the online web server Enrichr^70^ (https://maayanlab.cloud/Enrichr/, based on an update on June 8, 2023). DE genes for a typical B cell subpopulation were computed with Seurat embedded function ‘FindAllMarkers’ among only the compared cell subpopulations (for example, only three naïve B cell subpopulations were considered when computing DE genes for a typical naïve B subpopulation). All derived DE genes (padj>0.05) for a B cell subpopulation were used as the input in this analysis. Enriched terms were sorted by *p* values.

Single-cell gene regulatory network analysis was performed following the published pySCENIC (v0.12.1) protocol^71^. To save the runtime, 1000 cells from each B cell subpopulation were randomly sampled. Log-transformed counts were used as the input. The Seurat object of sampled cells was converted into ‘.loom’ format using ‘as.loom’ utility to enable data import by the recommended workflow. Then gene regulatory network inference, candidate regulon generation and regulon prediction, and cellular enrichment steps were consecutively executed with the utility ‘grn’, ‘ctx’, and ‘auc’ from the command line. The list of TFs, ranking databases and motif annotation were consistent with the protocol. Subsequently, regulon specificity scores across B cell subpopulations were calculated through the embedded ‘regulon_specificity_scores’ function based on cellular enrichment scores. Only regulons containing C1q components were considered for PPARG and NR1H3 in regulon network construction (visualized by Cytoscape (v3.9.1)).

### Analysis of external human and mice scRNA-seq data

The external human BM scRNA-seq data were obtained in three formats: raw sequencing reads^20,21^, gene expression matrices^10,19^ and well-annotated Seurat objects^18,22^. For data type of raw sequencing reads, we first obtained the gene expression matrices following the method described previously. The resultant gene expression matrices, together with the downloaded ones, were subjected to a standard scRNA-seq data analysis workflow, which includes data preprocessing, library size normalization, highly variable gene identification, dimensional reduction and unsupervised clustering. Notably, different samples within a dataset were integrated by the harmony (v1.2.0) algorithm for efficiency. After the primary clustering, B lineage cells were extracted and were subjected to a secondary clustering to identify B cell subpopulations. For the well-annotated Seurat objects, B cells were straightforwardly extracted based on the cell type label provided by original authors. Afterward, the extracted B cells were subjected to a secondary clustering to identify B cell subpopulations for examining C1QA, C1QB, C1QC, S100A8 and S100A9 expression. For the scRNA-seq data of mice of different ages, the processed h5ad format data was directly obtained through figshare website (https://figshare.com/projects/Tabula_Muris_Senis/64982). This well-annotated single-cell data was then imported and processed by Scanpy (v1.9.1). Firstly, B lymphocytes were extracted according to the cell label and the expression levels of S100A8/A9 for each cell were calculated by ‘sc.pp.calculate_qc_metrics’ utility. Subsequently, an average expression level of S100A8/A9 was calculated and its correlation (Spearman’s rank-order correlation) with age was measured using the ‘stats.spearmanr’ function in the ‘scipy’ (v1.7.3) module with Python (v3.9.7) programming.

### Prediction of non-B and B cell interactions with CellPhoneDB

Non-B and B cell CCIs were predicted by CellPhoneDB^72^ (v5.0.0, https://github.com/ventolab/CellphoneDB) in this study. Before this prediction, we split the entire dataset into three subsets (i.e. BM, GC and PBL) according to the tissue source to avoid false positive CCI predictions caused by tissue discrepancy (e.g. CCIs between cell populations not co- existing in the same tissue or CCIs constituted by molecules not co-expressed in the same tissue). After this dataset splitting, cell populations with a percentage less than 0.1% were discarded for a reasonable (removing cell populations that are generally believed to be not present in a typical tissue) and reliable (retaining only CCIs between cell populations with a certain frequency) prediction. The consequent clean datasets were used in downstream CCI prediction by CellPhoneDB statistical framework with default parameters. Because the major concern of this study is factors contributing to B cell development, we considered only CCIs where B cell serves as a signal receiver and CCIs mediated by adhesion molecules. The signaling directionality was inferred from column ‘receptor_a’ and ‘receptor_b’ in the output file (statistical_analysis_pvalues.txt). Adhesion CCIs had having their ‘directionality’ annotated as ‘Adhesion-Adhesion’ or ‘classification’ containing the key word ‘Adhesion’. In-house scripts were developed to process and visualize the prediction result from CellPhoneDB.

### Cell isolation from mouse organs and fluids

Male C57BL/6 mice were purchased from the Guangdong Medical Laboratory Animal Center. All animal experiments were performed in strict accordance with the ethical guidelines of Guangdong Provincial People’s Hospital and approved by the Ethics Committee of Guangdong Provincial People’s Hospital. These experiments adhered strictly to the principles of animal research. The mice were housed in cages under specific pathogen-free (SPF) conditions. Young mice (Y) were 3 months old, middle mice (M) were 12 months old, and old mice (O) were 22 months old.

Murine PBL was aseptically collected from the inferior palpebral vein and dispensed into EDTA-coated tubes to prevent coagulation. Subsequently, the total BM was meticulously flushed from the marrow cavities of the femurs and tibias employing a calibrated 1 ml syringe. This BM suspension was sequentially passed through a 70 µm nylon mesh cell strainer to obtain a purified single-cell suspension in phosphate-buffered saline (PBS). Additionally, spleens were excised from mice and subjected to mechanical dissociation by gently pressing them through a 70-μm nylon cell strainer utilizing a rubberized 1 ml syringe piston. This ensured gentle yet efficient disaggregation of the splenic tissue into a cellular suspension. Subsequently, the red blood cells were selectively lysed from the PBL, BM and splenic cell suspensions using an ammonium chloride-potassium (ACK) lysis buffer. This step facilitated the removal of erythrocytes, thereby enriching the final cell suspensions with leukocytes for further scientific analysis.

### Cell staining and flow cytometric analysis

BM, PBMC and spleen cell suspensions were meticulously collected and subsequently centrifuged at 300×g for 10 minutes at 4°C. The resulting cell pellet was gently resuspended in a solution of PBS supplemented with 0.2% bovine serum albumin (BSA), achieving a final cell concentration of 2 × 10^7 cells per milliliter. The cell suspensions were treated with an Fc Block agent (BD Biosciences, 564765) at a predetermined concentration, followed by a 15-minute incubation period at room temperature. Subsequently, the cells were incubated with a precisely formulated cocktail of fluorescently conjugated anti-mouse antibodies, inducing CD45-PE (BD, 553081), CD19-APC (BD, 561738) and S100A8-FITC (Novus Biologicals, NBP2-25269F). The incubation was carried out in the dark at 4°C for 30 minutes. Then the cells were washed twice with PBS supplemented with 0.2% BSA. Following the washes, the cells were resuspended and prepared for flow cytometric analysis. Flow cytometric data acquisition was performed on a suitable flow cytometer equipped with lasers (BD FACSymphony™ S6) and filters corresponding to the fluorochromes used in the antibody cocktail. Prior to data acquisition, compensation settings were adjusted using single-stained controls to correct for spectral overlap between the fluorochromes. The collected flow cytometric data were then analyzed using FlowJo software (FlowJo^TM^ v10).

### RNA extraction and RT‒qPCR analyses

Cell pellets were lysed in 1 mL of TRIZOL reagent (Life, 15596026) through vigorous mixing until a uniform lysate was achieved. To facilitate phase separation, 0.2 ml of chloroform was added to the lysate, which was then thoroughly mixed and allowed to equilibrate for 10 minutes at room temperature. Following this, centrifugation was performed at 12,000 × g for 15 minutes at 4°C to separate the phases. The upper aqueous layer, enriched in RNA, was carefully transferred to a clean tube. Subsequently, 0.5 mL of isopropanol was added to the aqueous phase, and the mixture was gently agitated and allowed to stand for an additional 10 minutes. The samples were then centrifuged at 12,000 × g for 10 minutes at 4°C to precipitate the RNA. The resulting RNA pellets were washed with 1 mL of 75% (v/v) ethanol by centrifuging at 7,500 × g for 5 minutes at 4°C. After removing the ethanol, the RNA pellets were air-dried for 5 to 10 minutes to ensure the removal of residual solvents. Finally, the dried RNA pellets were resuspended in RNase-free water to yield the purified RNA samples for downstream applications.

Reverse transcription was conducted utilizing SuperScript™ IV Reverse Transcriptase (Invitrogen, 18091050) as the enzymatic catalyst. Subsequently, quantitative polymerase chain reaction (qPCR) analyses were performed employing a TB-Green-based PCR kit (Takara, RR82WR) in conjunction with a Real-Time PCR system (Bio-Rad). The determination of fold changes in gene expression was achieved through the application of the comparative cycle threshold (ΔΔCT) method. As an internal control, β-actin served as the housekeeping gene to normalize data variations. The complete set of primer sequences utilized in this study was following: S100A8, AAATCACCATGCCCTCTACAAG and CCCACTTTTATCACCATCGCAA; S100A9, ATACTCTAGGAAGGAAGGACACC and TCCATGATGTCATTTATGAGGGC; β-actin, CCCTGAAGTACCCCATTGAAC and CCTTTCACGGTTGGCCTTAG.

### Lymphocyte cultures

Human BM aspirates (5-10 mL), collected from three healthy volunteers aged between 53 and 66 years following written informed consent, underwent BMMC isolation via density gradient centrifugation utilizing Ficoll-Paque (1.078 g/mL) as the separating medium. This process entailed overlaying the diluted BM fraction onto Ficoll-Paque and subsequently centrifuging the specimen at 600×g for 30 minutes at ambient temperature. Following centrifugation, BMMCs were carefully aspirated from the buffy coat interface and washed twice with PBS. Subsequently, the procured BMMCs were divided into two distinct fractions. One fraction was promptly introduced into the culture medium. The second fraction was subjected to negative selection employing the BD IMag™ Human B Lymphocyte Enrichment Set (BD, 558007), adhering strictly to the manufacturer’s protocol, to isolate Pan-B cells. The resulting Pan-B cell population was resuspended in culture medium and plated onto culture dishes, maintained under standard conditions of 37 °C and 5% CO2 for 96 hours in a serum-free environment. Furthermore, the B-cell line ARH-77 was employed as a comparative reference to ensure rigorous experimental control. Additionally, a parallel experiment involving a complete culture medium devoid of any cellular components was conducted under identical environmental conditions, serving as a blank control to mitigate potential confounding factors.

### ELISA

The concentrations of S100A8/S100A9 in the culture medium were quantitatively determined utilizing a Human Calprotectin ELISA Kit (S100A8/S100A9) (Abcam, ab267628), adhering strictly to the manufacturer’s prescribed protocol. Briefly, predetermined standards and experimental samples were dispensed into individual wells of a 96-well plate pre-coated with primary antibodies specific to S100A8/A9. After washing steps, a biotinylated anti-calprotectin antibody was introduced to the wells. Following another round of washing, HRP-conjugated streptavidin was added, and the plate was incubated for 45 minutes. Thereafter, the TMB substrate solution was added to each well, allowing a colorimetric reaction. Optical densities at 450 nm were recorded using the VersaMax microplate reader, and the concentrations of the S100A8/S100A9 heterodimer were subsequently derived from the absorbance values by interpolation against a standard curve.

## Data availability

The raw sequence data reported in this paper have been deposited in the Genome Sequence Archive (Genomics, Proteomics & Bioinformatics 2021) in National Genomics Data Center (Nucleic Acids Res 2022), China National Center for Bioinformation / Beijing Institute of Genomics, Chinese Academy of Sciences (GSA-Human: HRA006503) that are publicly accessible at https://ngdc.cncb.ac.cn/gsa-human.

## Code availability

The code generated during this study to analyze the single-cell datasets is available through GitHub at https://github.com/Xiujia-Yang/Review_of_B_Cell.

## Acknowledgements

Funding

This study was supported by the National Key R&D Program of China (2022YFF1203100 to Z.Z.), the Guangdong Basic and Applied Basic Research Foundation (2022B1515230005 to Z.Z.), the National Natural Science Foundation of China (31771479, 81991511, 81991510 and 32370593 to Z.Z.; 82170731 and 81470974 to W.W.; 32300537 to X.Y.), and the High-level Hospital Construction Project of Guangdong Province (DFJH201908 to W.W.).

## Author information

Authors and Affiliations

Guangdong Cardiovascular Institute, Guangdong Provincial People’s Hospital, Guangdong Academy of Medical Sciences, Guangzhou 510080, China

Xiujia Yang

Center for Precision Medicine, Medical Research Institute, Guangdong Provincial People’s Hospital (Guangdong Academy of Medical Sciences), Southern Medical University, Guangzhou 510080, China

Xiujia Yang, Haipei Tang, Chunhong Lan, Sen Chen, Huikun Zeng, Haoyu Wu & Zhenhai Zhang

Guangdong-Hong Kong Joint Laboratory on Immunological and Genetic Kidney Diseases, Guangdong Provincial People’s Hospital (Guangdong Academy of Medical Sciences), Southern Medical University, Guangzhou 510080, China

Xiujia Yang, Haipei Tang, Chunhong Lan, Huikun Zeng & Zhenhai Zhang

Department of Bioinformatics, School of Basic Medical Sciences, Southern Medical University, Guangzhou 510515, China

Chunhong Lan, Sen Chen, Haoyu Wu & Zhenhai Zhang

Division of Nephrology, Guangdong Provincial People’s Hospital (Guangdong Academy of Medical Sciences), Southern Medical University, Guangzhou, 510080, China

Weiting He, Huikun Zeng, Danfeng Liu & Wenjian Wang

Key Laboratory of Mental Health of the Ministry of Education, Guangdong-Hong Kong-Macao Greater Bay Area Center for Brain Science and Brain-Inspired Intelligence, Southern Medical University, Guangzhou 510515, China

Zhenhai Zhang

## Author contributions

X.Y., W.W. and Z.Z. conceived, designed and directed the study. W. H. and D. L. collected human bone marrow and matched peripheral blood samples. X.Y., C.L., S.C., H.Z. and H.W. performed data analysis and visualization. H.T. performed single-cell RNA sequencing, RT-qPCR, cell culture and ELISA. X.Y. and H.T. wrote the original manuscript. X.Y., W.W. and Z.Z. reviewed and edited the original manuscript.

Corresponding authors

Correspondence to Xiujia Yang, Wenjian Wang or Zhenhai Zhang.

Ethics declarations

Competing interests

The authors declare no competing interests.

---

**Supplementary figure 1.**
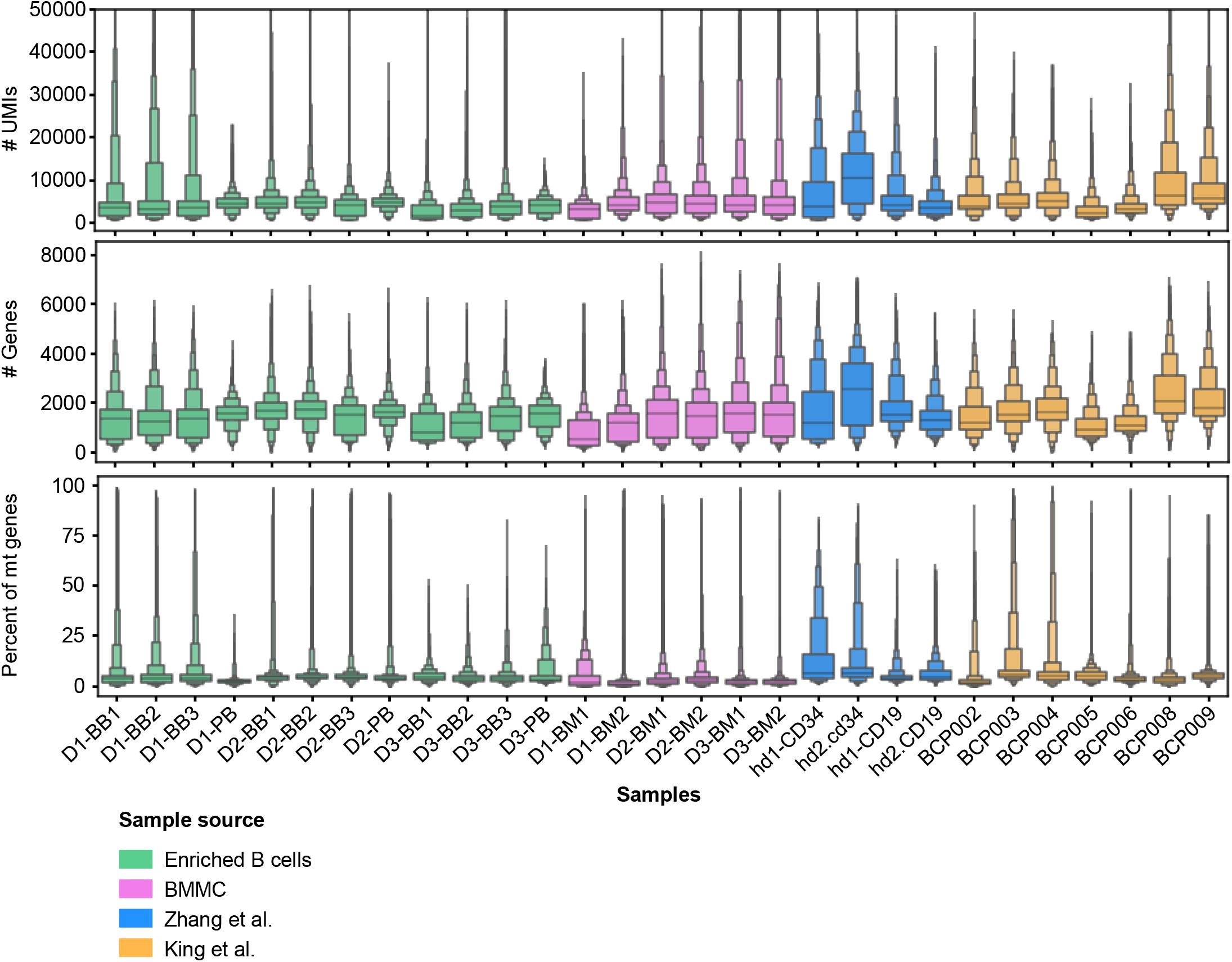
**Quality control metrics for scRNA-seq samples from this study (enriched B and BMMC) and two external studies**. Notably, the scRNA-seq samples from Zhu et al. were not included for evaluation here due to the unavailability of the raw expression matrices.

**Supplementary figure 2.**
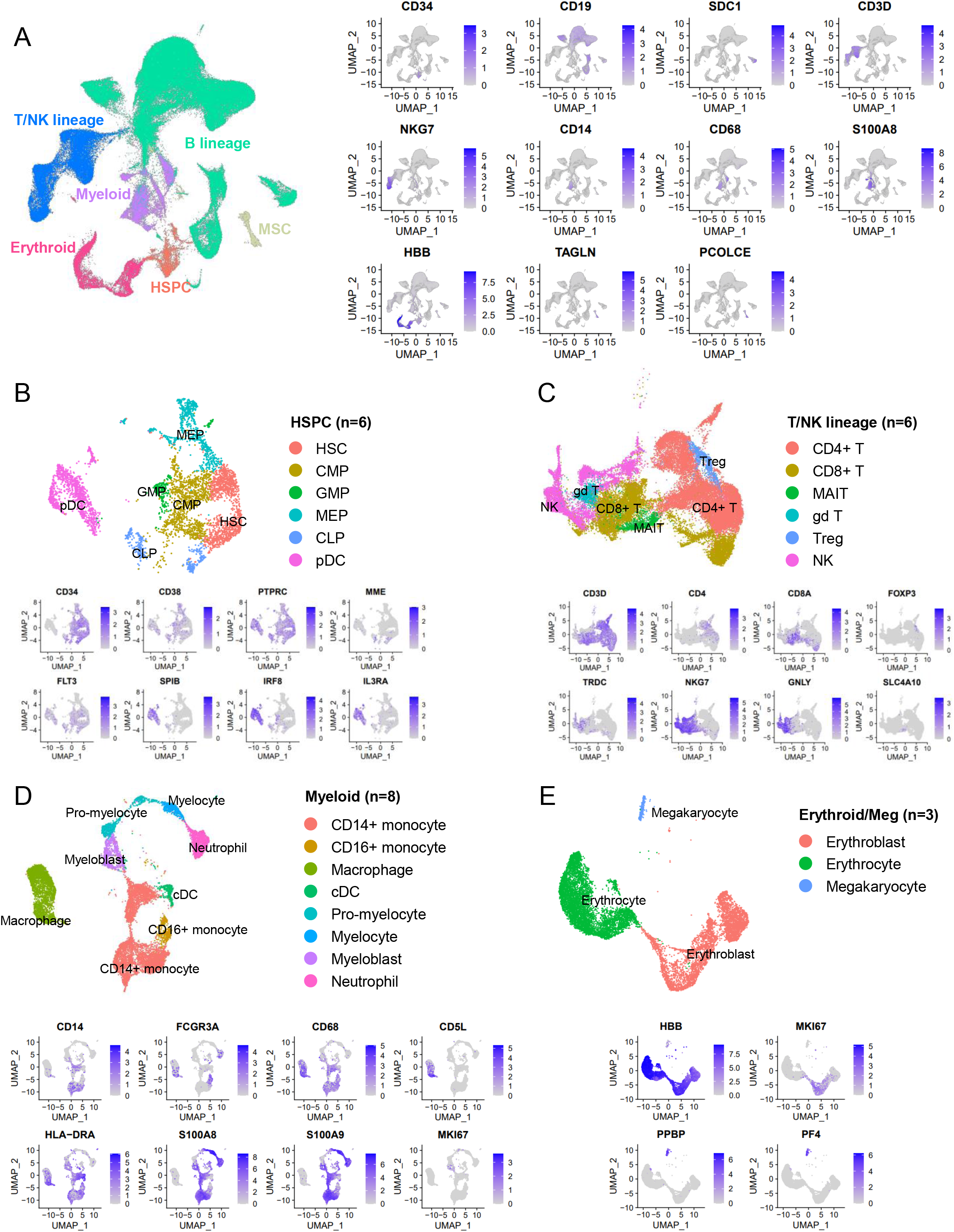
Cell population identification based on single-cell transcriptomic profiling data. Cell populations were identified by two sequential clusterings, the preliminary clustering **(A)** and the secondary clusterings **(B-E)**. In the preliminary cell clustering, cells of different lineages were distinguished (A), whereas in the secondary clusterings cell populations within a typical lineage (HSPC (B), B lineage (Fig. 1D), T/NK lineage (C), myeloid (D) and erythroid/meg (E)) were identified unbiasedly. Cells of different lineages or populations were denoted by different colors in the UMAP. The expression profile of marker genes used to distinguish lineages or cell populations was also provided. HSPC, hematopoietic stem and progenitor cell.

**Supplementary figure 3.**
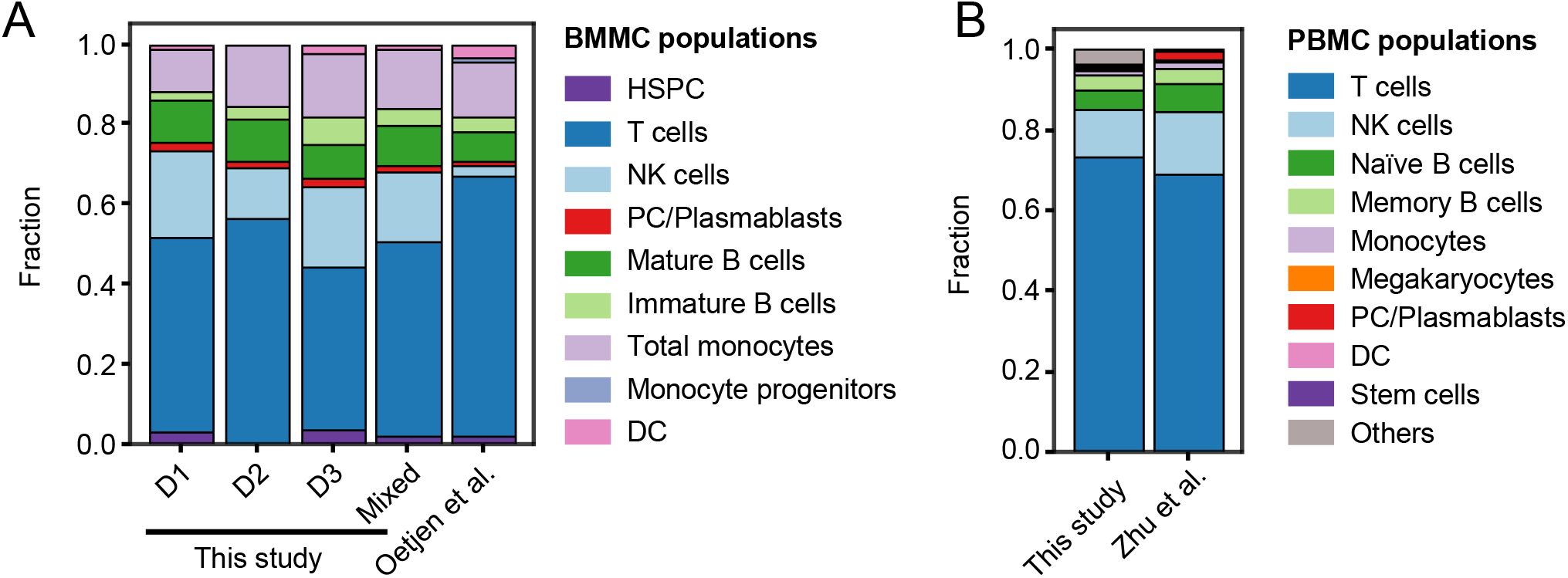
Cell population frequency comparison between this study and external references for CD45+ BMMCs (A) and PBMCs (B). Notably, only CD45+ cell populations (excluding erythrocytes and megakaryocytes) were considered here for comparison with data from Oetjen et al. HSPC, hematopoietic stem and progenitor cells; PC, plasma cell; DC, dendritic cell.

**Supplementary figure 4.**
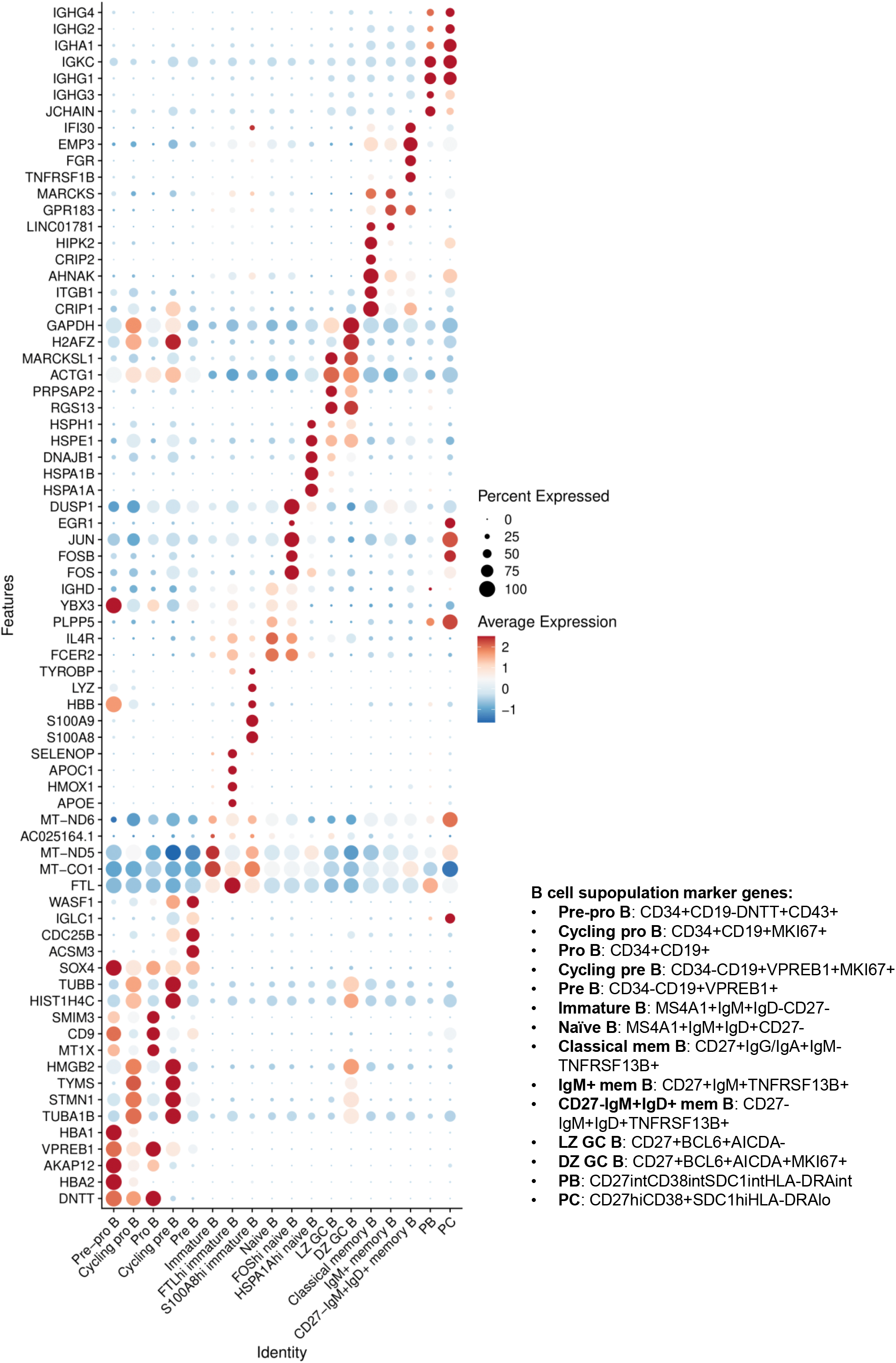
Expression of top 5 highly variable genes across 18 B cell subpopulations and a list of marker genes assisting the annotation of B cell subpopulations in this study.

**Supplementary figure 5.**
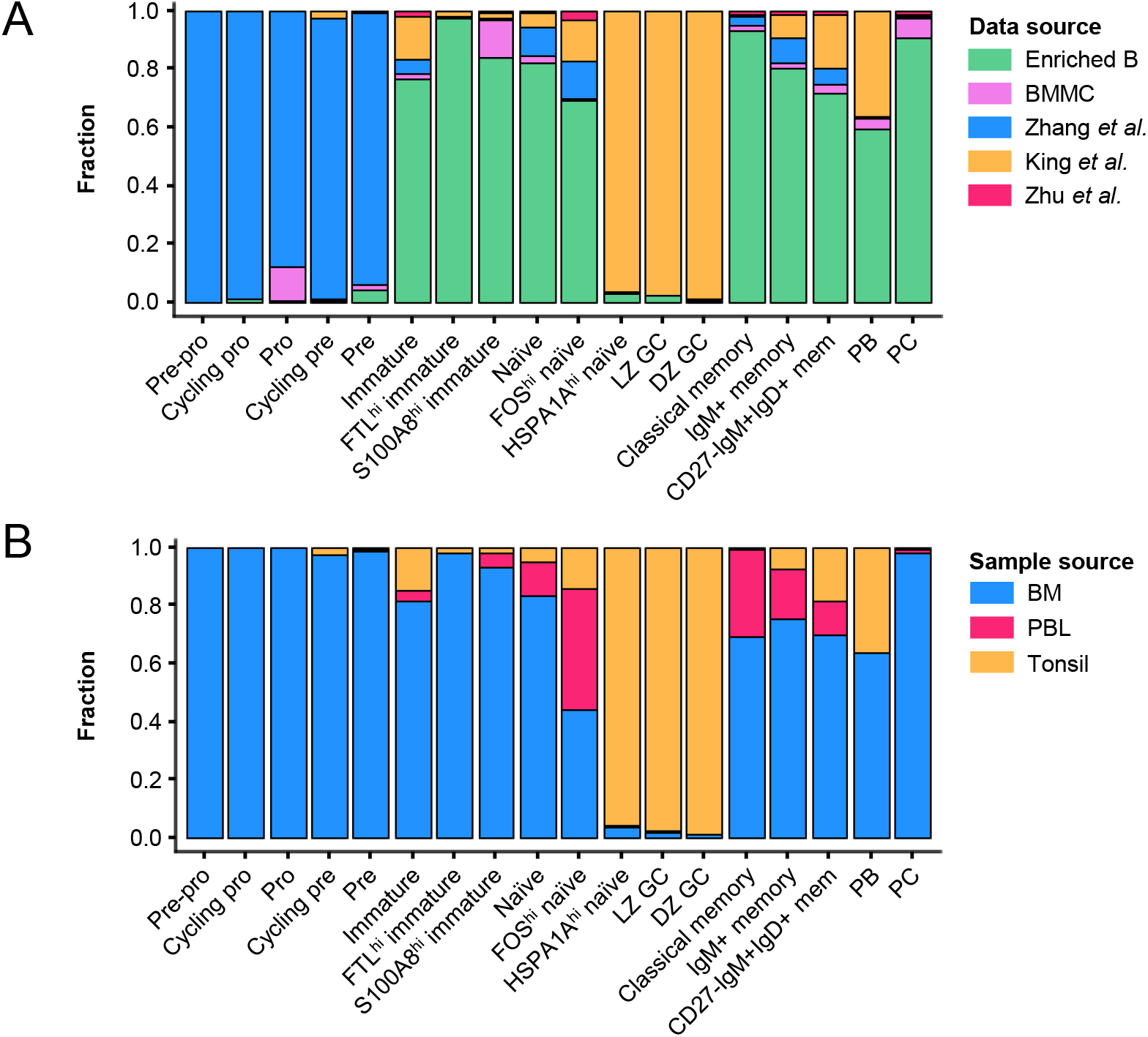
The data (A) and sample (B) source composition for 18 identified B cell clusters.

**Supplementary figure 6.**
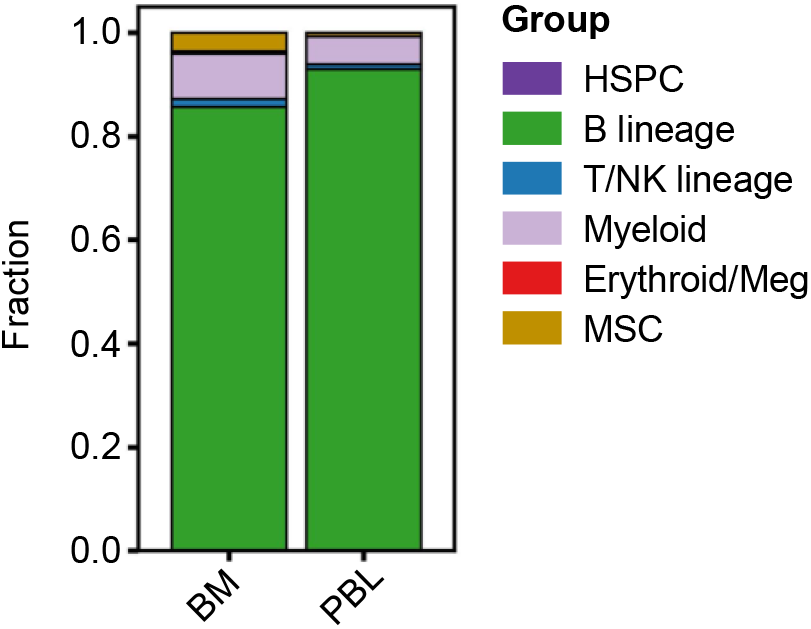
**Cell group composition of enriched BM and PBL B cells**. HSPC, hematopoietic stem and progenitor cells; MSC, mesenchymal stem cells.

**Supplementary figure 7.**
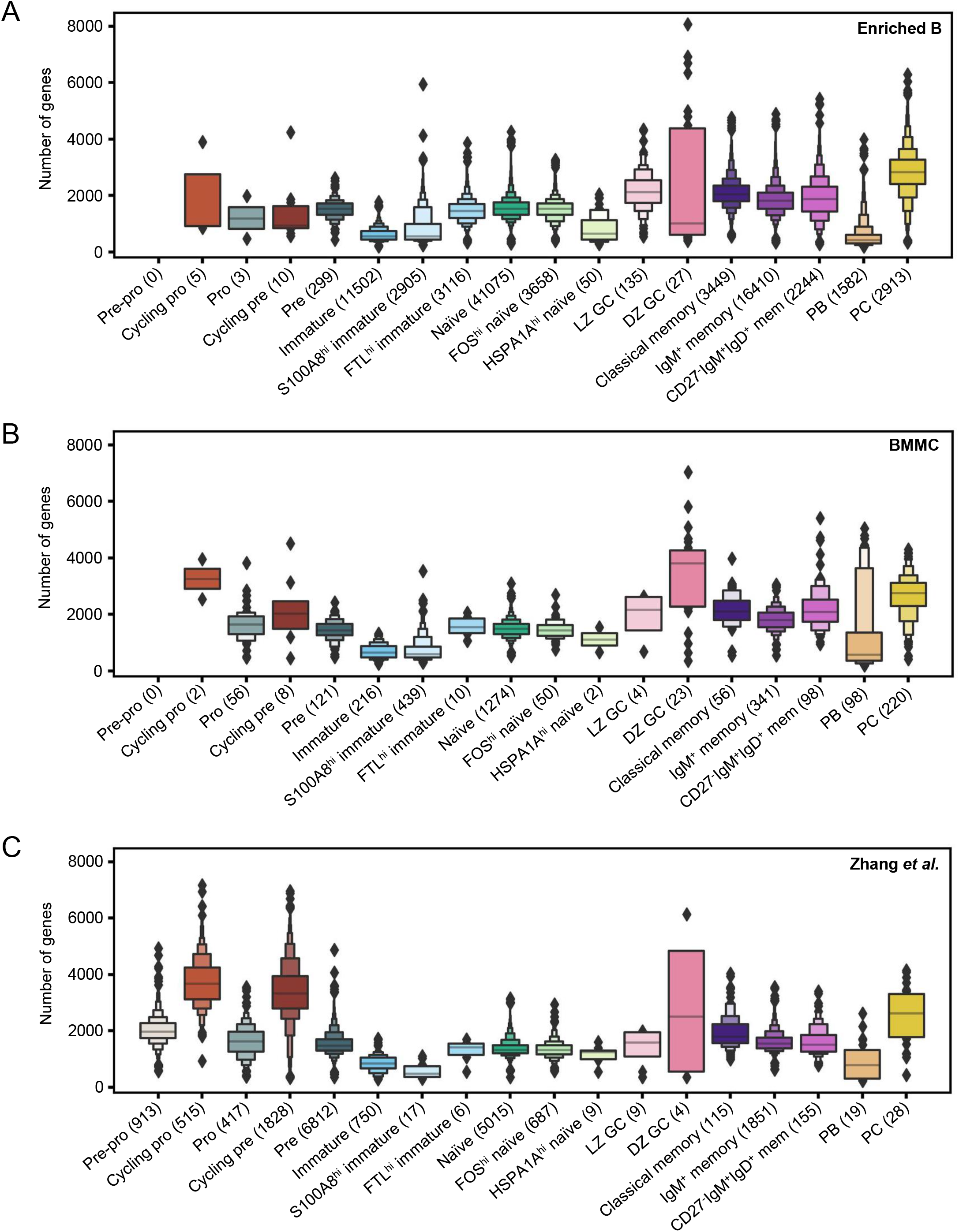
Distribution of number of genes captured by scRNA-seq across the 18 identified B cell subpopulations. These gene number distributions from individual projects or sample type (A, enriched B; B, BMMC; C, Zhang *et al.*) reproduce the trend shown in Fig. 2A.

**Supplementary figure 8.**
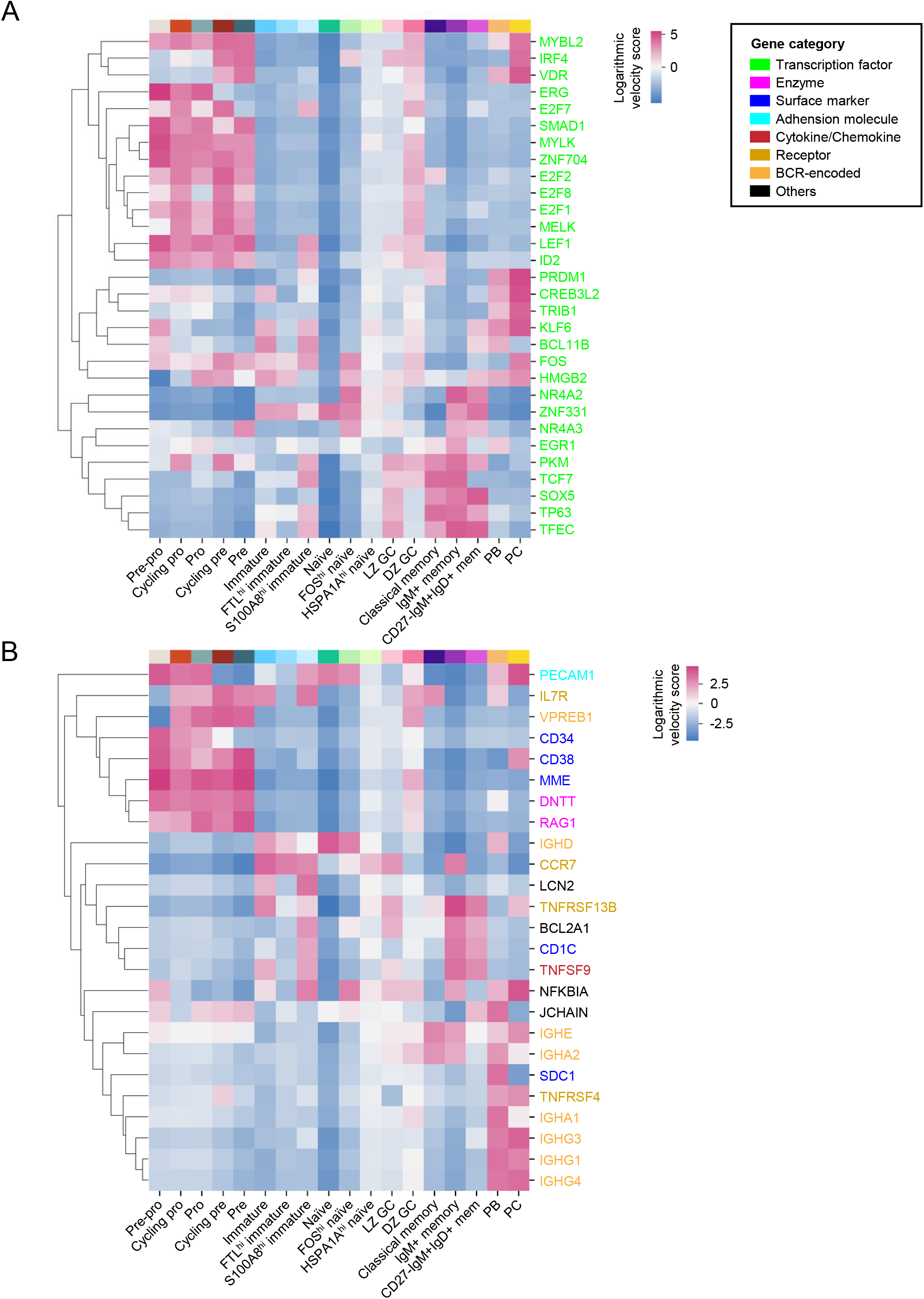
Velocity score profile for a selected list of genes (A, transcription factor; B, representative molecules) across B cell subpopulations (with GC B cells unavailable).

**Supplementary figure 9.**
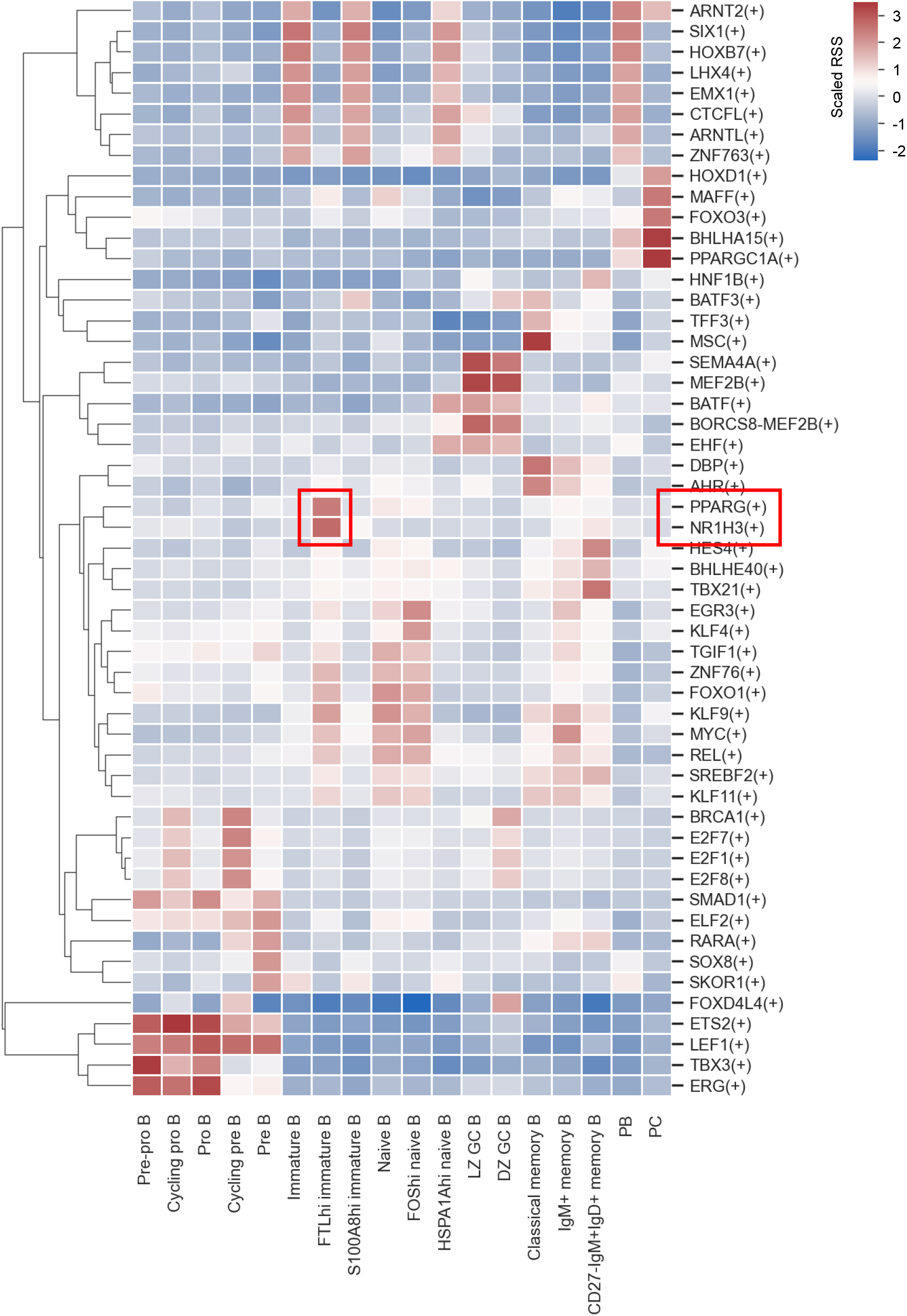
Scaled regulon specificity score (RSS) across B cell subpopulations. Regulons are hiearachically clustered. The two regulons most specific for FTL^hi^ immature B cell subpopulation were marked in the heatmap.

**Supplementary figure 10.**
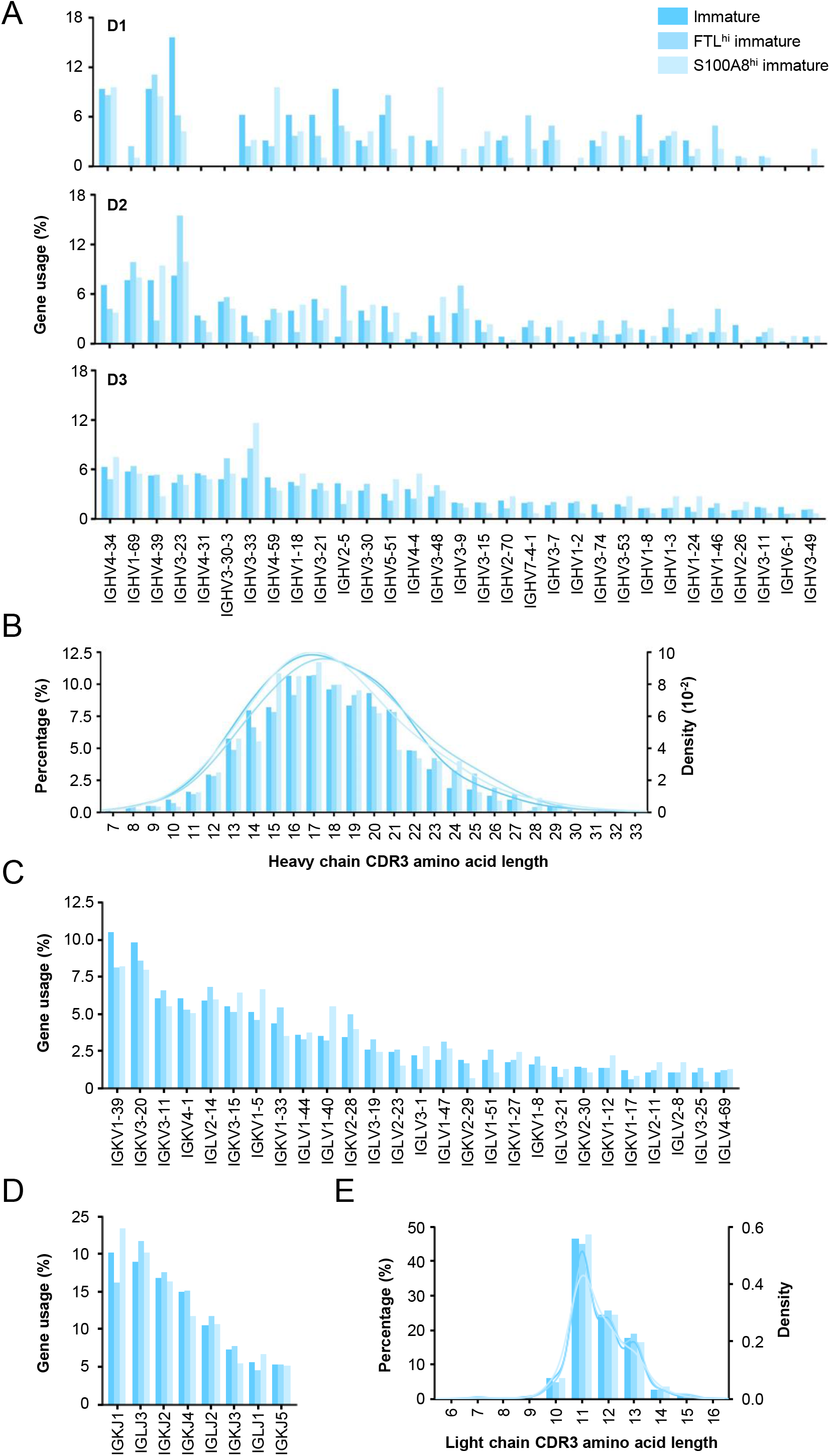
**BCR repertoire characteristics for immature B cell subpopulations**. **(A)** Donor-specific BCR heavy chain variable gene usage across 3 immature B cell subpopulations. **(B)** Heavy chain CDR3 length distribution across 3 immature B cell subpopulations. The paired smooth curves are distributions fitted from kernel density estimate. **(C-D)** BCR light chain variable (B) and joining (C) gene usage as determined by scVDJ-seq data across 3 immature B cell subpopulations. For variable genes, only those with a usage frequency of at least 1% were shown. **(E)** Light chain CDR3 length distribution. The paired smooth curves are distributions fitted from kernel density estimate.

**Supplementary figure 11.**
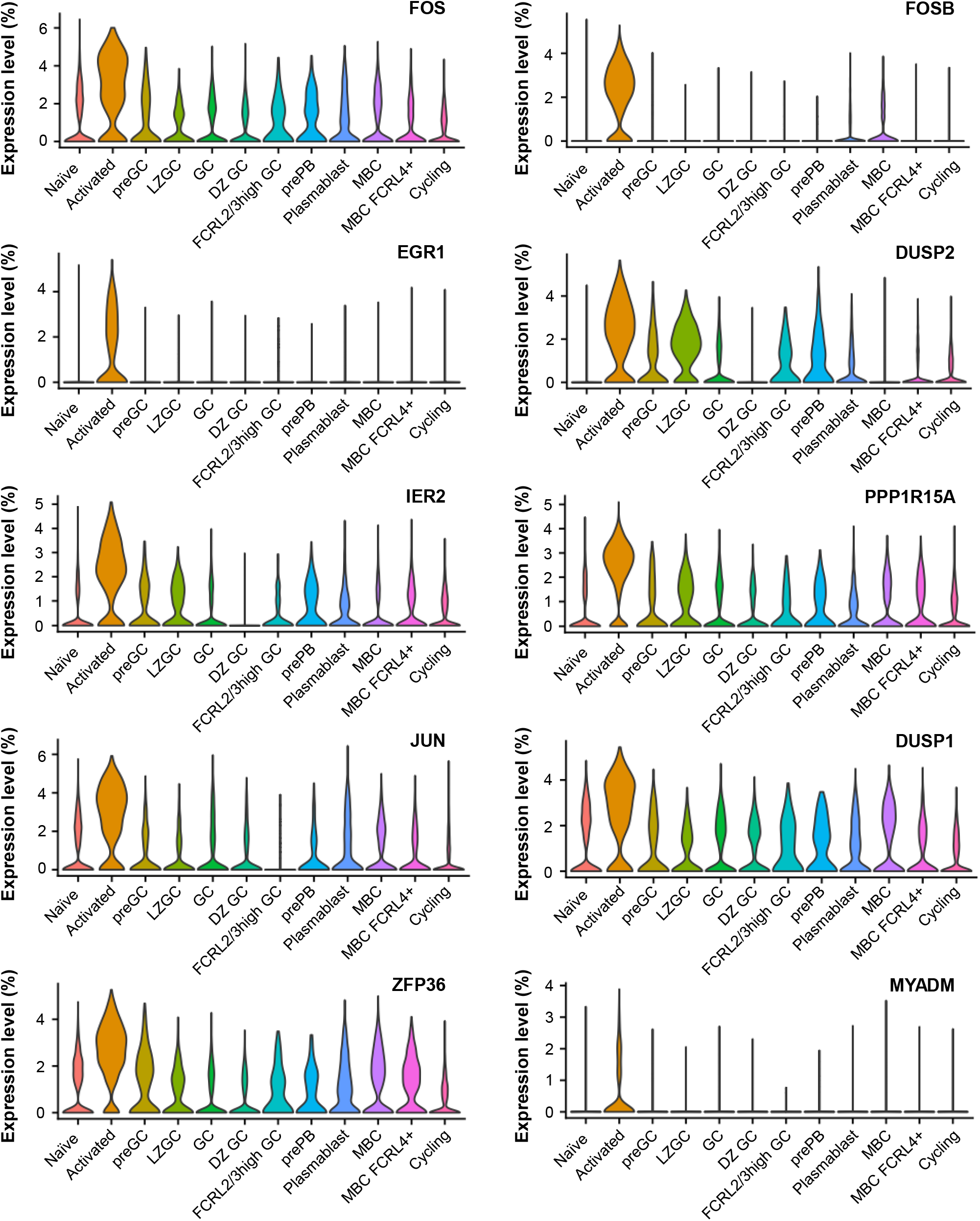
Expression of the top 10 genes of FOS^hi^ naïve B cells in cell types of King et al dataset. This gene expression pattern confirmed the ‘activated’ phenotype of FOS^hi^ naïve B cells.

**Supplementary figure 12.**
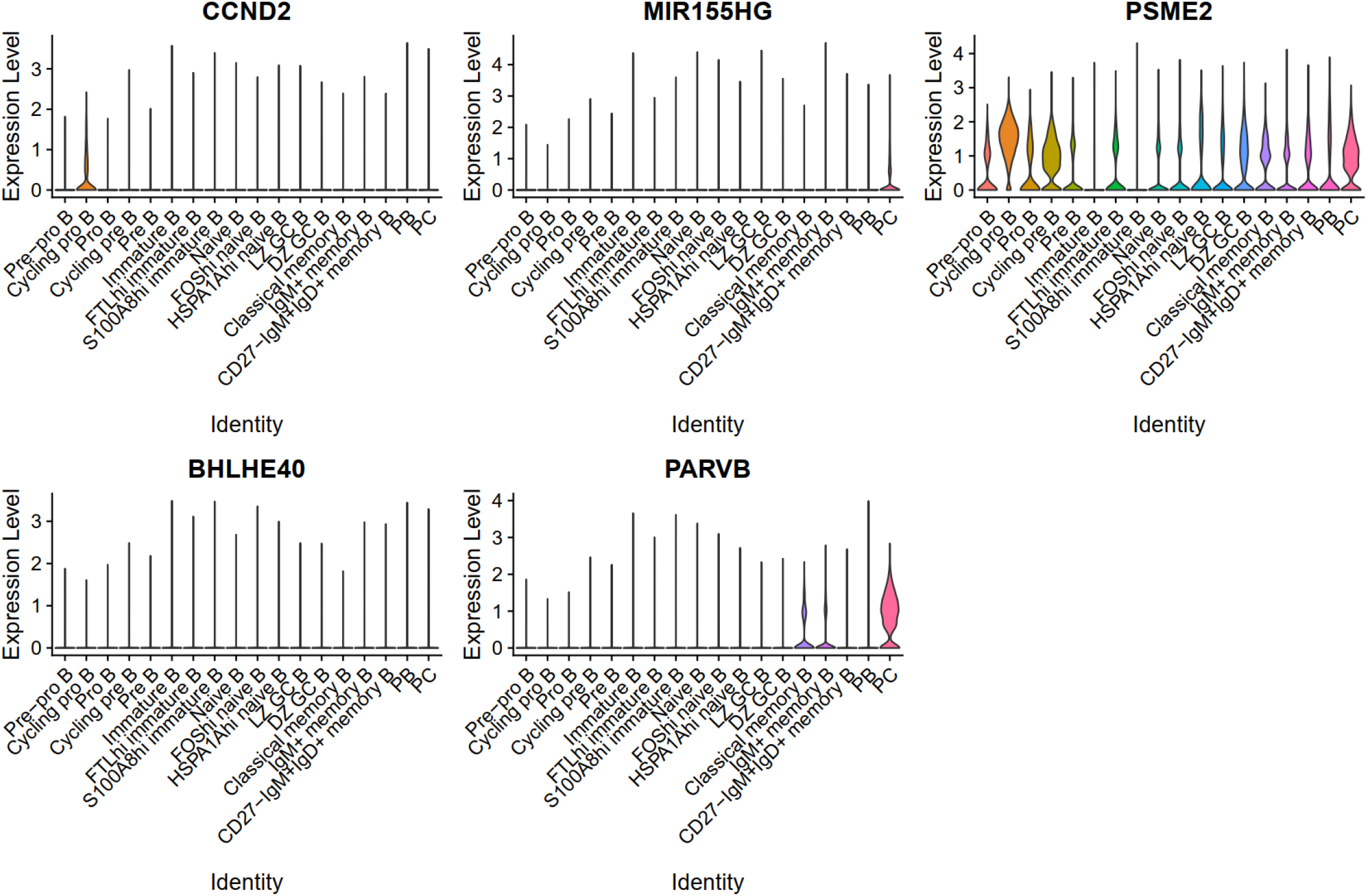
**Expression of the top 5 genes for the preGC subset of King *et al* dataset in HSPA1A^hi^ naïve B cells in this study**. This gene expression pattern doesn’t support that the HSPA1A^hi^ subpopulation is similar to the preGC subset.

**Supplementary figure 13.**
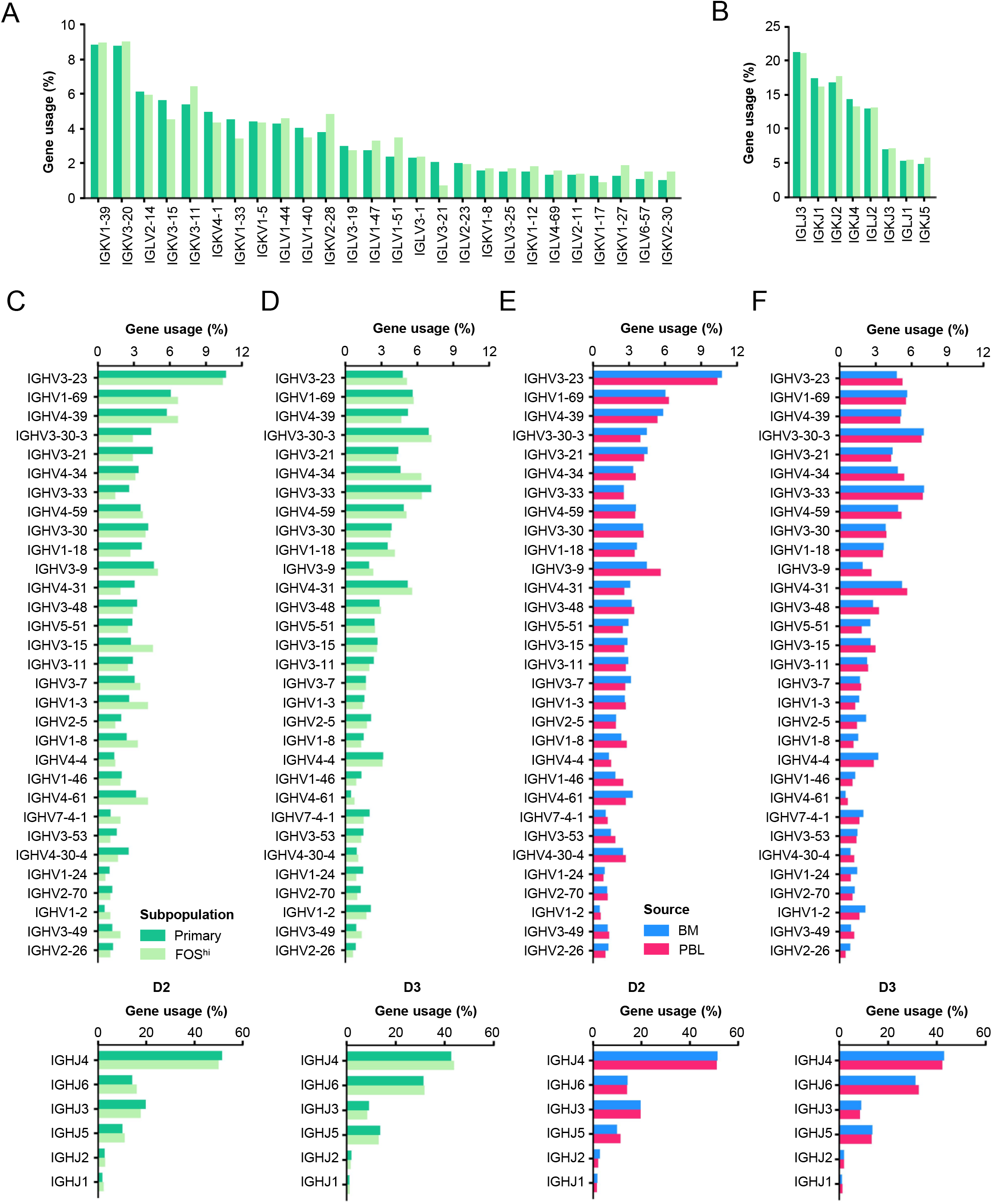
BCR variable (V) and joining (J) gene usage profile for primary and FOS^hi^ naïve subpopulations. (A-B) BCR light chain variable (A) and joining (B) gene usage comparison between subpopulations. **(C-D)** BCR heavy chain V and J gene usage comparison between subpopulations as split by donor (C for D2 and D for D3). **(E-F)** BCR heavy chain V and J gene usage comparison between two sample source as split by donor (primary and FOS^hi^ naïve subpopulations were merged, C for D2 and D for D3).

**Supplementary figure 14.**
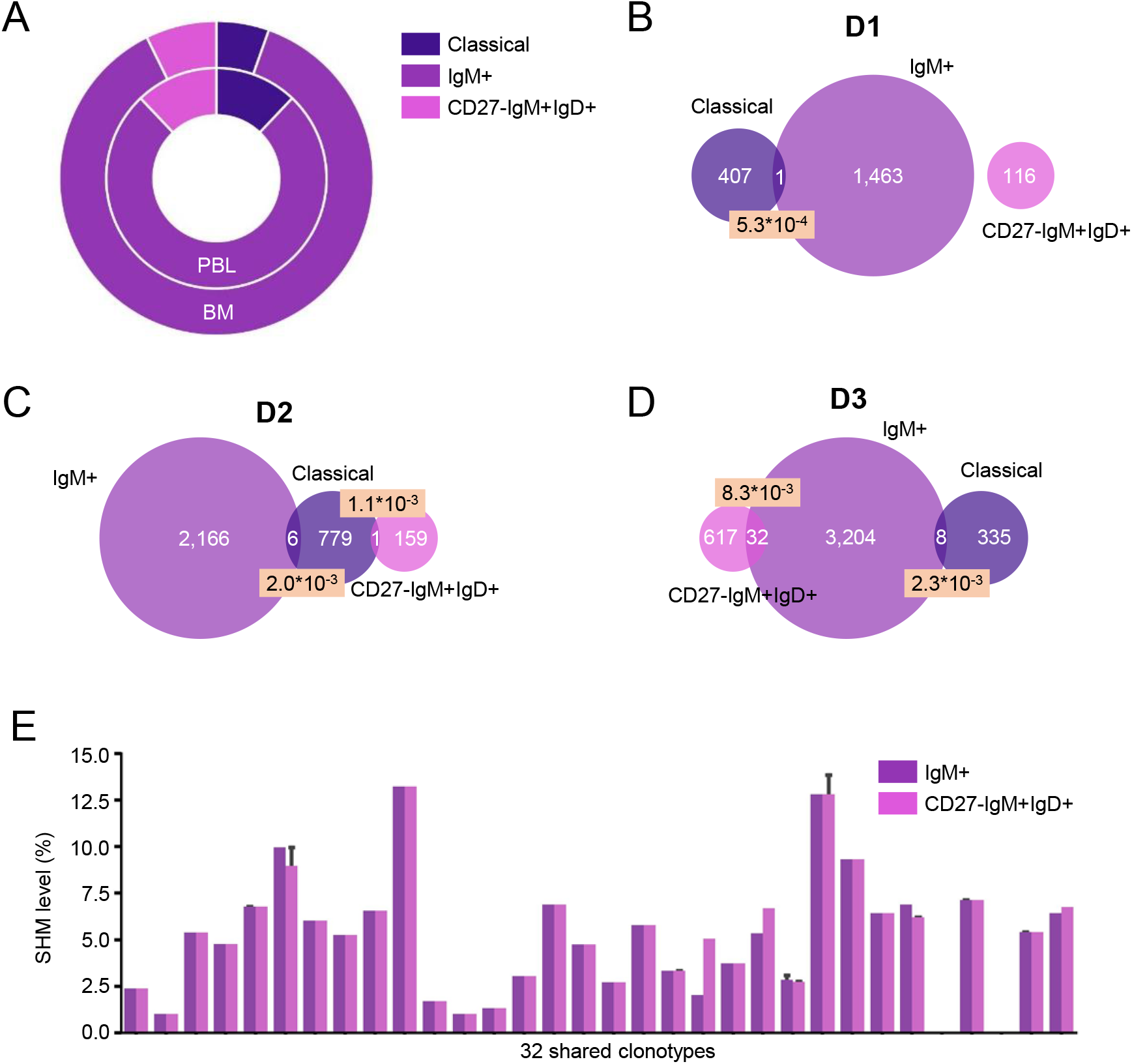
Characterization of memory B cell subpopulations. **(A)** The higher percentage of IgM/CD27- IgM+IgD+ compartment in memory B cells in BM (Zhang *et al*) compared to PBL (Zhu *et al*) was also confirmed by the external dataset. **(B-D)** Venn diagram showing clonotype sharing between classical, IgM+ and CD27-IgM+IgD+ memory B cell subpopulations for individual donors. The numbers with a pink background indicate the clonotype similarity as measured by the Jaccard similarity index. **(E)** Heavy chain variable gene SHM level comparison between IgM+ and CD27-IgM+IgD+ subpopulations for each of 32 shared clonotypes.

**Supplementary figure 15.**
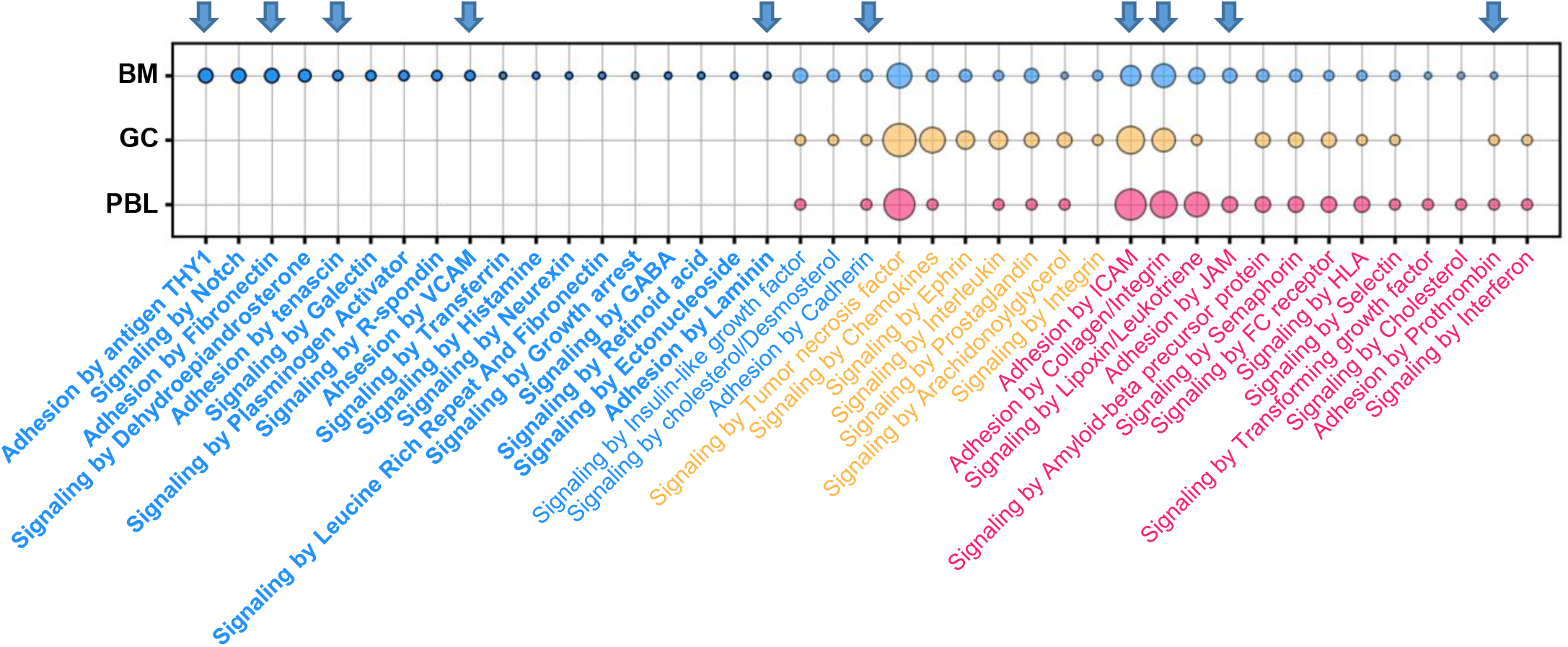
**Comparison of the predicted signaling pathways between different sources**. The dotplot corresponds to the venn diagrams (Fig. 7D-F). The dot size is proportional to the frequency of a signaling pathway. The label color indicates the source for which a signaling pathway frequency is the highest. Dots with a opaque facecolor indicate source-specific signaling pathways (with bold labels) whereas dots with a transparent facecolor indicate signaling pathways shared between sources. The upper arrows mark the 10 predicted adhesion signaling pathways.

**Supplementary figure 16.**
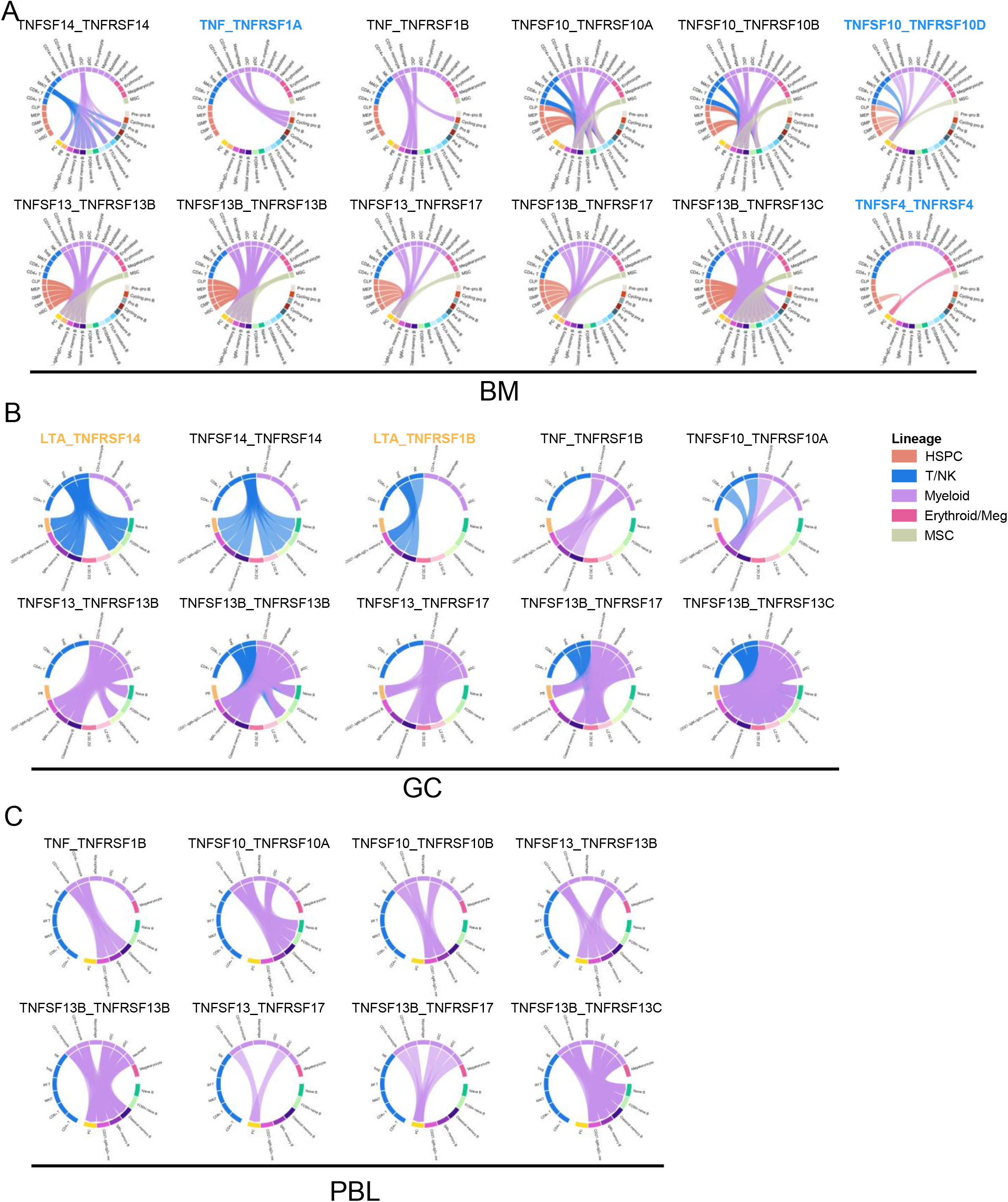
Signaling by tumor necrosis family between non-B and B cells in BM (A), GC (B) and PBL (C) data as demonstrated by the chord diagram. Tissue-specific CCIs are highlighted. Non-B cells are colored according to their lineage origin. All CCIs between non-B (expressing ligands) and B cells (expressing receptors) mediated by TNF family are represented by the colored arrows.

**Supplementary figure 17.**
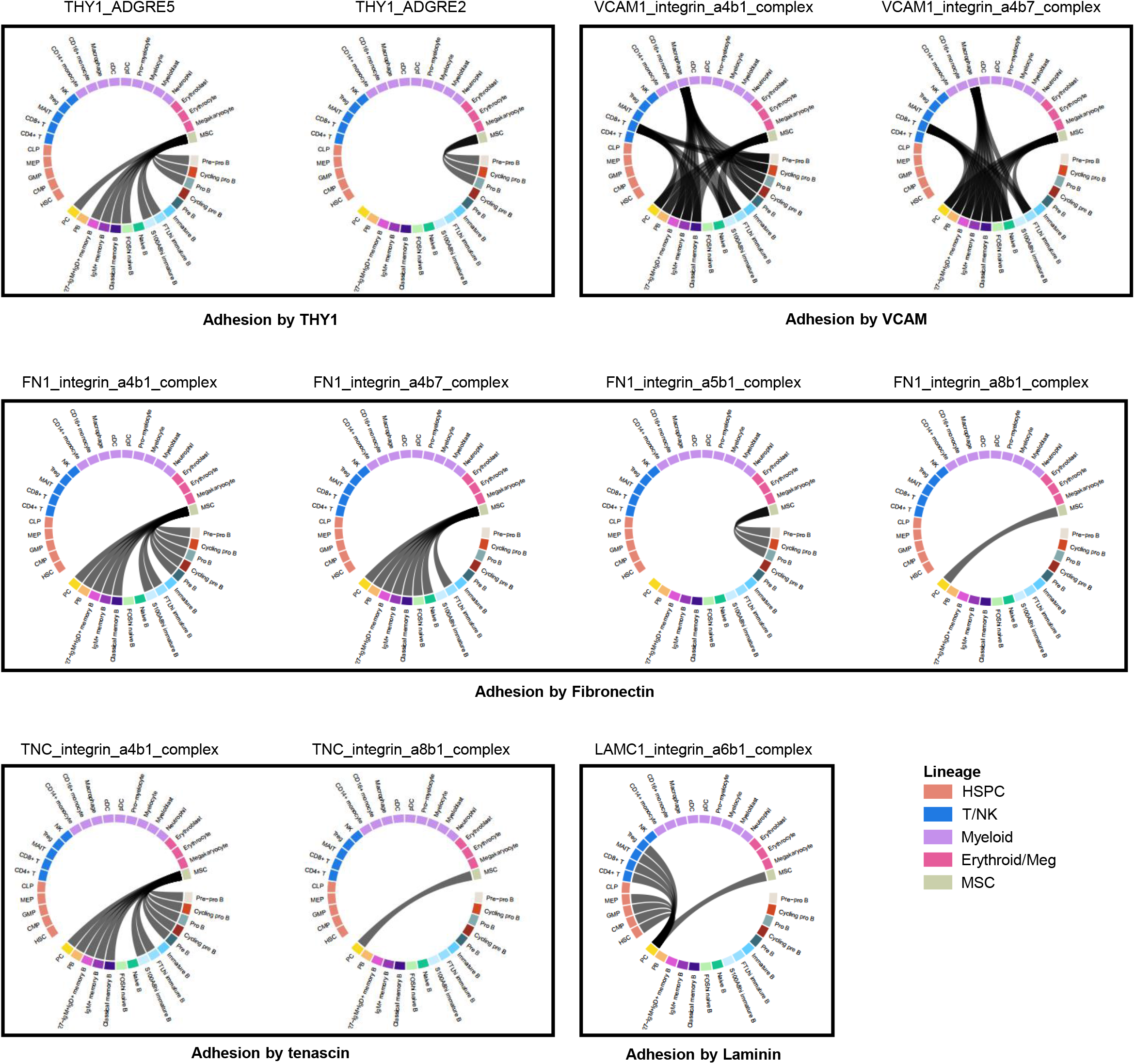
BM-specific adhesion categories as demonstrated by the chord diagram. The adhesion CCIs between non-B and B cells are represented by the undirected ribbons. Non-B cells are colored according to their lineage origin.

**Supplementary figure 18.**
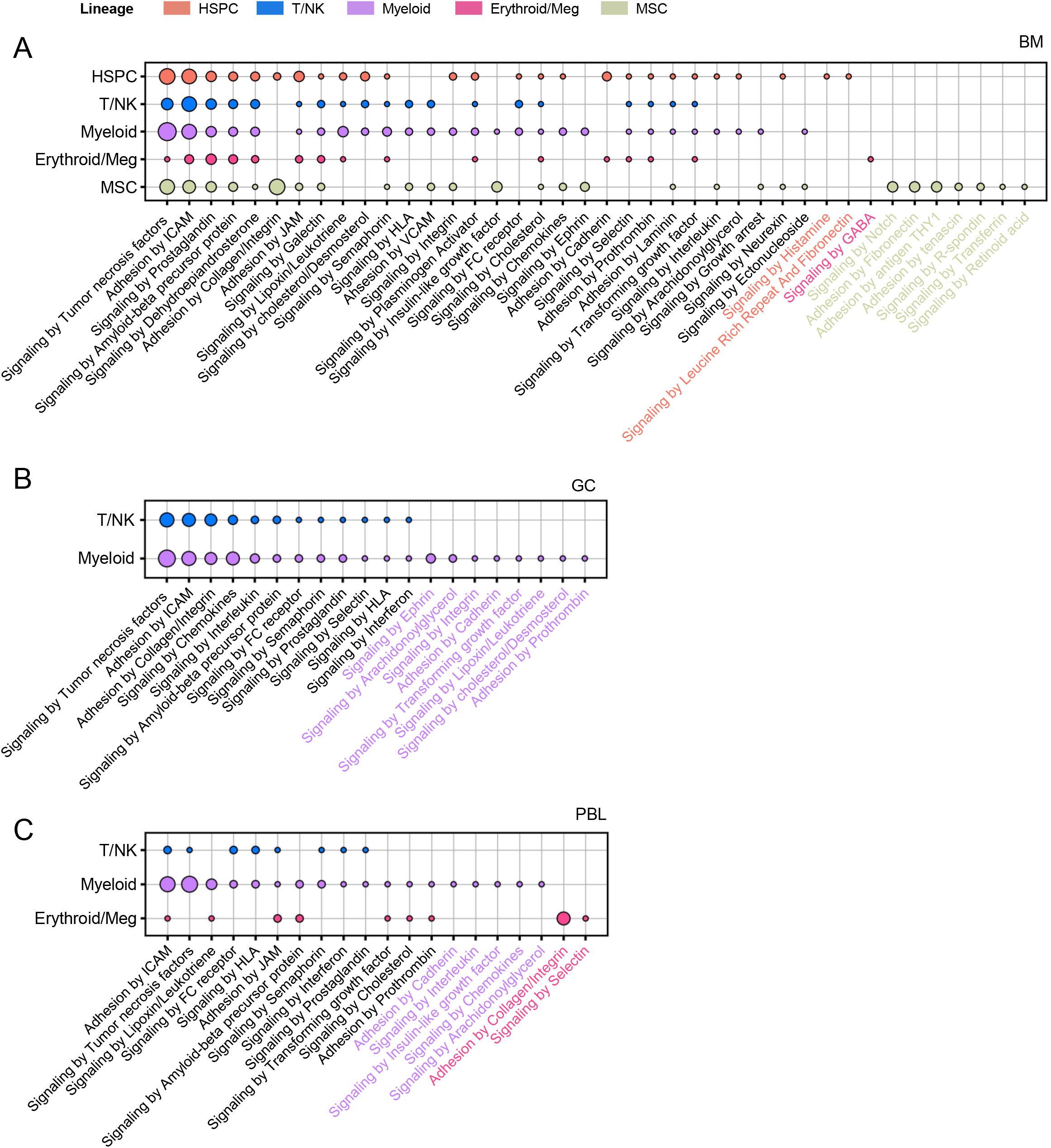
Lineage-specific signaling for non-B cells as demonstrated by dot arrays. These dot arrays show the lineage-specific signaling in BM **(A)**, GC **(B)** and PBL **(C)**, respectively. All dots and lineage-specific signaling are colored according to the lineage. The circle size is proportional to the number of CCIs.

**Supplementary figure 19.**
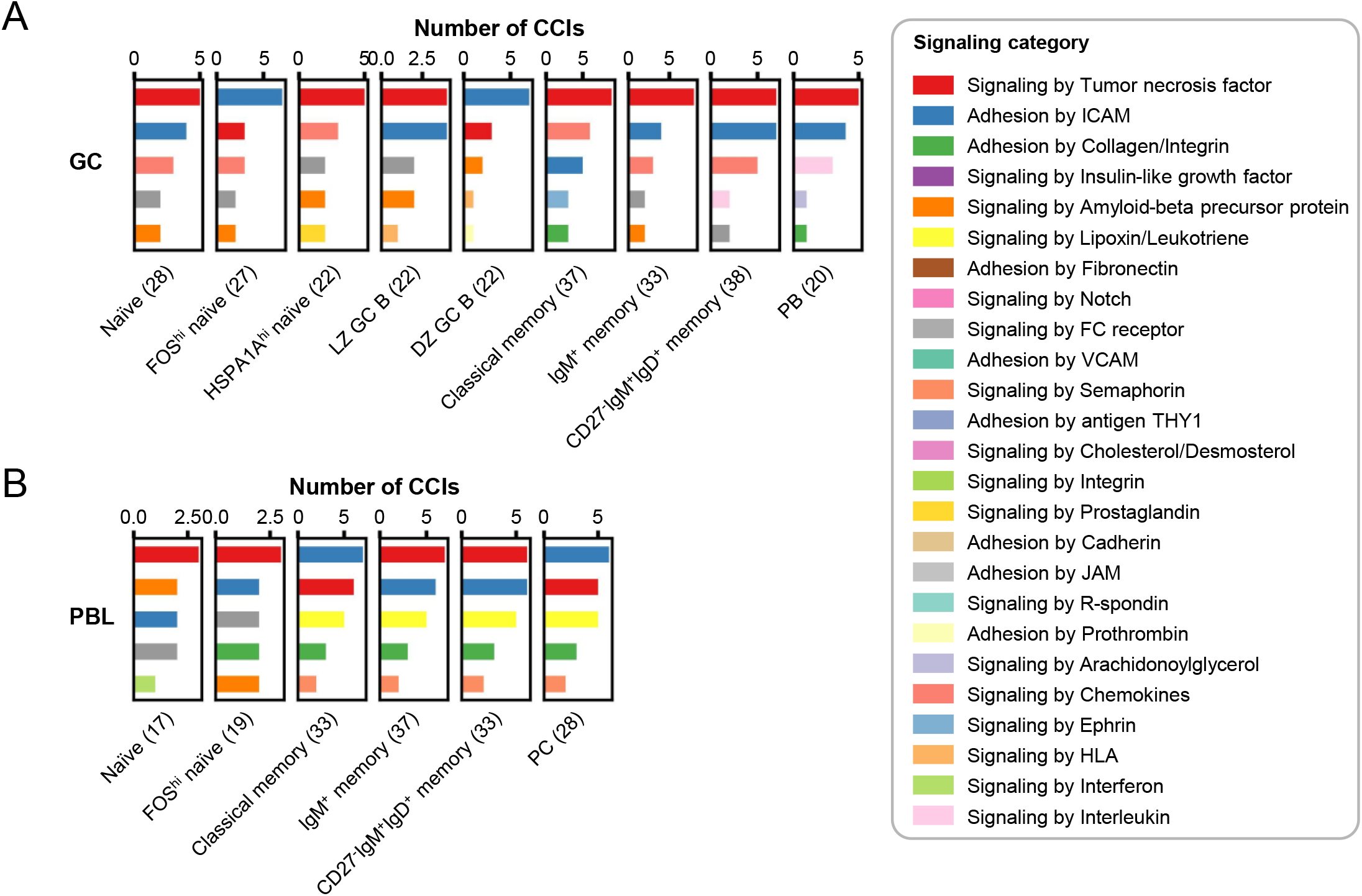
Top 5 signaling categories predicted by CellPhoneDB for B cell subpopulations in GC (A) and PBL (B). Signaling category is color-coded. In the parentheses is the total number of CCIs predicted for a typical B cell subpopulation.

**Supplementary figure 20.**
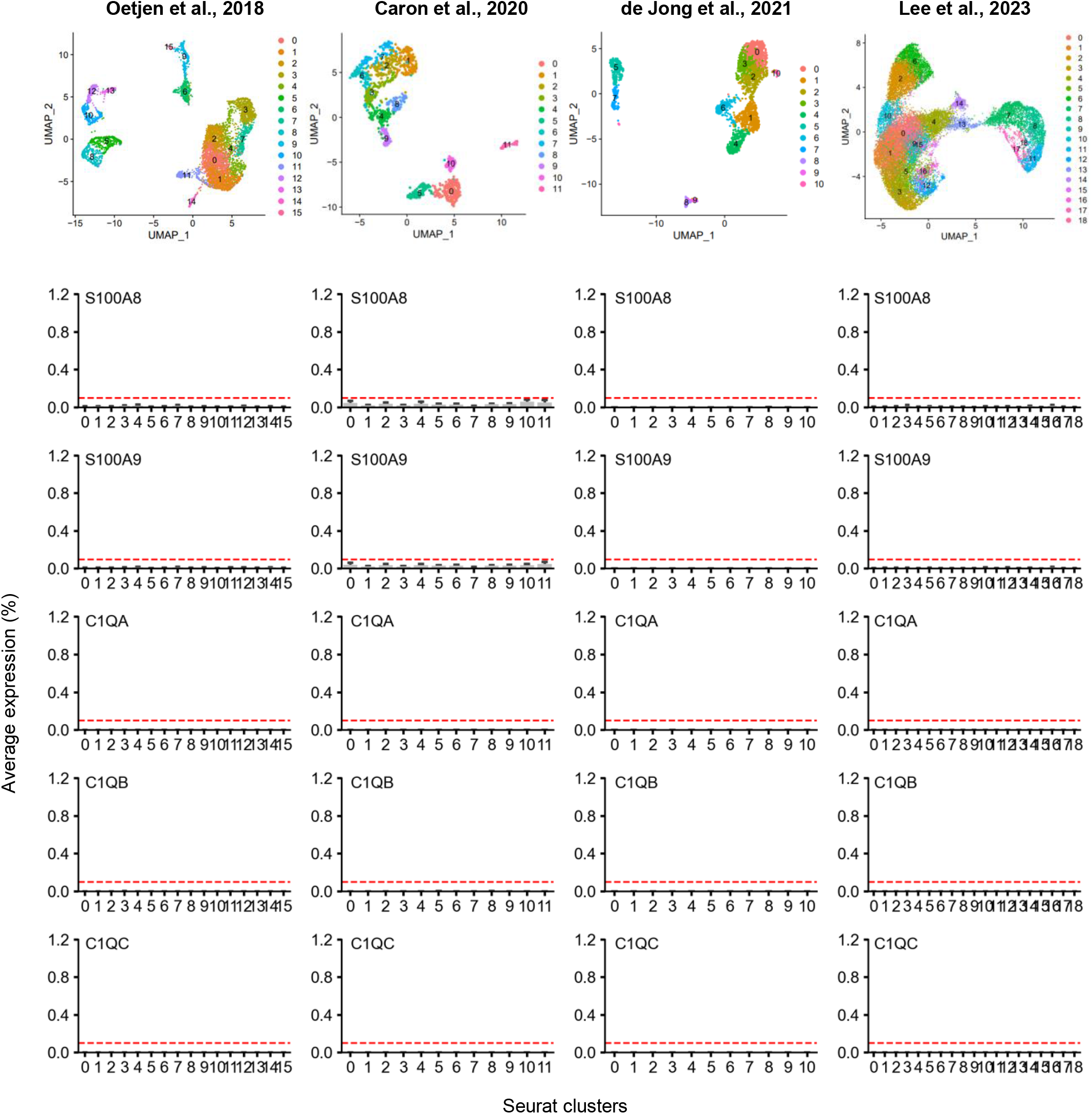
UMAP of B cell clusters with the resolution setting to 1.0 and the expression of C1QA, C1QB, C1QC, S100A8 and S100A9 across B cell clusters in four external datasets. The red dash lines mark an average expression level of 0.1%. The four external single-cell datasets showed an absence of BM S100A8^hi^ B cell subset (less than 01.%) in young subjects.

**Supplementary figure 21.**
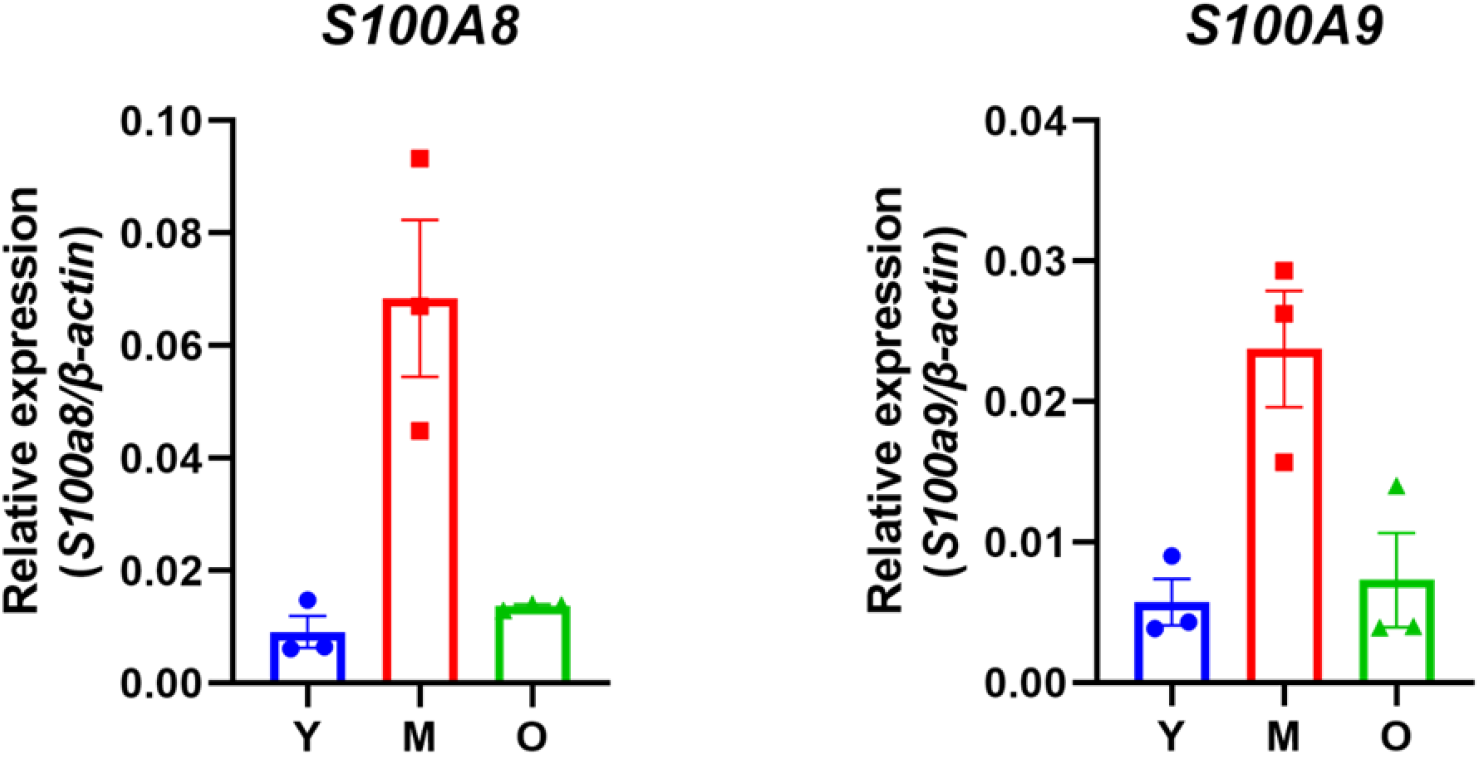
Age-associated expression of S100A8/A9 in bone marrow B cells of C57BL/6 mice as measured by qPCR. Y, young (3m); M, middle age (12m); O,1 old (22m).

**Supplementary figure 22.**
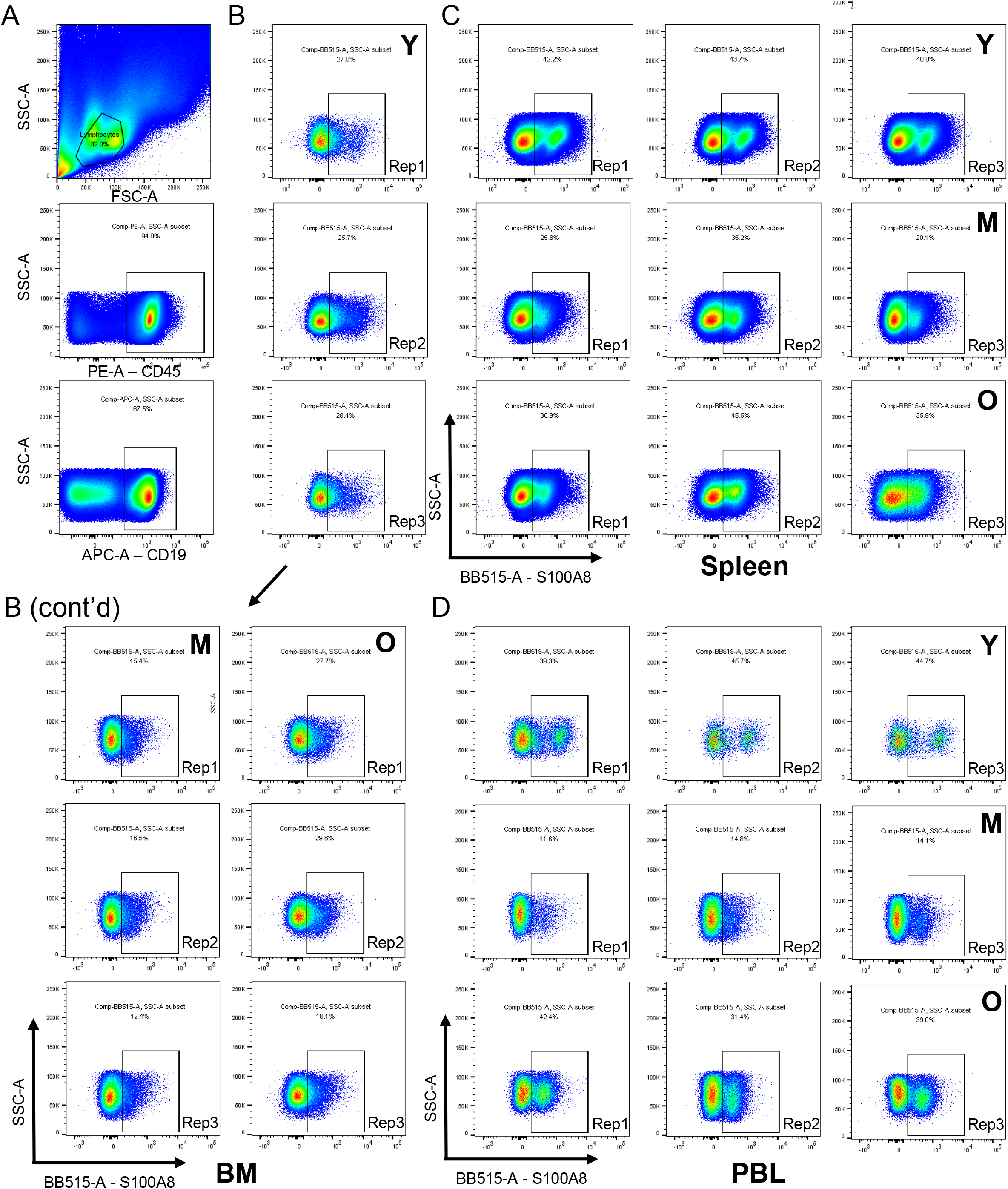
Age-associated proportion of S100A8+ B cells from C57BL/6J mice as determined by FACS. **(A)** B cell ing strategy (CD45+CD19+) as shown by a spleen sample from a young mouse. **(B-D)** Gating of S100A8+ B cells from BM (B), spleen (C) PBL (D) from mice of different ages (n=3 for each age group). Y, young (3m); M, middle age (12m); O, old (22m).

**Supplementary figure 23.**
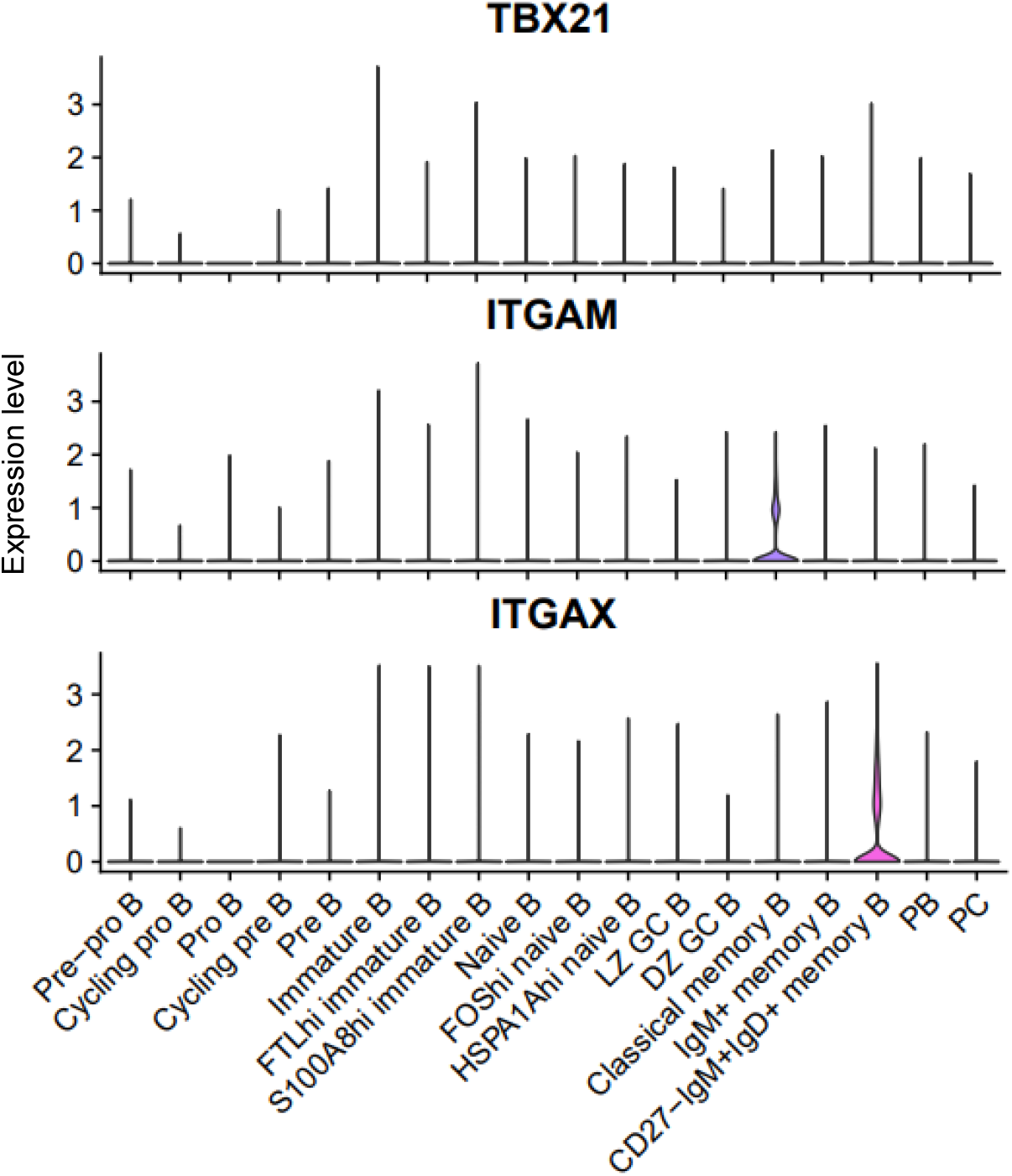
Expression level of the ABC marker genes, TBX21 (T-bet), ITGAM (CD11b), and ITGAX (CD11c) across B cell subpopulations as shown with Seurat VlnPlot methodology.

**Table S1.** Composition of the integrated dataset

| Index | Source | Cell type | Tissue | # Sample | # Donor | Status | # Total cells | # After QC |
| --- | --- | --- | --- | --- | --- | --- | --- | --- |
| 1 | This study | Enriched B cells | BM | 9 | 3 | FFHD | 114,058 | 89,762 (78.7%) |
| 2 | This study | Enriched B cells | PBL | 3 | 3 | FFHD | 15,661 | 13,522 (86.3%) |
| 3 | This study | BMMC | BM | 6 | 3 | FFHD | 34,796 | 26,548 (76.3%) |
| 4 | Zhang et al. | CD34+CD19- cells/CD34-CD19+ cells | BM | 4 | 2 | Healthy | 47,471 | 28,305 (59.6%) |
| 5 | Zhu et al. | PBMC | PBL | 3 | 3 | Healthy | 10,718 | 10,407 (97.1%) |
| 6 | King et al | Unsorted tonsillar immune cells | Tonsils | 7 | 7 | RT/OSA | 33,538 | 30,460 (90.8%) |
| **Total** |  | - | - | **32** | **15** | - | **256,242** | **199,004 (77.7%)** |

Note: BM, bone marrow; PBL, peripheral blood; PBMC, peripheral blood mononuclear cell; BMMC, bone marrow mononuclear cell; RT, recurrent tonsillitis; OSA, obstructive sleep apnea; FFHD, free from hematological diseases.

**Table S2.**
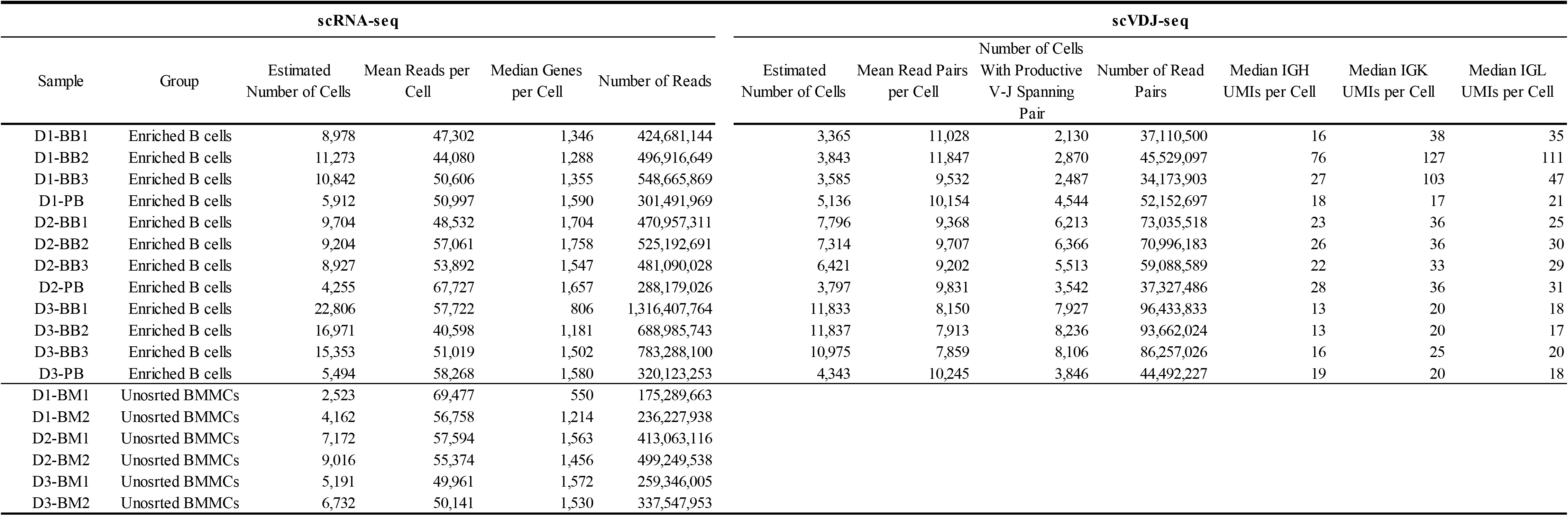
QC metrics for scRNA-seq and scVDJ-seq samples in this study

**Table S3.** Cell population/subpopulation frequency in the integrated data

| Index | Cell population/subpopulation | Frequency | Cell group |
| --- | --- | --- | --- |
| 1 | HSC | 1,251 | HSPC |
| 2 | CMP | 1,200 | HSPC |
| 3 | GMP | 266 | HSPC |
| 4 | MEP | 732 | HSPC |
| 5 | CLP | 385 | HSPC |
| 6 | Pre-pro B | 913 | B lineage |
| 7 | Cycling pro B | 523 | B lineage |
| 8 | Pro B | 476 | B lineage |
| 9 | Cycling pre B | 1,896 | B lineage |
| 10 | Pre B | 7,287 | B lineage |
| 11 | Immature B | 14,996 | B lineage |
| 12 | FTL<sup>hi</sup> immature B | 3,199 | B lineage |
| 13 | S100A8<sup>hi</sup> immature B | 3,457 | B lineage |
| 14 | Naive B | 50,181 | B lineage |
| 15 | FOS<sup>hi</sup> naive B | 5,307 | B lineage |
| 16 | HSPA1<sup>hi</sup> naive B | 1,683 | B lineage |
| 17 | LZ GC B | 6,597 | B lineage |
| 18 | DZ GC B | 4,902 | B lineage |
| 19 | Classical memory B | 3,698 | B lineage |
| 20 | IgM+ memory B | 20,465 | B lineage |
| 21 | CD27-IgM+IgD+ memory B | 3,125 | B lineage |
| 22 | PB | 2,669 | B lineage |
| 23 | PC | 3,225 | B lineage |
| 24 | CD4+ T | 14,689 | T/NK lineage |
| 25 | CD8+ T | 8,814 | T/NK lineage |
| 26 | MAIT | 1,213 | T/NK lineage |
| 27 | gd T | 996 | T/NK lineage |
| 28 | Treg | 1,027 | T/NK lineage |
| 29 | NK | 6,117 | T/NK lineage |
| 30 | CD14+ monocyte | 5,349 | Myeloid |
| 31 | CD16+ monocyte | 500 | Myeloid |
| 32 | Macrophage | 2,941 | Myeloid |
| 33 | cDC | 470 | Myeloid |
| 34 | pDC | 1,114 | Myeloid |
| 35 | Pro-myelocyte | 729 | Myeloid |
| 36 | Myelocyte | 483 | Myeloid |
| 37 | Myeloblast | 881 | Myeloid |
| 38 | Neutrophil | 1,120 | Myeloid |
| 39 | Erythroblast | 4,845 | Erythroid/Meg |
| 40 | Erythrocyte | 5,669 | Erythroid/Meg |
| 41 | Megakaryocyte | 236 | Erythroid/Meg |
| 42 | MSC | 3,378 | Non-hematopoietic |

**Table S4.**
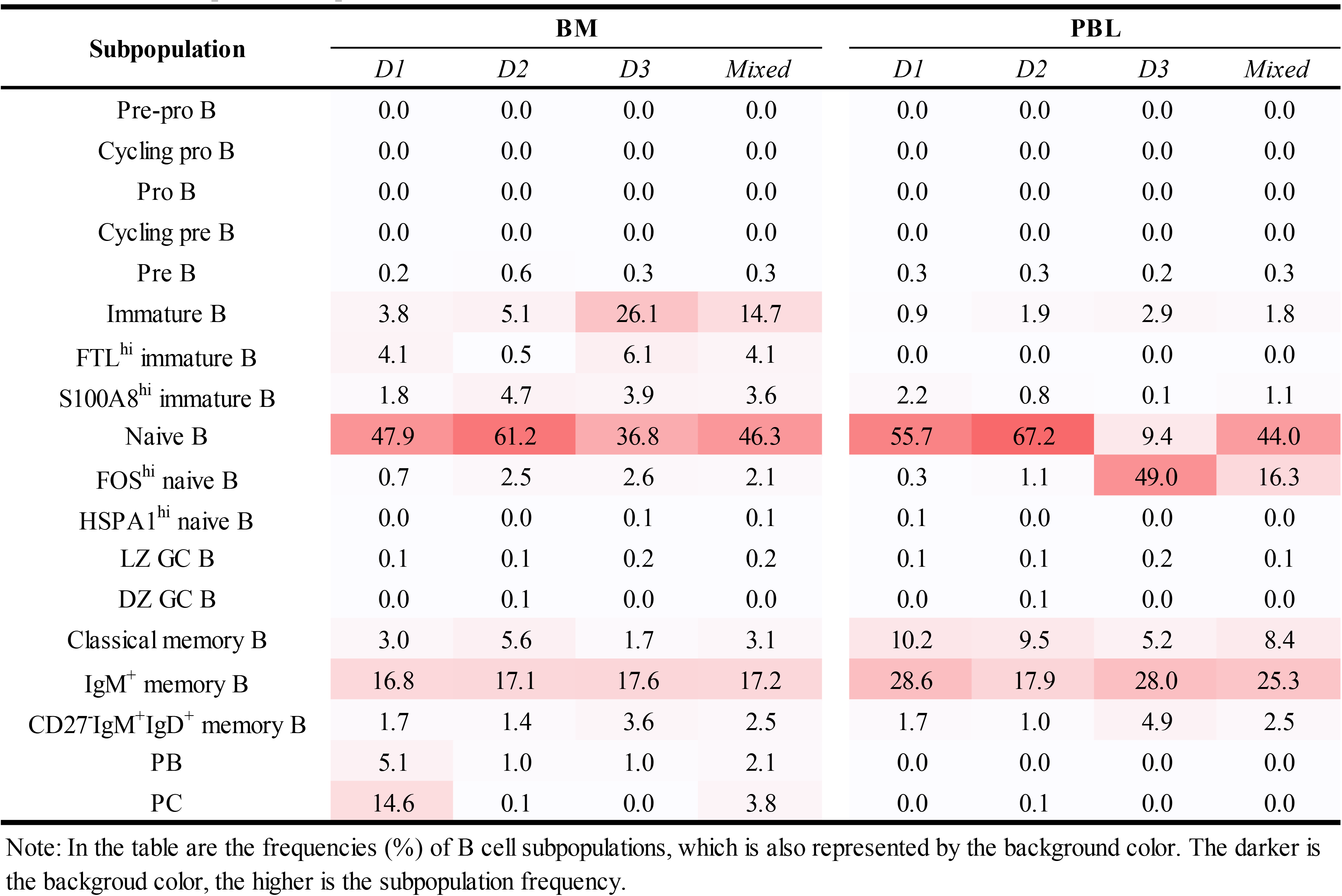
Tissue-specific composition of B cells across donors and as a whole

**Table S5.** The B cell subpopulation composition in enriched B cell samples (BM, PBL and BM & PBL)

|  | BM |  | PBL |  | BM & PBL |  | scVDJ-seq |  | Clonotypes |
| --- | --- | --- | --- | --- | --- | --- | --- | --- | --- |
|  | # | % | # | % | # | % | # | % | # |
| Cycling pro B | 5 | 0.01 | - | 0.00 | 5 | 0.01 | - | - | - |
| Pro B | 3 | 0.00 | - | 0.00 | 3 | 0.00 | - | - | - |
| Cycling pre B | 10 | 0.01 | - | 0.00 | 10 | 0.01 | 3 | 30.00 | 3 |
| Pre B | 266 | 0.35 | 33 | 0.26 | 299 | 0.33 | 222 | 74.25 | 218 |
| Immature B | 11,272 | 14.67 | 230 | 1.83 | 11,502 | 12.87 | 1,849 | 16.08 | 1,635 |
| FTLhi immature B | 3,113 | 4.05 | 3 | 0.02 | 3,116 | 3.49 | 1,451 | 46.57 | 1,419 |
| S100A8hi immature B | 2,766 | 3.60 | 139 | 1.11 | 2,905 | 3.25 | 463 | 15.94 | 452 |
| Naive B | 35,550 | 46.28 | 5,525 | 43.99 | 41,075 | 45.95 | 26,472 | 64.45 | 26,110 |
| FOShi naive B | 1,611 | 2.10 | 2,047 | 16.30 | 3,658 | 4.09 | 2,930 | 80.10 | 2,865 |
| HSPA1Ahi naive B | 45 | 0.06 | 5 | 0.04 | 50 | 0.06 | 14 | 28.00 | 14 |
| LZ GC B | 118 | 0.15 | 17 | 0.14 | 135 | 0.15 | 74 | 54.81 | 72 |
| DZ GC B | 24 | 0.03 | 3 | 0.02 | 27 | 0.03 | 6 | 22.22 | 6 |
| Classical memory B | 2,395 | 3.12 | 1,054 | 8.39 | 3,449 | 3.86 | 1,568 | 45.46 | 1,537 |
| IgM+ memory B | 13,228 | 17.22 | 3,182 | 25.33 | 16,410 | 18.36 | 7,208 | 43.92 | 6,880 |
| CD27-IgM+IgD+ memory B | 1,926 | 2.51 | 318 | 2.53 | 2,244 | 2.51 | 1,140 | 50.80 | 925 |
| PB | 1,579 | 2.06 | 3 | 0.02 | 1,582 | 1.77 | 1,414 | 89.38 | 1,039 |
| PC | 2,911 | 3.79 | 2 | 0.02 | 2,913 | 3.26 | 2,522 | 86.58 | 1,913 |
| Total | 76,822 | 100.00 | 12,561 | 100.00 | 89,383 | 100.00 | 47,336 | 52.96 | 45,088 |

**Note:** B cell subpopulations highlighted in red in theory do not exist in BM and PBL. B cell subpopulations highlighted in gray are rare (<0.5%) in both BM and PBL. Therefore, only the rest subpopulations can be investigated with in-house scVDJ-seq data. whereas the transcriptomic-based analyses apply to all subpopulations.

## References

1. Lee, R.D. et al. Single-cell analysis identifies dynamic gene expression networks that govern B cell development and transformation. Nat Commun 12, 6843 (2021).

2. King, H.W., et al. Single-cell analysis of human B cell maturation predicts how antibody class switching shapes selection dynamics. Sci Immunol 6 (2021).

3. Morgan, D. & Tergaonkar, V. Unraveling B cell trajectories at single cell resolution. Trends Immunol 43, 210–229 (2022).

4. Glass, D.R. et al. An Integrated Multi-omic Single-Cell Atlas of Human B Cell Identity. Immunity 53, 217–232 e215 (2020).

5. Stewart, A. et al. Single-Cell Transcriptomic Analyses Define Distinct Peripheral B Cell Subsets and Discrete Development Pathways. Front Immunol 12, 602539 (2021).

6. Brioschi, S. et al. Heterogeneity of meningeal B cells reveals a lymphopoietic niche at the CNS borders. Science 373 (2021).

7. Zhang, Y. et al. Elucidating minimal residual disease of paediatric B-cell acute lymphoblastic leukaemia by single-cell analysis. Nat Cell Biol 24, 242–252 (2022).

8. Zhu, L. et al. Single-Cell Sequencing of Peripheral Mononuclear Cells Reveals Distinct Immune Response Landscapes of COVID-19 and Influenza Patients. Immunity 53, 685–696 e683 (2020).

9. Wang, Q. et al. Dual UMIs and Dual Barcodes With Minimal PCR Amplification Removes Artifacts and Acquires Accurate Antibody Repertoire. Front Immunol 12, 778298 (2021).

10. Oetjen, K.A., et al. Human bone marrow assessment by single-cell RNA sequencing, mass cytometry, and flow cytometry. JCI Insight 3 (2018).

11. Carrion, C. et al. Adult Bone Marrow Three-Dimensional Phenotypic Landscape of B-Cell Differentiation. Cytometry B Clin Cytom 96, 30–38 (2019).

12. Perez-Andres, M. et al. Human peripheral blood B-cell compartments: a crossroad in B-cell traffic. Cytometry B Clin Cytom 78 **Suppl 1**, S47–60 (2010).

13. Hardy, R.R. & Hayakawa, K. B cell development pathways. Annu Rev Immunol 19, 595–621 (2001).

14. Zeng, H. et al. Discrete roles and bifurcation of PTEN signaling and mTORC1-mediated anabolic metabolism underlie IL-7-driven B lymphopoiesis. Sci Adv 4, eaar5701 (2018).

15. LeBien, T.W. & Tedder, T.F. B lymphocytes: how they develop and function. Blood 112, 1570–1580 (2008).

16. Sontheimer, R.D., Racila, E. & Racila, D.M. C1q: its functions within the innate and adaptive immune responses and its role in lupus autoimmunity. J Invest Dermatol 125, 14–23 (2005).

17. Wang, S. et al. S100A8/A9 in Inflammation. Front Immunol 9, 1298 (2018).

18. Bandyopadhyay, S. et al. Mapping the cellular biogeography of human bone marrow niches using single-cell transcriptomics and proteomic imaging. Cell 187, 3120–3140 e3129 (2024).

19. Caron, M. et al. Single-cell analysis of childhood leukemia reveals a link between developmental states and ribosomal protein expression as a source of intra-individual heterogeneity. Sci Rep 10, 8079 (2020).

20. Chen, M. et al. Dynamic single-cell RNA-seq analysis reveals distinct tumor program associated with microenvironmental remodeling and drug sensitivity in multiple myeloma. Cell Biosci 13, 19 (2023).

21. de Jong, M.M.E. et al. The multiple myeloma microenvironment is defined by an inflammatory stromal cell landscape. Nat Immunol 22, 769–780 (2021).

22. Lee, N.Y.S., Li, M., Ang, K.S. & Chen, J. Establishing a human bone marrow single cell reference atlas to study ageing and diseases. Front Immunol 14, 1127879 (2023).

23. Li, X. et al. Inflammation and aging: signaling pathways and intervention therapies. Signal Transduct Target Ther 8, 239 (2023).

24. Naito, A.T. et al. Complement C1q activates canonical Wnt signaling and promotes aging- related phenotypes. Cell 149, 1298–1313 (2012).

25. Albakova, Z., Mangasarova, Y. & Sapozhnikov, A. Impaired Heat Shock Protein Expression in Activated T Cells in B-Cell Lymphoma. Biomedicines 10 (2022).

26. Woodland, R.T. & Schmidt, M.R. Homeostatic proliferation of B cells. Semin Immunol 17, 209–217 (2005).

27. Harms Pritchard, G. & Pepper, M. Memory B cell heterogeneity: Remembrance of things past. J Leukoc Biol 103, 269–274 (2018).

28. Berkowska, M.A. et al. Human memory B cells originate from three distinct germinal center- dependent and -independent maturation pathways. Blood 118, 2150–2158 (2011).

29. MacLennan, I.C. Germinal centers. Annu Rev Immunol 12, 117–139 (1994).

30. Mond, J.J., Vos, Q., Lees, A. & Snapper, C.M. T cell independent antigens. Curr Opin Immunol 7, 349–354 (1995).

31. Weill, J.C., Weller, S. & Reynaud, C.A. Human marginal zone B cells. Annu Rev Immunol 27, 267–285 (2009).

32. Clavarino, G. et al. Novel Strategy for Phenotypic Characterization of Human B Lymphocytes from Precursors to Effector Cells by Flow Cytometry. PLoS One 11, e0162209 (2016).

33. Wu, Y.C., Kipling, D. & Dunn-Walters, D.K. The relationship between CD27 negative and positive B cell populations in human peripheral blood. Front Immunol 2, 81 (2011).

34. Wu, Y.C. et al. High-throughput immunoglobulin repertoire analysis distinguishes between human IgM memory and switched memory B-cell populations. Blood 116, 1070–1078 (2010).

35. Nagasawa, T. Microenvironmental niches in the bone marrow required for B-cell development. Nat Rev Immunol 6, 107–116 (2006).

36. De Silva, N.S. & Klein, U. Dynamics of B cells in germinal centres. Nat Rev Immunol 15, 137–148 (2015).

37. Wang, X. & Lin, Y. Tumor necrosis factor and cancer, buddies or foes? Acta Pharmacol Sin 29, 1275–1288 (2008).

38. Ryan, D.H., Nuccie, B.L., Abboud, C.N. & Winslow, J.M. Vascular cell adhesion molecule-1 and the integrin VLA-4 mediate adhesion of human B cell precursors to cultured bone marrow adherent cells. J Clin Invest 88, 995–1004 (1991).

39. Takeda, S., Shimizu, T. & Rodewald, H.R. Interactions between c-kit and stem cell factor are not required for B-cell development in vivo. Blood 89, 518–525 (1997).

40. Waskow, C., Paul, S., Haller, C., Gassmann, M. & Rodewald, H.R. Viable c-Kit(W/W) mutants reveal pivotal role for c-kit in the maintenance of lymphopoiesis. Immunity 17, 277–288 (2002).

41. Cheng, P. et al. Inhibition of dendritic cell differentiation and accumulation of myeloid-derived suppressor cells in cancer is regulated by S100A9 protein. J Exp Med 205, 2235–2249 (2008).

42. Goyette, J. & Geczy, C.L. Inflammation-associated S100 proteins: new mechanisms that regulate function. Amino Acids 41, 821–842 (2011).

43. Roth, J., Vogl, T., Sorg, C. & Sunderkotter, C. Phagocyte-specific S100 proteins: a novel group of proinflammatory molecules. Trends Immunol 24, 155–158 (2003).

44. Sinha, P. et al. Proinflammatory S100 proteins regulate the accumulation of myeloid-derived suppressor cells. J Immunol 181, 4666–4675 (2008).

45. Kitagori, K. et al. Expression of S100A8 protein on B cells is associated with disease activity in patients with systemic lupus erythematosus. Arthritis Res Ther 25, 76 (2023).

46. Lee, J., Kim, H., Kim, M., Yoon, S. & Lee, S. Role of lymphoid lineage cells aberrantly expressing alarmins S100A8/A9 in determining the severity of COVID-19. Genes Genomics 45, 337–346 (2023).

47. Zhang, B. et al. SenoIndex: S100A8/S100A9 as a novel aging biomarker. Life Med 2 (2023).

48. Tabula Muris, C. A single-cell transcriptomic atlas characterizes ageing tissues in the mouse. Nature 583, 590–595 (2020).

49. de Mol, J., Kuiper, J., Tsiantoulas, D. & Foks, A.C. The Dynamics of B Cell Aging in Health and Disease. Front Immunol 12, 733566 (2021).

50. Yu, Y. et al. B Cells Dynamic in Aging and the Implications of Nutritional Regulation. Nutrients 16 (2024).

51. Knox, J.J., Myles, A. & Cancro, M.P. T-bet(+) memory B cells: Generation, function, and fate. Immunol Rev 288, 149–160 (2019).

52. Burrows, N. et al. Dynamic regulation of hypoxia-inducible factor-1alpha activity is essential for normal B cell development. Nat Immunol 21, 1408–1420 (2020).

53. Mehtonen, J. et al. Single cell characterization of B-lymphoid differentiation and leukemic cell states during chemotherapy in ETV6-RUNX1-positive pediatric leukemia identifies drug- targetable transcription factor activities. Genome Med 12, 99 (2020).

54. van Zelm, M.C., Szczepanski, T., van der Burg, M. & van Dongen, J.J. Replication history of B lymphocytes reveals homeostatic proliferation and extensive antigen-induced B cell expansion. J Exp Med 204, 645–655 (2007).

55. Agenes, F. & Freitas, A.A. Transfer of small resting B cells into immunodeficient hosts results in the selection of a self-renewing activated B cell population. J Exp Med 189, 319–330 (1999).

56. Cabatingan, M.S., Schmidt, M.R., Sen, R. & Woodland, R.T. Naive B lymphocytes undergo homeostatic proliferation in response to B cell deficit. J Immunol 169, 6795–6805 (2002).

57. Yang, Y. et al. Pan-cancer single-cell dissection reveals phenotypically distinct B cell subtypes. Cell 187, 4790–4811 e4722 (2024).

58. Seifert, M. & Kuppers, R. Human memory B cells. Leukemia 30, 2283–2292 (2016).

59. LeBien, T.W. B Cell Development. (Elsevier, 2017).

60. Victora, G.D. & Nussenzweig, M.C. Germinal Centers. Annu Rev Immunol 40, 413–442 (2022).

61. Bae, S., et al. Existence of blood circulating immune-cell clusters (CICs) comprising antigen- presenting cells and B cells. bioRxiv (2022).

62. Cravedi, P. et al. Increased Complement Gene Expression in Circulating B Cells From Kidney Transplant Recipients With Chronic Antibody-Mediated Rejection. Kidney Int Rep 7, 2752–2753 (2022).

63. Yang, X. et al. Novel Allele Detection Tool Benchmark and Application With Antibody Repertoire Sequencing Dataset. Front Immunol 12, 739179 (2021).

64. Zhu, Y. et al. Antibody upstream sequence diversity and its biological implications revealed by repertoire sequencing. J Genet Genomics 48, 936–945 (2021).

65. Stephenson, E. et al. Single-cell multi-omics analysis of the immune response in COVID-19. Nat Med 27, 904–916 (2021).

66. La Manno, G. et al. RNA velocity of single cells. Nature 560, 494–498 (2018).

67. Bergen, V., Lange, M., Peidli, S., Wolf, F.A. & Theis, F.J. Generalizing RNA velocity to transient cell states through dynamical modeling. Nat Biotechnol 38, 1408–1414 (2020).

68. Wolf, F.A., Angerer, P. & Theis, F.J. SCANPY: large-scale single-cell gene expression data analysis. Genome Biol 19, 15 (2018).

69. Qiu, X. et al. Reversed graph embedding resolves complex single-cell trajectories. Nat Methods 14, 979–982 (2017).

70. Kuleshov, M.V. et al. Enrichr: a comprehensive gene set enrichment analysis web server 2016 update. Nucleic Acids Res 44, W90–97 (2016).

71. Van de Sande, B. et al. A scalable SCENIC workflow for single-cell gene regulatory network analysis. Nat Protoc 15, 2247–2276 (2020).

72. Troule, K. et al. CellPhoneDB v5: inferring cell-cell communication from single-cell multiomics data. arXiv (2023).

