## Supplementary figures and images for "Single-cell analysis reveals multi-faceted features of B cell development together with age-associated B cell subpopulations"

### Supp. Data 1

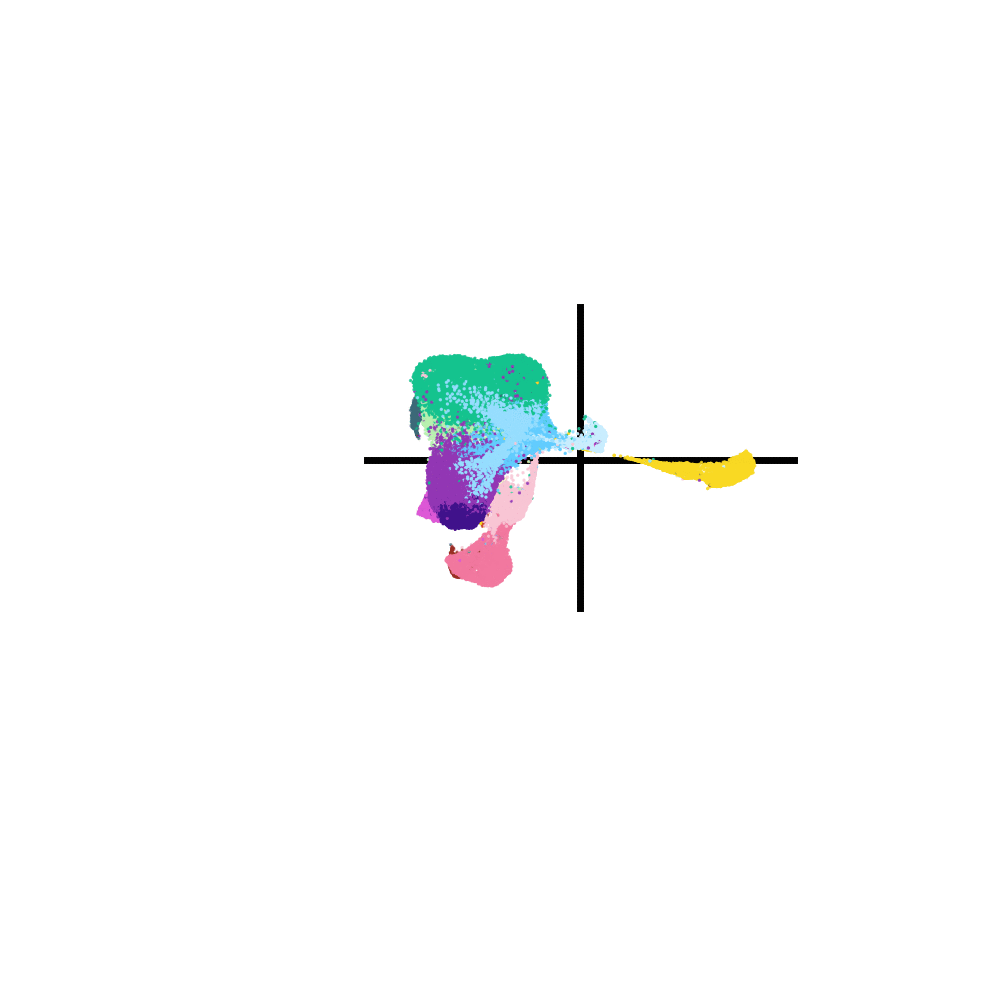
